# Genetic diversity and population structure analyses in the Alpine plum (*Prunus brigantina* Vill.) confirm its affiliation to the Armeniaca section

**DOI:** 10.1101/2020.08.12.247718

**Authors:** Liu Shuo, Decroocq Stephane, Harte Elodie, Tricon David, Chague Aurelie, Balakishiyeva Gulnara, Kostritsyna Tatiana, Turdiev Timur, Fisher-Le Saux Marion, Dallot Sylvie, Giraud Tatiana, Decroocq Veronique

## Abstract

In-depth characterization of the genetic diversity and population structure of wild relatives of crops is of paramount importance for genetic improvement and biodiversity conservation, and is particularly crucial when the wild relatives of crops are endangered. In this study, we therefore sampled the Alpine plum (Briançon apricot) *Prunus brigantina* Vill. across its natural distribution in the French Alps, where its populations are severely fragmented and its population size strongly impacted by humans. We analysed 71 wild *P. brigantina* samples with 34 nuclear markers and studied their genetic diversity and population structure, with the aim to inform *in situ* conservation measures and build a core collection for long-term ex-situ conservation. We also examined the genetic relationships of *P. brigantina* with other species in the Prunophora subgenus, encompassing the Prunus (Eurasian plums), Prunocerasus (North-American plums) and Armeniaca (apricots) sections, to check its current taxonomy. We detected a moderate genetic diversity in *P. brigantina* and a Bayesian model-based clustering approach revealed the existence of three genetically differentiated clusters, endemic to three geographical regions in the Alps, which will be important for *in situ* conservation measures. Based on genetic diversity and population structure analyses, a subset of 36 accessions were selected for *ex-situ* conservation in a core collection that encompasses the whole detected *P. brigantina* allelic diversity. Using a dataset of cultivated apricots and wild cherry plums (*P. cerasifera*) genotyped with the same markers, we detected gene flow neither with European *P. armeniaca* cultivars nor with diploid plums. In contrast with previous studies, dendrograms and networks placed *P. brigantina* closer to Armeniaca species than to Prunus species. Our results thus confirm the classification of *P. brigantina* within the Armeniaca section; it also illustrates the importance of the sampling size and design in phylogenetic studies.

## Introduction

Many wild crop relatives are endangered, because of fragmented and reduced habitats as well as crop-to-wild gene flow (Cornille et al. 2013b). In order to protect the biodiversity of wild crop relatives, we need to understand their population subdivision and genetic diversity distribution (Allendorf et al. 2012; Fahrig 2003). Studying the genetic diversity of crop-related species is not only important for biodiversity conservation but also for the sustainable use of valuable genetic resources through the set-up of *ex-situ* germplasm collections (Li and Pritchard 2009). Developing such collections requires obtaining a sufficient number of individuals to be representative of the species diversity (Frankel and Brown 1983; Glaszmann et al. 2010; Govindaraj et al. 2014). Core collections of woody perennial species have the additional advantages of being propagated vegetatively and maintained for decades, as clonemates, in field collections (Escribano et al. 2008).

Within the genus *Prunus* L. (stone fruit species), the subgenus Prunus (also called Prunophora Neck. to avoid confusion with the Prunus section and the *Prunus* genus) includes three sections: the Eurasian and North American plums (sections Prunus and Prunocerasus Koehne, respectively) and apricots (section Armeniaca (Mill.) K. Koch), which are all native from the Northern hemisphere (Rehder 1940) (Figure 1). It was shown that an ancient radiation of the *Prunus* genus through the Old and New Worlds and independent dispersal events across the North-American and Eurasian continents gave rise to, on one side, the Prunocerasus species, and on the other side, species of the Prunus and Armeniaca sections (Chin et al. 2014). Following the Rehder’s classification, within the Prunophora subgenus the section Armeniaca comprises only diploid species, in which six species are recognized, based on morphological features: *P. armeniaca* L. (common apricot), *P. sibirica* L. (wild apricot in Northeastern Asia), *P. mandshurica* Maxim. (Northeast China and Eastern Russia), *P. mume* (Sieb.) Sieb. & Zucc. (South China and Japan), *P. holosericeae* Batal (South-West China) and *P. brigantina* Vill. (Figure 1). The first five species all originate from Asia, ranging from Central to North-East Asia, while *P. brigantina* (synonym *P. brigantiaca*, http://www.theplantlist.org) is native from Europe (Villars 1786). This species still grows in wild, patchy thickets in the Alps along the border between France and Italy in the Northern Mediterranean area, where it is considered either as an apricot or a plum species (Hagen et al. 2002; Pignatti 1982).

**Figure 1.**
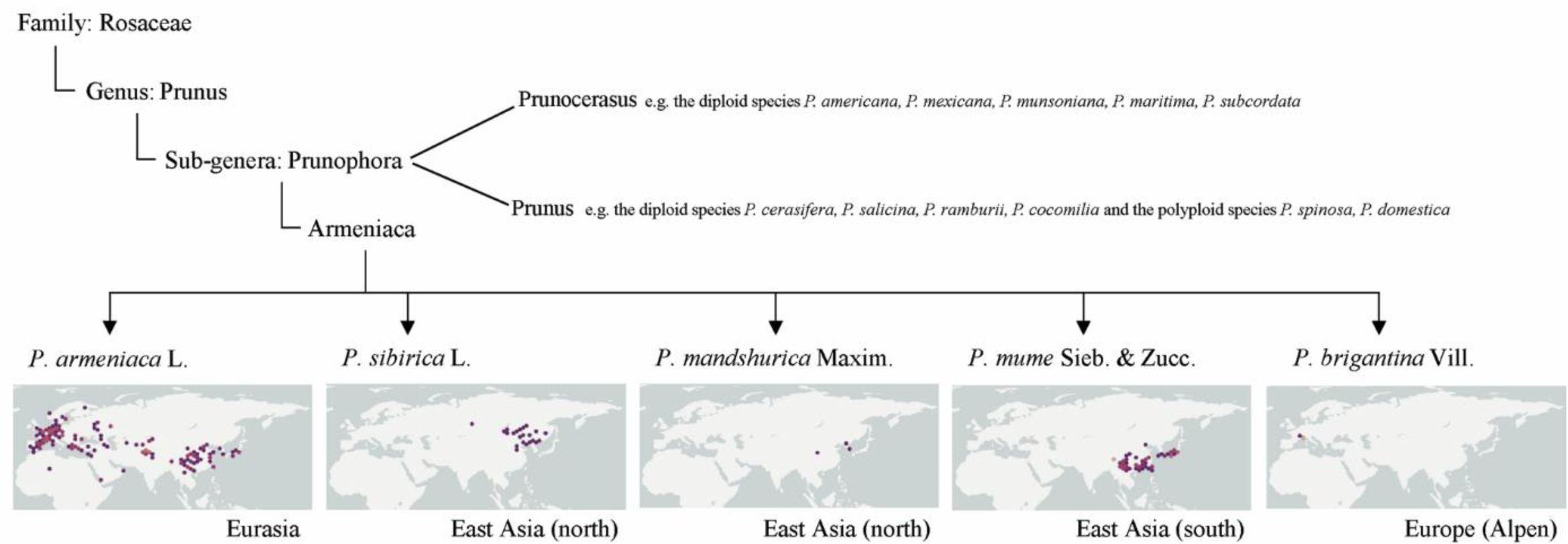
Taxonomy and geographic distribution of the different species in the Armeniaca section. Species classification is based on reports by Rehder (1940). Data on species distribution were retrieved from the global biodiversity information facility (GBIF) (https://doi.org/10.15468/39omei). Dots represent georeferenced species records from 1910 to 2017.

*Prunus brigantina*, alternatively called the Briançon apricot or the Alpine plum, was first reported in the French book <*Histoire des Plants de Dauphiné*> (Villars 1786). It grows in arid places in shrub and sparse thickets in the Alps, above 1,400 m altitude. Like other *Prunus* species, *P. brigantina* is hermaphrodite and is pollinated by insects, it flowers in May and its fruits ripen from August to September (Noble et al. 2015; Tison and De Foucault 2014). In natural stands, *P. brigantina* trees grow 2 to 5 meters high with non-spiny branches and have heart leaves with double-serrated teeth (Figure 2a). The full-fledged drupe from *P. brigantina* has a small size and appears glabrous with yellowish fruit skin (Figure 2a). In the Alps, *P. brigantina* fruits are collected by locals to make jam (Couplan 2009), and their seeds used to be processed for oil production instead of olive or almond (Dupouy 1959). It is locally called ‘Marmottier’ or ‘Afatoulier’ and is recognized as an endemic fruit tree in Europe. Its small and fragmented distribution suggests that it may be threatened. However, there is currently insufficient information available to evaluate the current genetic diversity of *P. brigantina* or its population subdivision, which could contribute to determine the potential threats to this species or its conservation status (IUCN Red List) (Branca and Donnini 2011), and to develop and inform conservation programs.

**Figure 2.**
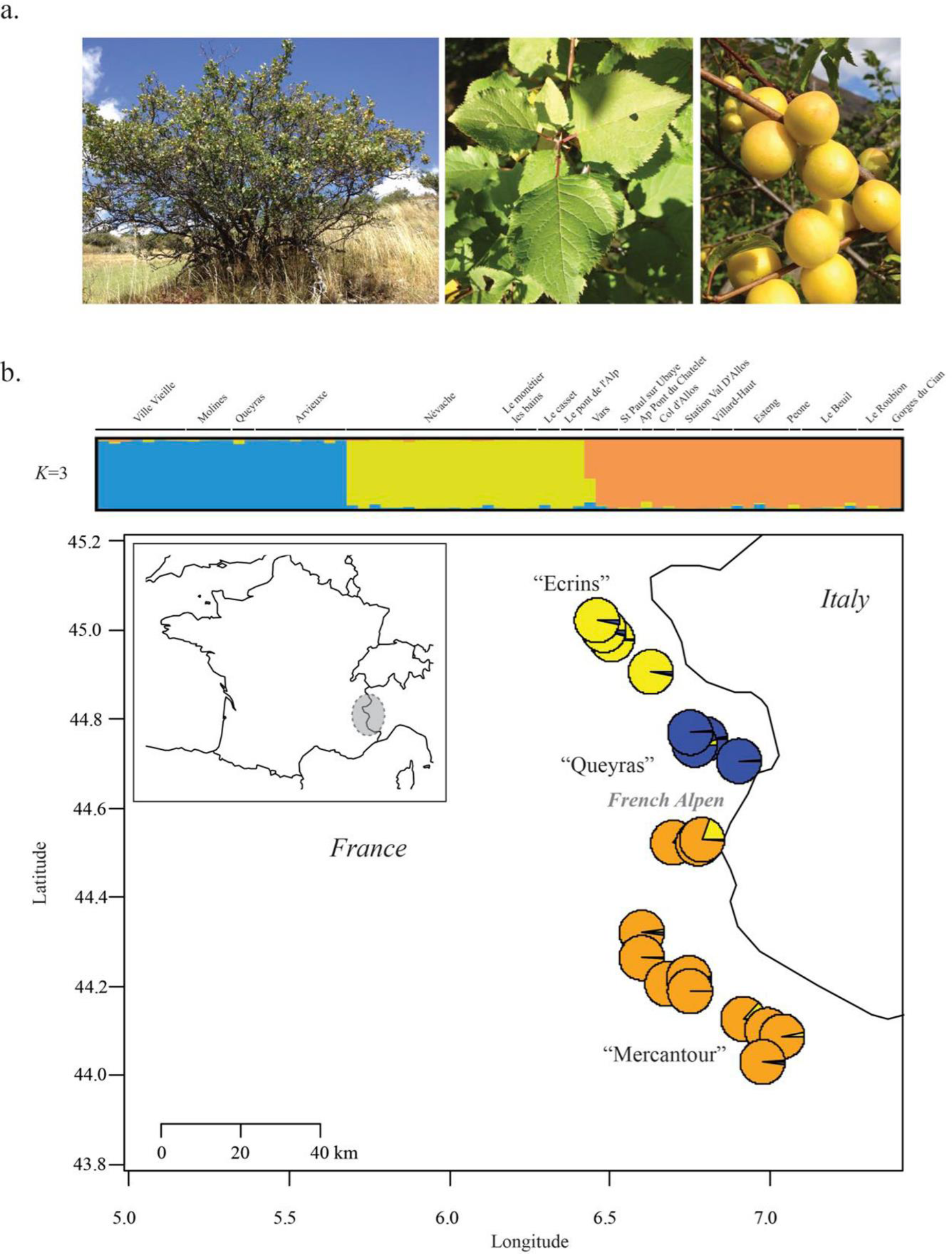
*Prunus brigantina* morphological features, genetic clustering and spatial distribution in the French Alps. a. A *P. brigantina* small tree in its natural habitat (Arvieux) (left), summery leaves (middle) and ripening fruits (right). b. The three genetic clusters of *P. brigantina* inferred from the STRUCTURE analysis (Figure S2 at *K*=3) and their spatial distribution in the French Alps. “Ecrins”, “Queyras” and “Mercantour” refer to the three national parks in the southeast of France.

Previous phylogenetic studies questioned the Rehder’s classification of *P. brigantina* in the Armeniaca section (Hagen et al. 2002; Reales et al. 2010; Takeda et al. 1998; Zhebentyayeva et al. 2019). However, only one or two *P. brigantina* samples were analysed, that had not been collected *in situ* but instead obtained from germplasm repositories such as the Kew Royal Botanical Garden (UK), the Czech national genetic resources of Lednice, the French Centre of genetic resources at INRAE-Montfavet or the Japanese Chiyoda experimental station. Because of the ability of *P. brigantina* to be propagated by grafting and its interfertility with species from both sections Prunus and Armeniaca, the analyzed trees could be clonemates or hybrids, and their origin was unknown. Moreover, sampling only one or two individuals per species is known to lower the accuracy of phylogenetic analyses (Heled and Drummond 2010; Wiens and Servedio 1997). The genetic relationships between *P. brigantina* and other apricot species in the section Armeniaca therefore remain unclear.

To provide useful guidelines for *P. brigantina* conservation, a critical first task is first assessing its genetic diversity distribution and population subdivision, in order to assess whether local specificities need to be conserved. A second important aspect is to clarify the taxonomic position of the different populations of *P. brigantina*. Indeed, differentiated populations assigned to a given Latin species may actually represent different species placed far apart in phylogeny, as found recently for *P. sibirica* (Liu et al. 2019). However, there is currently no robust data on the genetic diversity or population structure of *P. brigantina*. In the current study, we therefore conducted extensive sampling of *P. brigantina* in its natural habitat and genotyped samples using 34 nuclear markers (Liu et al. 2019). We assessed the genetic diversity of *P. brigantina* and its population structure, as well as its relationship with Eurasian Armeniaca and Prunus species. We questioned the affiliation of *P. brigantina* to either the Armeniaca or Prunus sections. For this purpose, we performed a population genetic analysis using datasets with the main members of the Armeniaca and Prunus sections and five outgroup species from the North-American section Prunocerasus. Based on our molecular data, we identified a collection of unique genotypes and selected the best subset for building a *P. brigantina* core collection, maximizing allelic diversity, which will be very useful for further *P. brigantina* characterization and for stone fruit crop improvement. In contrast to previous phylogenetic studies, our population trees and networks further confirmed that *P. brigantina* is closer to apricot species than to plum species.

## Materials and Methods

### *In situ P. brigantina* sampling

A total of 71 wild *P. brigantina* trees were collected in 2017 from three sampling sites, in southeast France, across the Alps (Figure 2a and Table S1). Young leaves and mature fruits from each tree were collected for DNA extraction and seedling growth, respectively. At least one seedling from each sampled tree was kept for possible inclusion in a core collection.

Representatives of other species of the Armeniaca section (*P. armeniaca, P. mume, P. sibirica, P. mandshurica*), including two *P. brigantina* accessions maintained by the Centre of genetic resources at INRA-Montfavet but with unknown origin, were previously described and genotyped with the same set of molecular markers (Liu et al. 2019).

### *In situ* and *ex situ* sampling of representatives of Prunus and Prunocerasus species

Part of the plum and plum-related material analysed in this study was kindly provided by the North-American national repository (ARS-USDA, Davis, California, USA), the Bourran’s collection of Prunus (Prunus Genetic Resources Centre or Prunus GRC, France) or was collected *in situ*, between 2008 and 2019, in Azerbaijan (Caucasia), Kazakhstan and Kyrgyzstan (Central Asia) (Table S1b). One *P. cerasifera* sample (X29) was collected *in situ* in South-West of France, along the Garonne river (Le Tourne-Langoiran) and another one in the French Alps (FR_070) (Table S1b). In total, 82 diploid samples were genotyped in the current study, genotypes for polyploids being difficult to analyse. The diploid samples included representatives from *P. cerasifera* (*N*=66) (or ‘cherry plum’, alternatively called ‘myrobolan’ in Europe, *P. divaricata* Ledeb. in Caucasia and *P. sogdiana* in Central Asia) and other species from the *Prunophora* subgenus: *P. mexicana* (*N*=1), *P. munsoniana* (*N*=1, also called *Prunus rivularis*), *P. maritima* (*N*=1), *P. americana* (*N*=1) and *P. subcordata* (*N*=1). *P. salicina* (Japanese plum) samples (*N*=10) were composed of five cultivated accessions which included one plumcot, a hybrid between *P. salicina* and *P. armeniaca* (called ‘Rutland’ in the ARS-USDA database, P0489) and five wild *P. salicina* accessions, sampled in China (Table S1b). *Prunus cerasifera* accessions used in this study originated from Europe (*N*=13), from Caucasia and Russia (*N*=29) and from Central Asia (more precisely from Kazakhstan and Kyrgyzstan, *N*=24) (see Table S1b for details).

### DNA extraction, microsatellite markers and polymerase chain reaction (PCR) amplification

Genomic DNA was extracted as described previously (Decroocq et al. 2016), either from lyophilized leaves, bark or fresh flowers. We used 34 microsatellite markers distributed across the eight *P. armeniaca* chromosomes and showing good amplification success as well as substantial polymorphism within the different species of the section *Armeniaca* (Liu et al. 2019). The same set of microsatellite markers were used to amplify PCR fragments in species of the Armeniaca (*P. brigantina* incl.), Prunus (*P. cerasifera* incl., see supplemental information) and Prunocerasus sections. Detailed information on these microsatellite markers, including their repeat motifs, sequences, and amplification conditions are available in (Liu et al. 2019). PCR amplification and fragment size genotyping were performed on an ABI PRISM 3730 (Applied Biosystems) as described previously (Decroocq et al. 2016). Alleles were scored with the GENEMAPPER 4.0 software (Applied Biosystems).

### Analyses of population subdivision and genetic relationships

To assess the probability of observing unrelated individuals with the detected similar genotypes given the population allelic frequencies, we used GENODIVE and the corrected *Nei’s* diversity estimate with a threshold of 50 (Meirmans and Van Tienderen 2004). We later retained only one individual of each pair detected as clonemates or siblings for further analyses.

We identified population subdivision with the STRUCTURE software v. 2.3.3 (Pritchard et al. 2000), without the use of *a priori* grouping information and assuming that individuals had mixed ancestry with correlated allele frequencies among populations. The clustering method implemented in STRUCTURE is based on Monte Carlo Markov Chain (MCMC) simulations and is used to infer the proportion of ancestry of genotypes in *K* distinct clusters. We simulated *K* values ranging from 2 to 10 for the *P. brigantina* population and three additional datasets (Table 1 and S1c), obtained with the same genetic markers on Armeniaca and Prunus species originating from Central and Eastern Asia (Liu et al. 2019) (Supplemental information). For each *K*, we ran 10,000 generations of ‘burn-in’ and 100,000 MCMC. Simulations were repeated 10 times for each *K* value; the resulting matrices of estimated cluster membership coefficients (Q) were permuted with CLUMPP (Jakobsson and Rosenberg 2007). STRUCTURE barplots were displayed with DISTRUCT 1.1 (Rosenberg 2004). The strongest level of the genetic subdivision was determined using Δ*K* (Evanno et al. 2005), as implemented in the online post-processing software Structure Harvester (http://taylor0.biology.ucla.edu/structureHarvester/) (Earl and vonHoldt 2012). Principal components analyses (PCA) were performed to investigate the genetic structure of *P. brigantina* using the scatterplot3d R package (Ligges and Mächler 2003) or among the five *Prunophora* species, using the DARwin software package v 6.0.017 (Perrier and Jacquemoud-Collet 2006). Further genetic differentiation and relationships were also estimated using a weighted neighbour-joining tree as implemented in the DARwin software package v 6.0.017 (Perrier and Jacquemoud-Collet 2006).

**Table 1.**
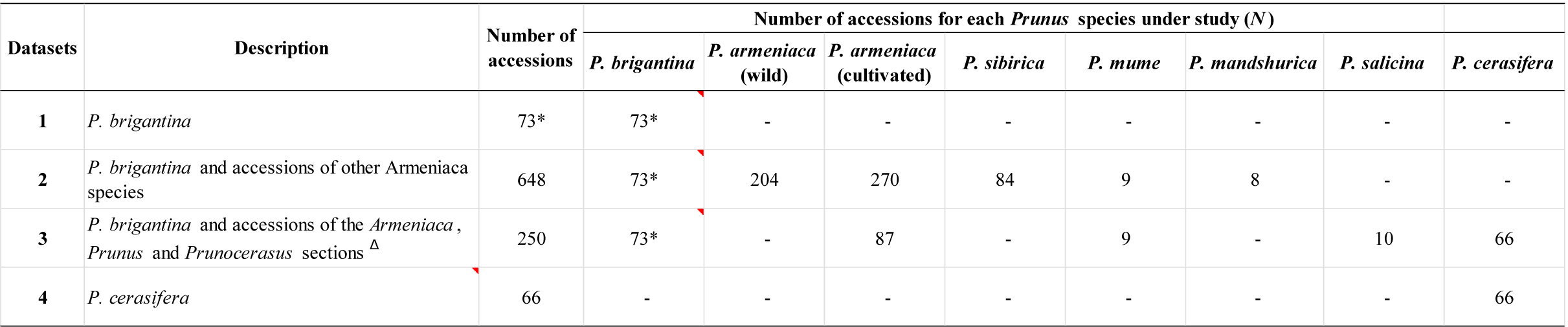
Different datasets including *Prunus brigantina* and other apricot species in this study. * indicate a *P. brigantina* dataset that includes 71 individuals sampled from the French Alps and 2 samples from the French GRC repository. ^Δ^ Prunocerasus species are represented by *P. mexicana* (*N*=1), *P. munsoniana* (*N*=1), *P. maritima* (*N*=1), *P. americana* (*N*=1), *P. subcordata* (*N*=1).

We performed a three-step population subdivision analysis, the first one with only *P. brigantina* samples, and the second one adding previously obtained datasets of *Armeniaca* species, including *P. armeniaca, P. sibirica, P. mandshurica* and *P. mume* wild and cultivated samples (Liu et al. 2019) (dataset 2 in Table 1). In the third step of the analysis, we added samples of the Prunus (*P. cerasifera* and *P. salicina*) and Prunocerasus (*P. mexicana, P. munsoniana, P. maritima, P. americana* and *P. subcordata)* sections (dataset 3 in Table 1). The same procedure to investigate population subdivision and structure analysis was used with the Armeniaca, Prunus and Prunocerasus diploid samples. In parallel, we also performed a population subdivision analysis along the native distribution of *P. cerasifera* (Supplemental information ‘*Prunus cerasifera* diversity and population structure analysis’ and dataset 4 in Table 1). In *P. cerasifera*, a Neighbour-Joining tree based on Nei’s standard genetic distance was built with a bootstrap of 30,000 in PopTreeW (Takezaki et al. 2014).

In order to test whether there was a pattern of isolation by distance, we performed a Mantel test between a matrix of Edwards’ genetic distances and a matrix of Euclidean geographic distances in *P. brigantina* using the R adegenet package (Jombart and Ahmed 2011).

### Genetic diversity, differentiation and core collection constitution

We used GENALEX 6.501 (Peakall and Smouse 2012) to estimate the number of alleles (*N*_a_), the effective number of alleles (*N*_e_), i.e., the number of equally frequent alleles that would achieve the same expected heterozygosity as in the sample, the observed heterozygosity (*H*_O_), the unbiased expected heterozygosity (*H*_E_) and the Shannon index (*I*) (Shannon 1948). Genetic differentiation among genetic clusters (Jost*’* D) was estimated in GENODIVE (Meirmans and Van Tienderen 2004). The allelic richness (*A*_*r*_) and the private allelic richness (*A*_*p*_) were calculated after adjustment for sample size differences among groups through the rarefaction procedure implemented in ADZE Allelic Diversity Analyzer v1.0 (Szpiech et al. 2008), setting the sample size to five.

The maximization (M) strategy (Schoen and Brown 1993) implemented in the COREFINDER software was used to generate a core *P. brigantina* tree collection maximizing the number of alleles based on our dataset. The maximization strategy consisted in detecting the smallest sample that captured 100% of the genetic diversity present within the entire germplasm collection. We further used the Mann-Whitney U test to check the genetic diversity difference between the core collection and the entire *P. brigantina* sample.

## Results

### Genetic diversity and population structuration in *P. brigantina*

Thirty-four microsatellite markers used in a previous study (Liu et al. 2019) were tested for our *P. brigantina* population study. Four markers (AMPA109, ssr02iso4G, BPPCT008 and BPPCT038) failed to amplify or generated over 50% of missing data and were consequently eliminated. Six other markers (BPPCT030, CPPCT022, CPSCT004, UDP98-412, UDA-002 and PacB26) gave poor amplification in *P. brigantina*, yielding more than 10% missing data. This may be because of poor marker transferability, as most of the above microsatellite markers were developed from genomic data on other *Prunus* species, such as peach, almond, apricot and Japanese plum. The remaining 24 microsatellite markers performed well in *P. brigantina* and were used in this study (Table S2). In our *P. brigantina* sample, the number of alleles (*N*_A_) was 121 (mean of 5.04 per marker) and the number of effective alleles (*N*_E_) 59.57 (mean of 2.48 per marker) (Table S2).

The biologically most relevant genetic clustering of *P. brigantina* was found to be *K*=3: the DeltaK statistics indicated that it was the strongest population subdivision level (Figure S1A) and further increasing *K* yielded many admixed individuals (Figure S2). The three inferred genetic clusters (blue, yellow and orange colours in Figure 2b and S2) corresponded to three French national parks “Queyras”, “Ecrins” and “Mercantour”, respectively. Weak but significant genetic differentiation (mean Jost’s *D*=0.117) was found among these three *P. brigantina* populations (Table 2). Both the Josts’ *D* and the PCA indicated that the *P. brigantina* “Queyras” cluster was the most differentiated from the two other ones, the “Ecrins” and “Mercantour” clusters being found genetically closer one to each other (Table 2, Figure 3).

**Table 2.**
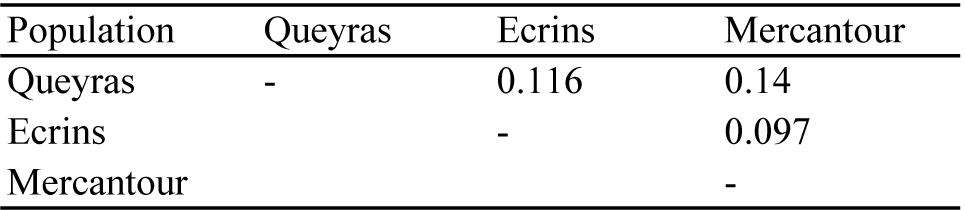
Pairwise population Jost’ *D* of *P. brigantina*.

**Figure 3.**
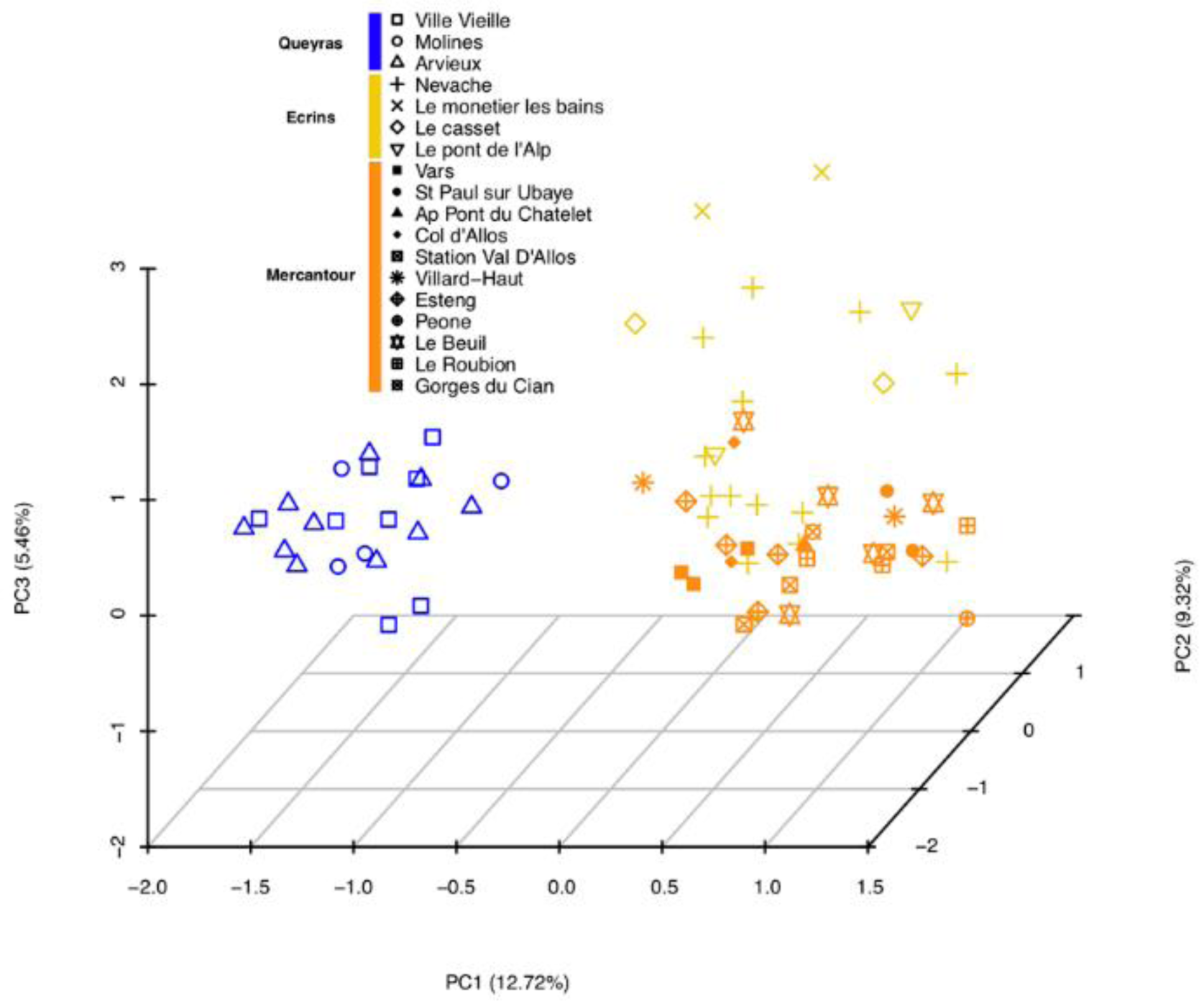
Principal components analysis on *Prunus brigantina*. Colours refer to the genetic clusters inferred from the STRUCTURE analysis, according to the barplots at *K*=3 in Figure S2.

An additional subdivision of the Ecrins cluster was found at *K*=8, revealing differentiation between the dark blue and yellow clusters (Figure S2). The Mantel test on the three *P. brigantina* clusters indicated no significant relationship between genetic differentiation and geographic distance (*P*=0.308, by Monte Carlo permutation tests, Figure S3), indicating a lack of isolation by distance.

### Genetic relationships between *P. brigantina* and other Prunophora species

To obtain a better understanding of the genetic relationships between *P. brigantina* and other Armeniaca species, we combined the current *P. brigantina* data with a former Armeniaca dataset built with the same 24 microsatellite markers (Liu et al. 2019) (Tables 1 and S1c). We performed a Bayesian clustering analysis on the full Armeniaca dataset, including wild and cultivated *P. armeniaca, P. sibirica, P. mandshurica* and *P. mume* (Table 1). We obtained a similar structure as the one described in (Liu et al. 2019) for the previous dataset, and *P. brigantina* differentiated in a distinct cluster, from *K*=3 and above (yellow colour in Figure S4). No gene flow with other species of the section Armeniaca was detected (i.e. no individuals who would have admixed ancestry between the yellow cluster and other clusters), in particular in between wild *P. brigantina* and cultivated apricots which, yet, partly share habitats over Western Europe (Figure S4).

We further questioned the genetic relationship of *P. brigantina* with other members of the Prunophora subgenus, i.e. species of the Prunus and Prunocerasus sections. Because *P. cerasifera* (cherry plum), a species of the Prunus section, is partly sharing habitats with *P. brigantina*, we significantly extended the sampling of *P. cerasifera* species compared to (Horvath et al. 2008), including accessions from the cherry plum native area, i.e. Caucasia and Central Asia, to obtain a better representation at the species level. We then explored the genetic differentiation of this species over its Eurasian distribution. We found genetically differentiated clusters of cherry plums, with contrasted geographical distributions from Central Asia to Europe (detailed results are presented in the supplemental information ‘*Prunus cerasifera* diversity and population structure analysis). Caucasia appears to be a diversification center of wild cherry plums, with two distinct genetic clusters that may result from geographical isolation. This dataset was later merged with representatives of the Prunus, Armeniaca and Prunocerasus sections, to infer the origin of *P. brigantina* and its genetic relationships with species of the Prunophora subgenus (Tables 1, S1b and S1c). In the following step, we focused on species that shared, partly, habitats with *P. brigantina*, i.e. *P. cerasifera* and cultivated *P. armeniaca*, together with other Armeniaca (*P. mume*), Prunus (*P. salicina*) and Prunocerasus species. For this, we used genotyping data based on 23 microsatellite markers (see the supplemental information ‘*Prunus cerasifera* diversity and population structure analysis’).

The delta *K* peaked at *K*=3, indicating that this was the strongest level of population subdivision (Figure S1B). However, further relevant clustering was observed at higher *K* values (Figure S5). From *K*=7 and above, all taxonomic species separated in specific clusters: green for *P. brigantina*, pink for *P. armeniaca*, blue for *P. cerasifera*, grey for *P. mume*, orange for *P. salicina* and black for Prunocerasus (Figure S5). Again, we could not find any admixture footprints between *P. brigantina* and other Prunus species, while there may be some footprints of introgression from *P. cerasifera* into *P. salicina* (see admixed individuals indicated by blue stars in Figure S5), although the blue and orange heterogeneous bars may alternatively result from low assignment power due to the low number of *P. salicina* individuals.

We further explored the genetic differentiation and relationships among all Prunophora samples using an unrooted weighted neighbour-joining tree (Figure 4). In the tree, the delimitation of *P. brigantina* as a distinct species from other apricot and plum taxonomic species was well supported (100% bootstrap support). *Prunus brigantina* trees appeared genetically closer to the Armeniaca species (*P. armeniaca* and *P. mume*) than to other Prunus and Prunocerasus species, which is consistent with Rehder’s taxonomy. The principal component analysis (PCA) supported the differentiation of *P. brigantina* from other species of the Armeniaca section, and from the Prunus and Prunocerasus sections (Figure 5). Both the NJ tree and the PCA indicated that plum species (*P. cerasifera* and *P. salicina*) were partly overlapping, in particular the cultivated Japanese plums and cherry plums; the wild *P. salicina* trees in contrast appeared well separated from *P. cerasifera* (Figures 4 and 5). The overlapping may be the result of low power to distinguish the groups based on few individuals or of hybridization between cultivated trees. One particular case of hybridization is P0489, cv. Rutland plumcot. Breeders’ information indicates that it is a hybrid between plum and apricot. In our structure barplots, NJ tree and PCA (Figures S5, 4 and 5), P0489 in fact appeared admixed between the two plum species, *P. salicina* and *P. cerasifera*, and not with apricot.

**Figure 4.**
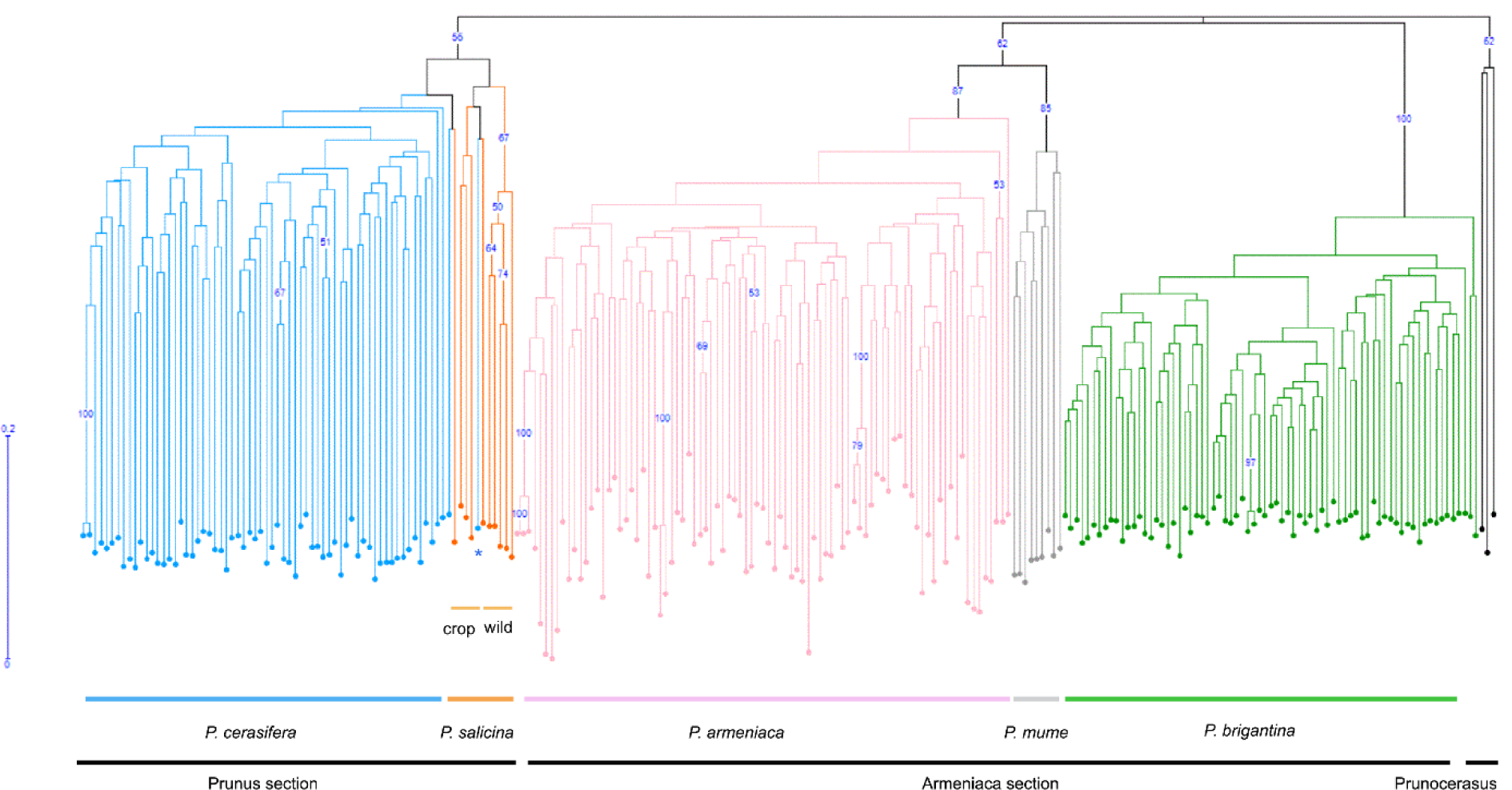
Unrooted weighted neighbour-joining (NJ) tree of *Prunus brigantina* and other Prunophora species. The species are represented by the same colour as the ones used in STRUCTURE barplots (*K*=8, Figure S5). The NJ tree was built with DARwin, bootstrap support values were obtained from 30,000 repetitions. Bootstrap values when greater than 50% are shown above the branches. (*) corresponds to the P0489 plumcot sample. Classification into sections was made according to Krüssmann (1978) and Reales et al (2010).

**Figure 5.**
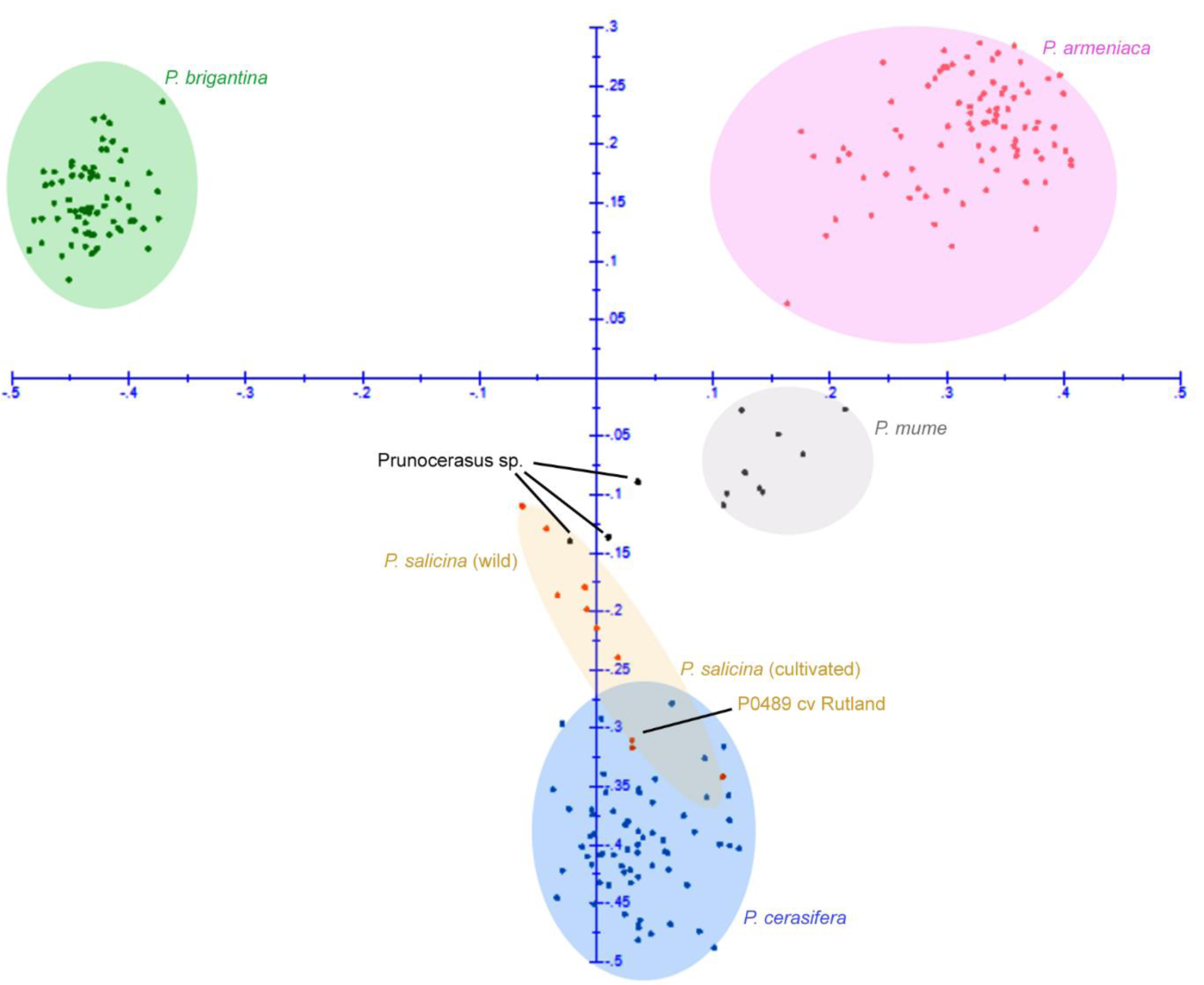
Principal components analysis (PCA) on five Prunophora species performed with DARwin. The sampling for this analysis included *P. cerasifera* (*N*=66) in blue, *P. armeniaca* (*N*=87) in pink, *P. brigantina* (*N*=73) in green, the Chinese apricot tree *P. mume* (*N*=9) in grey and Japanese plum, *P. salicina* (*N*=10) in orange. Black dots correspond to *Prunocerasus* species (*P. mexicana, P. munsoniana* and *P. maritima*). Colours refer to the genetic clusters inferred from the STRUCTURE analysis, according to the barplots at *K*=8 in Figure S5.

### Construction of a *P. brigantina* core collection

We used the COREFINDER program to identify the smallest core collection that would be sufficient to capture the whole diversity detected based on our 24 microsatellite markers. Based on the maximizing strategy implemented in COREFINDER, we propose the use of a core set of 36 individuals (∼49% of the whole *P. brigantina* sample) that captures 100% of the detected diversity (Figure 6, Table S3). Pairwise comparisons using Mann-Whitney U tests showed no significant differences in diversity indexes *(I, H*_*O*_, and *H*_*E*_) between the *P. brigantina* entire Alpine sample (*N*=71) and the core collection (*N*=36) (Tables S2, S4 and S5). This indicates that our core collection can be used as an *ex-situ* germplasm repository.

**Figure 6.**
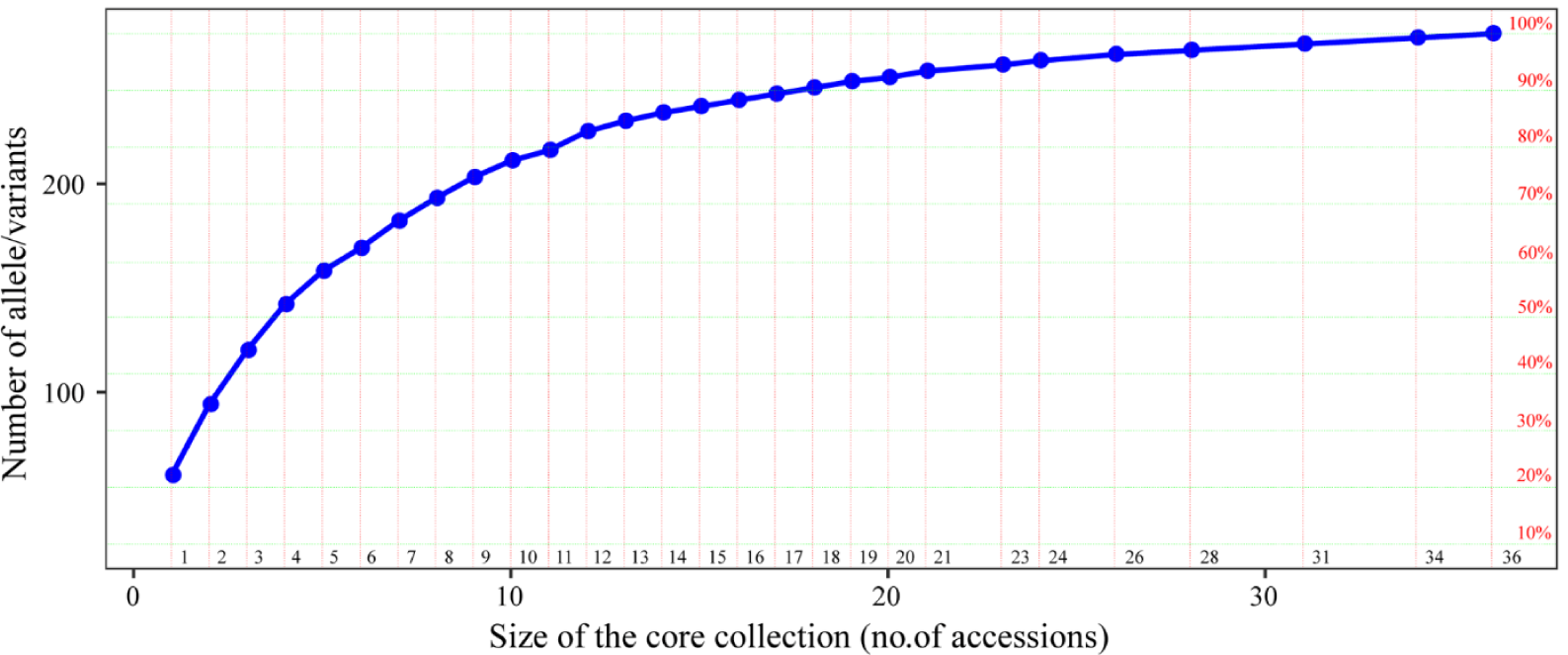
Identification of the core collection of *Prunus brigantina* population based on the strategy maximizing allelic diversity. The genetic diversity in terms of number of alleles (left) or percentage of variation compared to the whole dataset (right) is plotted for different core collection sizes. Details on the accessions retained for each percentage rate are presented in Table S3.

## Discussion

The current study showed that *P. brigantina* is still found in a few Alpine valleys, along the border between France and northwest Italy, where it grows above 1,400 m altitude as single isolated trees (except for the plateau of Nevache, where they are present as a denser population), in arid places such as shrub thickets. In France, it is confined to the three southeastern departments of Alpes-Maritimes, Alpes-de-Haute-Provence and Hautes-Alpes. The sustainability of *P. brigantina* habitat is threatened by forest fragmentation. This raises the question of the long-term conservation of this species and no germplasm accessions of *P. brigantina* are reported by EURISCO to be held in European ex-situ collections. Because large field collections of perennial crops are expensive to maintain, the identification of a restricted number of representatives of *P. brigantina* population for *ex situ* conservation would be very useful in the perspective of Alpine ecosystem restauration and future breeding programs. Core collections are representative subsets of germplasm collections that are developed to improve the efficiency of germplasm evaluation while increasing the probability of finding genes of interest (Simon and Hannan 1995). Therefore, our current core collection will serve in the future for *P. brigantina* conservation as well as for stone fruit breeding programs benefiting from *P. brigantina* resilience characteristics, especially in a context of Mediterranean climate changes. However, the most efficient strategy for biodiversity conservation remains the preservation of the natural habitat of endangered species.

Thanks to an extensive dataset of Prunophora species, we also questioned here the genetic relationships of *P. brigantina* with other species of the Prunus and Armeniaca sections. Species of the Prunocerasus section were not integrated in the analysis except as outgroups because they are naturally distributed on different continents and do not overlap in their respective natural habitats with *P. brigantina*. Through Bayesian analyses, *P. brigantina* appears as a *bona fide* species, clearly distinct from other apricot species and from plum species, with no footprint of admixture. Our results are in accordance with previous studies that indicate a clear differentiation of *P. brigantina* from other Armeniaca apricot species (i.e. *P. armeniaca* and *P. mume*) but do not support a close relationship with species of the Prunus section (Chin et al. 2014; Reales et al. 2010; Shi et al. 2013; Zhebentyayeva et al. 2019). This might be due to the fact that our sampling covers a larger diversity panel than in the former studies, both in *Armeniaca* and *Prunus* sections, *P. brigantina* included. Indeed, sampling only one or two individuals per species is expected to lower the accuracy of phylogenetic analyses (Wiens and Servedio 1997). In our analyses, *P. brigantina* was closer to species of the Armeniaca section than to the Prunus section. While *P. brigantina* should still be considered as an Armeniaca species, it has diverged from *P. armeniaca* long before *P. mume*, thus representing the most genetically distant apricot-related species within the Armeniaca section (Hagen et al. 2002; Liu et al. 2019).

Contradictory results had been obtained from a phylogeny of Eurasian plum species based on chloroplast DNA sequences (Reales et al. 2010), where *P. brigantina* grouped together with European Prunus species, such as the polyploid *P. spinosa, P. insititia* and *P. domestica*, and the diploid *P. ramburii* Boiss.species; it was clearly separated from *P. armeniaca* (apricot). The proximity in chloroplast genotypes between *P. brigantina* and the polyploid Prunus species might indicate the Alpine plum as a parental contributor in interspecific hybridization of polyploid Prunus species (Zhebentyayeva et al. 2019). Organelles are however known to introgress much more often than nuclear DNA and chloroplast genealogies are often discordant from nuclear phylogenies (Coyne and Orr 2004).The other plum species that grouped with *P. brigantina* in chloroplast genealogy, *P. ramburii*, is a relict, wild species endemic in the southern Spanish mountains (Sierra Nevada and Sierra Baz) While its distribution is in Europe, it does not overlap with that of *P. brigantina.* Hence, its morphological features are distinct from Alpine plum, forming bushes with tiny, blue/violet drupes and narrow leaves (http://www.anthos.es/index.php?lang=en). Therefore, the incongruence between our results with those obtained earlier based on the chloroplast genome echoes the conclusions of others that despite the many advantages and widespread use of chloroplast DNA in phylogenetic studies, caution has to be taken in the use of organellar variation for inferring phylogeny (Doyle 1992; Lee-Yaw et al. 2019; Soltis and Kuzoff 1995).

Nevertheless, by extending the sampling set of both *P. brigantina* and plum species, our study provides compelling evidence that *P. brigantina* grouped in the Armeniaca section. It illustrates the importance of the sample size and sampling design that encompasses here a larger genetic diversity at the species level than in previous studies (Hagen et al. 2002; Horvath et al. 2008; Reales et al. 2010; Zhebentyayeva et al. 2019). It also questioned the relevance of the classification into sections of the Prunophora subgenus, at least for the Eurasian sections, i.e. Armeniaca and Prunus. Species of the two sections are sharing habitats and they are interfertile, in particular between diploid species, thus resulting in a number of hybrids and probably new species (Cici and Van Acker 2010; Layne and Sherman 1986). Although the genetic differentiation of the Prunus and Armeniaca sections from the Prunocerasus section is clear (Krüssmann 1978), the relationships among taxa of the two Eurasian sections are not well resolved as illustrated by the role of cross taxa hybridization in Japanese apricot (*P. mum*e) adaptive evolution (Numaguchi et al. 2020). The previous controversial classification of *P. brigantina* either in the Armeniaca section or in the Prunus section reflects the difficulty of assigning a clear barrier between species of those two sections; an analysis of the entire subgenus using a shared set of same nuclear markers could provide greater resolution and would place the findings presented here into a Prunophora-wide perspective.

## Conclusion

In this study, we found a low level of genetic diversity in natural *P. brigantina* populations and identified three genetically differentiated populations, in the Ecrins, Queyras and Mercantour national parks, respectively. We further successfully established in Bordeaux a core collection of 36 individuals representing the *P. brigantina* diversity that will be publicly available through the French Genetic Resources Center. In addition, a population NJ tree did not support a close relationship between *P. brigantina* and the other Prunus species, *P. brigantina* being closer to Armeniaca species whilst remaining clearly distinct. While most of the fruit species originate from Asia or America, many crop wild relatives still exist both in their center of origin and along their dispersal routes. For example, in pit and stone fruits, several *Prunus, Malus* and *Pyrus* wild species are endemic in Europe and are often threatened by the rapid changes of land use (Cornille et al. 2013a; Welk et al. 2016). To inform *in situ* and *ex situ* conservation measures and add value to fruit tree genetic resources, we recommend in-depth characterization of those wild relatives, similarly to the current study in the Alpine plum.

## Supporting information

Supplemental information on Prunus cerasifera diversity

## Acknowledgements

S.L. is a recipient of a Chinese Scholarship Council PhD grant. Molecular analysis was performed at the GenoToul Get-PlaGe (INRAE Center of Toulouse) and GENTYANE (INRAE Center of Clermont-Ferrand) platforms. The authors wish to acknowledge all the people who helped in collecting the samples, in particular, collaborators from Le Plantivore at Château Ville-vieille, Histoire de Confiture at Plampinet, Nevache, C. Gatineau from Cervières and the managers of the Queyras and Mercantour national parks. We thank the curators of the French Genetic Resources Centre (Marine Delmas) and of the US ARS-USDA repository (John Preece); we acknowledge the care of the plants at the UMR BFP (INRAE) by Jean-Philippe Eyquard and Pascal Briard.

## Statements

Appropriate permissions from responsible authorities for collecting and using *Prunus* samples from Central Asia and Caucasia were obtained by the local collaborators. The official authorization for the survey and sampling of *P. brigantina* genetic resources is registered and accessible through the following link: https://absch.cbd.int/database/ABSCH-IRCC-FR-246978. The rest of the samples were kindly provided, with due authorizations, by the curators of the French INRAE Genetic Resources Centre (GRC) and the US ARS-USDA repository, further details are available on their respective databases.

## Data availability

The datasets generated by the current study, *i.e*. the SSR genotyping, are available at the INRAE data portal (https://data.inrae.fr/) where they can be freely retrieved.

## Captions for the supplementary Figures presented in a separate PDF file

**Figure S1. DeltaK plot as a function of K for the *Prunus brigantina* (A) and Prunophora (B) dataset.**

**Figure S2. Bayesian clustering on *Prunus brigantina* samples in the French Alps.**

*Prunus brigantina* dataset included 71 individuals sampled from the French Alps and two samples from the French GRC repository. Each individual is represented by a vertical bar, partitioned into *K* segments representing the inferred proportions of ancestry of its genome.

**Figure S3. Isolation by distance (IBD) test in *Prunus brigantina***.

a. Distribution of correlation values between genetic and geographic distances under the assumption of lack of isolation by distance, drawn from permutations; the observed value of the correlation between the distance matrices, represented by the black diamond, falls within the expected distribution which indicates the lack of isolation by distance pattern.

b. Pairwise Edwards’ distances plotted against Euclidean geographic distances, with local density of points plotted using a two-dimensional kernel density estimate, displayed in colour from white to red. The solid line represents the fitted linear regression between Edwards’ genetic and Euclidean geographic distances.

**Figure S4. Bayesian analysis on Armeniaca and wild *Prunus brigantina* accessions.**

Genetic subdivision among Armeniaca species, *P. brigantina* included, was inferred with STRUCTURE with 24 microsatellite markers. The 648 samples belong to the six Armeniaca species as follows: *P. brigantina* (*N*=73), *P. armeniaca* (European and Chinese cultivated *N*=270 and wild, *N*=204), *P. sibirica* (*N*=84), *P. mume* (*N*=9), *P. mandshurica* (*N*=8). Each individual is represented by a vertical bar, partitioned into *K* segments representing the inferred proportions of ancestry of its genome. Species and origin of the accessions are indicated on the top of the figure.

**Figure S5. Bayesian analysis on the *Prunus brigantina* dataset together with an extended Prunophora dataset.**

Genetic subdivision among Armeniaca, Prunus and Prunocerasus species was inferred with STRUCTURE with 23 microsatellite markers (supplemental information for the list of markers). The 226 samples belong to three different Prunophora species including *P. brigantina* (*N*=73), *P. cerasifera* (*N*=66), *P. armeniaca* (*N*=87), *P. salicina* (*N*=10), *P. mume* (*N*=9), *P. mexicana* (*N*=1), *P. munsoniana* (*N*=1), *P. maritima* (*N*=1), *P. americana* (*N*=1) and *P. subcordata* (*N*=1). The blue stars (*), at the bottom of the bar plots, correspond to Japanese plums (*P. salicina*) admixed with *P. cerasifera*.

## Legends for the supplementary tables presented in a separate PDF file

**Table S1a. Sampling locations, geographic regions and assigned genetic cluster of *Prunus brigantina* in the French Alps.**

FR for an origin from the French Alps. Sampling site is indicated in GPS coordinates, N for North, E for East.

**Table S1b. Sampling locations, geographic regions and/or germplasm repositories of *Prunus cerasifera* samples.**

^1^ Species affiliation as indicated by the curator of the germplasm collection where the sample is maintained or as identified *in situ*. ^2^ Sampling location in decimal degrees. ^3^ Origin as indicated in the database of the germplasm repository. n/a, not applicable because admixed and thus not used in the correlation tests

**Table S1c: List of individuals included in the different datasets.**

^1^ FR refers to France, AZ to Azerbaijan, CH to China, KR to Kyrgyzstan, KZ to Kazakhstan, OUZ to Uzbekistan, TCH to Czech republic (Lednice repository), TURC to Turkey (Malatya repository), US to USA (ARS-USDA Prunus germplasm repository). For more details, see Liu et al (2019). Accession numbers starting with A indicate apricot cultivars and with P, plum cultivars, as displayed in the French GRC database. The sign (-) means that the sample is maintained in germplasm repository and was not collected *in situ*. The cross in the last four columns (dataset 1 to 4) means that this sample was used in the corresponding dataset.

**Table S2. Analysis of genetic variability from microsatellite markers for *Prunus brigantina* population.**

*Na*: number of different alleles, and *Ne*: number of effective alleles. *I*: Shannon diversity index. *He* and *Ho*: expected and observed heterozygosities.

**Table S3. The description of individuals retained for the core collection of *Prunus brigantina***

**Table S4. Genetic variability of microsatellite markers for the *Prunus brigantina* core collection.**

*Na*: number of different alleles, and *Ne*: number of effective alleles. *I*: Shannon diversity index. He and *Ho*: expected and observed heterozygosities.

**Table S5. Mann-Whitney U tests (two-tailed) between the whole *Prunus brigantina* dataset and its core collection.**

## Supplemental information presented in a separate PDF file

**Supplemental information ‘*Prunus cerasifera* diversity and population structure analysis’**

## References

Allendorf FW, Luikart GH, Aitken SN (2012) Conservation and the Genetics of Populations., 2nd Edition edn. Wiley-Blackwell Ltd,

Branca F, Donnini D (2011) Prunus brigantina. vol e.T172164A121228349.. doi: http://dx.doi.org/10.2305/IUCN.UK.2011-1.RLTS.T172164A6840507.en.

Chin S-W, Shaw J, Haberle R, Wen J, Potter D (2014) Diversification of almonds, peaches, plums and cherries – Molecular systematics and biogeographic history of Prunus (Rosaceae). Molecular Phylogenetics and Evolution 76:34–48 doi: https://doi.org/10.1016/j.ympev.2014.02.024

Cici S, Van Acker R (2010) Gene flow in Prunus species in the context of novel trait risk assessment. Environmental biosafety research 9:75–85 doi: 10.1051/ebr/2010011

Cornille A et al. (2013a) Postglacial recolonization history of the European crabapple (Malus sylvestris Mill.), a wild contributor to the domesticated apple. Molecular Ecology 22:2249–2263 doi: 10.1111/mec.12231

Cornille A, Gladieux P, Giraud T (2013b) Crop-to-wild gene flow and spatial genetic structure in the closest wild relatives of the cultivated apple. Evolutionary Applications 6:737–748 doi: 10.1111/eva.12059

Couplan F (2009) Le régal végétal: plantes sauvages comestibles. Collection L’encyclopédie des plantes sauvages comestibles et toxiques de l’Europe.

Coyne JA, Orr HA (2004) Speciation. OUP USA,

Decroocq S et al. (2016) New insights into the history of domesticated and wild apricots and its contribution to Plum pox virus resistance. Molecular Ecology 25:4712–4729 doi: 10.1111/mec.13772

Doyle JJ (1992) Gene Trees and Species Trees: Molecular Systematics as One-Character Taxonomy. Systematic Botany 17:144–163 doi: 10.2307/2419070

Dupouy J (1959) Le prunier de Briançon ou Marmottier (Prunus brigantiaca Vill.) et les huiles de marmotte. Bull SAJA 141:69–71

Earl DA, vonHoldt BM (2012) STRUCTURE HARVESTER: a website and program for visualizing STRUCTURE output and implementing the Evanno method. Conservation Genetics Resources 4:359–361 doi: 10.1007/s12686-011-9548-7

Escribano P, Viruel M, Hormaza J (2008) Comparison of different methods to construct a core germplasm collection in woody perennial species with simple sequence repeat markers. A case study in cherimoya (Annona cherimola, Annonaceae), an underutilised subtropical fruit tree species. Annals of Applied Biology 153:25–32 doi: 10.1111/j.1744-7348.2008.00232.x

Evanno G, Regnaut S, Goudet J (2005) Detecting the number of clusters of individuals using the software structure: a simulation study. Molecular Ecology 14:2611–2620 doi: 10.1111/j.1365-294X.2005.02553.x

Fahrig L (2003) Effects of Habitat Fragmentation on Biodiversity. Annual Review of Ecology, Evolution, and Systematics 34:487–515 doi: 10.1146/annurev.ecolsys.34.011802.132419

Frankel OH, Brown AHD Current Plant genetic resources today - a critical appraisal. In: Proceedings of the XV International Congress of Genetics. Crop Genetic Resources: Conservation and Evaluation, New Delhi, India, 1983. Oxford & IBH publishing CO., pp 241–256

Glaszmann JC, Kilian B, Upadhyaya HD, Varshney RK (2010) Accessing genetic diversity for crop improvement. Current Opinion in Plant Biology 13:167–173 doi: 10.1016/j.pbi.2010.01.004

Govindaraj M, Vetriventhan M, M S (2014) Importance of Genetic Diversity Assessment in Crop Plants and Its Recent Advances: An Overview of Its Analytical Perspectives. Genetics Research International 2015 doi: 10.1155/2015/431487

Hagen L, Khadari B, Lambert P, Audergon J-M (2002) Genetic diversity in apricot revealed by AFLP markers: species and cultivar comparisons. Theoretical and Applied Genetics 105:298–305 doi: 10.1007/s00122-002-0910-8

Heled J, Drummond AJ (2010) Bayesian Inference of Species Trees from Multilocus Data.Molecular Biology and Evolution 27:570–580 doi: 10.1093/molbev/msp274

Horvath A, Christmann H, Laigret F (2008) Genetic diversity and relationships among Prunus cerasifera (cherry plum) clones. Botany 86:1311–1318 doi: 10.1139/b08-097

Jakobsson M, Rosenberg NA (2007) CLUMPP: a cluster matching and permutation program for dealing with label switching and multimodality in analysis of population structure. Bioinformatics 23:1801–1806 doi: 10.1093/bioinformatics/btm233

Jombart T, Ahmed I (2011) Adegenet 1.3-1: new tools for the analysis of genome-wide SNP data. Bioinformatics 27:3070–3071 doi: 10.1093/bioinformatics/btr521

Krüssmann G (1978) Manual of cultivated broad-leaved trees and shrubs. vol 3 Pru-Z. Timber Press, Portland

Layne RE, Sherman CWB (1986) Interspecific hybridization of Prunus. HortScience 21:48–51

Lee-Yaw JA, Grassa CJ, Joly S, Andrew RL, Rieseberg LH (2019) An evaluation of alternative explanations for widespread cytonuclear discordance in annual sunflowers (Helianthus). New Phytologist 221:515-526 doi: 10.1111/nph.15386

Li DZ, Pritchard HW (2009) The science and economics of ex situ plant conservation. Trends in plant science 14:614–621

Ligges U, Mächler M (2003) Scatterplot3d - an R package for Visualizing Multivariate Data. Journal of Statistical Software 8:1–20 doi: 10.17877/de290r-15194

Liu S et al. (2019) The complex evolutionary history of apricots: Species divergence, gene flow and multiple domestication events. Molecular Ecology 28:5299-5314 doi: 10.1111/mec.15296

Meirmans PG, Van Tienderen PH (2004) Genotype and genodive: two programs for the analysis of genetic diversity of asexual organisms. Molecular Ecology Notes 4:792–794 doi: 10.1111/j.1471-8286.2004.00770.x

Noble V, Van Es J, Michaud H, Garraud L (2015) Liste Rouge de la flore vasculaire de Provence-Alpes-Côte d’Azur. Provence-Alpes-Côte d’Azur, France

Numaguchi K, Akagi T, Kitamura Y, Ishikawa R, Ishii T (2020) Interspecific introgression and natural selection in the evolution of Japanese apricot (Prunus mume). bioRxiv:2020.2006.2023.141200 doi: 10.1101/2020.06.23.141200

Peakall R, Smouse P (2012) GenAIEx V5: Genetic Analysis in Excel. Populations Genetic Software for Teaching and Research. Bioinformatics (Oxford, England) 28:2537-2539 doi: 10.1093/bioinformatics/bts460

Perrier X, Jacquemoud-Collet JP (2006) DARwin software. http://darwin.cirad.fr/.

Pignatti S (1982) Flora d’Italia. vol 2. Edagricole, Bologna, Italy

Pritchard JK, Stephens M, Donnelly P (2000) Inference of Population Structure Using Multilocus Genotype Data. Genetics 155:945–959

Reales A, Sargent DJ, Tobutt KR, Rivera D (2010) Phylogenetics of Eurasian plums, Prunus L. section Prunus (Rosaceae), according to coding and non-coding chloroplast DNA sequences. Tree Genetics & Genomes 6:37-45 doi: 10.1007/s11295-009-0226-9

Rehder A (1940) Manual of cultivated trees and shrubs hardy in North America. vol 1. Biosystematics, floristic & phylogeny series. Collier Macmillan Ltd, New York

Rosenberg NA (2004) Distruct: a program for the graphical display of population structure. Molecular Ecology Notes 4:137–138 doi: 10.1046/j.1471-8286.2003.00566.x

Schoen DJ, Brown AHD (1993) Conservation of allelic richness in wild crop relatives is aided by assessment of genetic markers. Proceedings of the National Academy of Sciences 90:10623–10627 doi: 10.1073/pnas.90.22.10623

Shannon CE (1948) A Mathematical Theory of Communication. Bell System Technical Journal 27:379–423 doi: 10.1002/j.1538-7305.1948.tb01338.x

Shi S, Li J, Sun J, Yu J, Zhou S (2013) Phylogeny and Classification of Prunus sensu lato (Rosaceae). Journal of Integrative Plant Biology 55:1069–1079 doi: 10.1111/jipb.12095

Simon CJ, Hannan RM (1995) Development and Use of Core Subsets of Cool-season Food Legume Germplasm Collections. HortScience 30:907C doi: 10.21273/hortsci.30.4.907c

Soltis DE, Kuzoff RK (1995) Discordance between nuclear and chloroplast phylogenies in the Heuchera group (Saxifragaceae). Evolution 49:727–742 doi: 10.1111/j.1558-5646.1995.tb02309.x

Szpiech ZA, Jakobsson M, Rosenberg NA (2008) ADZE: a rarefaction approach for counting alleles private to combinations of populations. Bioinformatics 24:2498–2504 doi: 10.1093/bioinformatics/btn478

Takeda T, Shimada T, Nomura K, Ozaki T, Haji T, Yamaguchi M, Yoshida M (1998) Classification of Apricot Varieties by RAPD Analysis. Engei Gakkai Zasshi 67:21–27 doi: 10.2503/jjshs.67.21

Takezaki N, Nei M, Tamura K (2014) POPTREEW: web version of POPTREE for constructing population trees from allele frequency data and computing some other quantities. Mol Biol Evol 31:1622–1624 doi: 10.1093/molbev/msu093

Tison J-M, De Foucault B (2014) Flora Gallica - Flore de France. Biotope, Mèze

Villars D (1786) Histoire des plantes de Dauphiné: Contenant une Préface Historique, un Dictionnaire des Termes de Botanique, les Classes, les Familles, les Genres, & les Herborisations des Environs de Grenoble, de la Grande Chartreuse, de Briançon, de Gap & de Montelimar. vol 1. Prevost, Paris

Welk E, de Rigo D, Caudullo G (2016) Prunus avium in Europe: distribution, habitat, usage and threats. In: San-Miguel-Ayanz J, de Rigo D, Caudullo G, Houston Durrant T, A. M (eds) European Atlas of Forest Tree Species. Publication Office of the European Union, Luxembourg,

Wiens JJ, Servedio MR (1997) Accuracy of Phylogenetic Analysis Including and Excluding Polymorphic Characters. Systematic Biology 46:332–345 doi: 10.1093/sysbio/46.2.332

Zhebentyayeva T et al. (2019) Genetic characterization of worldwide Prunus domestica (plum) germplasm using sequence-based genotyping. Horticulture Research 6 doi: 10.1038/s41438-018-0090-6

