## Supplemental information on Prunus cerasifera diversity for "Genetic diversity and population structure analyses in the Alpine plum (*Prunus brigantina* Vill.) confirm its affiliation to the Armeniaca section"

**Supplemental Information on *Prunus cerasifera* diversity and population structure analysis for:**

The following Supporting Information is available for this article:



| **Supplemental notes** | **Pages** |
|---|---|
| Supplementary note 1: Genetic variability and population structure in *Prunus cerasifera* population | 2-3 |
| Supplementary note 2: Isolation by distance and genetic differentiation in *Prunus cerasifera* population | 3-4 |
| **Supplemental figures** | |
| Figure SX1. Delta *K* plotted against *K* values for the *Prunus cerasifera* STRUCTURE analysis. | 5 |
| Figure SX2. Bayesian clustering inferred with STRUCTURE for *K*=2 to *K*= 10 for the 66 *Prunus cerasifera* analyzed in this study with 34 microsatellite markers. | 6 |
| Figure SX3. Genetic structure among the 66 *Prunus cerasifera* samples and their distribution over Europe, Caucasia and Central Asia. | 7-8 |
| Figure SX4: Neighbour-joining (NJ) tree based on Nei's standard genetic distance with sample size correction ($D_{ST}$) among genetic groups of *Prunus cerasifera* (*N*=39) and *P. mume* (*N*=9). | 9 |
| **Supplemental tables** | |
| Table SX1. List of the 23 microsatellite markers used in *Prunus cerasifera* diversity analysis and their characteristics: given name of the locus, number of different alleles (*Na*) and number of effective alleles (*Ne*) | 10 |
| Table SX2. Genetic variation within each of the four *Prunus cerasifera* clusters inferred with STRUCTURE at *K*=4 based on 23 microsatellite markers. | 11 |
| Table SX3. Pairwise population matrix of Jost's estimate of differentiation (Jost's *D*) among the four *Prunus cerasifera* clusters | 11 |

### Supplementary note 1: Genetic diversity and population structure in *Prunus cerasifera* population

After genotyping with the twenty-four microsatellite markers retained for *P. brigantina*, one marker (UDP96 018) was removed from the study due to poor amplification. Indeed, this marker presented a high level of missing data (57%). The remaining 23 microsatellites markers were used to assess the genetic diversity at the species level, with a mean value of alleles ($N_A$) = 19.826 and of effective alleles ($N_E$) = 7.611 (Table SX1 in this document) which was three to four times higher than for *P. brigantina* (5.042 and 2.48, respectively, Table S2). Through GENODIVE analysis, one pair of *P. cerasifera* had identical genotypes, i.e. az25 and az239. Identifying and eliminating clonemates in population genetic analyses is important since spurious replicates generate a bias in allele frequencies. Consequently, only one of the two trees was retained for further analysis.

To evaluate the extent of genetic diversity within *P. cerasifera*, genetic subdivision at the intra-specific level was explored, based on the 66 cherry plum samples (Table S1) and the 23 microsatellites markers (Table SX1, in this document). $\Delta K$ was highest at $K$=2 (Figure SX1), at which Central Asian cherry plums separated from the other plums (Figure SX2). However, based on barplots from $K$=2 to $K$=10, the most relevant value of *K, i.e.,* identifying the highest number of well-delimited clusters and above which no new cluster was observed, was $K$=4 (Figure SX2).

At $K$=4, the individuals clustered according to their geographical origin, and with two groups in Caucasia (Figures SX2 and SX3, below). Indeed, when analysing *P. cerasifera* alone, four groups were detected, corresponding to European (red cluster in Figure SX2), Caucasian (two clusters, purple and yellow in Figure SX2) and Central Asian (in blue in Figure SX2) clusters. The additional population found at $K$=4 indicated a subdivision in the Caucasian region, more precisely in the autonomous province of Naxçivan (or Nakhichevan, in yellow, Figure SX3), which is a region of Azerbaijan separated from the rest of the country by Armenia.

Some *P. cerasifera* accessions appeared to be assigned to a genetic cluster not common in the region where the trees were sampled. One example is depicted in Figure SX2, by a star. This sample was collected from a garden in Kazakhstan (kz303, Table S1), but it did not cluster with the other Central Asian accessions, instead clustering with Caucasian cherry plums (in purple, Figure SX2). In consequence, its origin was likely Azerbaijan or other areas of Caucasia, from where it was transferred to Kazakhstan. Indeed, exchanges of biological material between fruit growers, or between curators of germplasm collections and botanical gardens often resulted in the past in the loss of information on the native origin of the accessions, or at least in confusion on the true origin of the plant material, especially when it is vegetatively propagated. In addition, it appears that admixture (between different *P. cerasifera* clusters in Figure XS2 or between *P. cerasifera* and other plum species, in Figure S5) is rather common. Therefore, in order to estimate genetic distance between *P. cerasifera* populations, admixed individuals were removed from the dataset using a 90% assignment threshold to one of the four genetic clusters. At the 90% threshold, we still detected four cherry plum clusters of distinct geographical origins which encompass well assigned, non-admixed accessions from Central Asia ($N$=15), Azerbaijan (Caucasia 1, $N$=13), Naxcivan (Caucasia 2, $N$=6) and Europe ($N$=5) (Table SX2). The European cluster of cherry plums showed widespread admixture footprints with other *Prunus* species (Figure S5). On the contrary, in Caucasia, two distinct clusters were observed with little admixture (Figures SX2 and SX3), most probably due to the fact that the two Azerbaijan regions are isolated from each other by Armenia, thus preventing gene flow between the two *P. cerasifera* populations.

### Supplementary note 2: Isolation by distance and genetic differentiation in *Prunus cerasifera* population

To test if there was a positive correlation between the genetic matrix and geographic distances, i.e., an isolation-by-distance (IBD) pattern, a Mantel test for matrix correspondence was performed using GenAlEx. The genetic ($x$) and geographic ($y$) distances were correlated using both individuals ($R_{xy}$=0.366; $P$-value=0.0001) and populations as data points ($R_{xy}$=0.989; $P$-value=0.042).

The genetic diversity and differentiation among the above four *P. cerasifera* clusters were assessed using only non-admixed accessions (Table SX2). The fixation index ($F_{ST}$ in Table SX2) was highest among Central Asian cherry plums ($F_{ST}$ =0.207 +/-0.069). However, $F_{ST}$ and number of alleles per population ($N_A$) are biased when sample size is small or varies between populations, which is the case here (Table

SX2). In such cases, it is recommended to use instead the Jost's *D* coefficient of differentiation and allelic richness ($A_r$ and $A_p$, Table SX2) through the rarefaction procedure implemented in ADZE (Table SX3). The Caucasian 1 group showed significantly higher allelic richness ($A_r$=3.575 ± 0.160) than the three other *P. cerasifera* populations (Table SX2). The Caucasian 2 cluster showed the lowest genetic diversity ($H_E$=0.553 ± 0.043) and allelic richness ($A_r$=2.601 ± 0.157), as a possible consequence of a bottleneck and founder effect. This result was confirmed by the construction of a tree depicting the relationships among clusters based on the Nei's standard genetic distance ($D_{ST}$) (Figure SX4), which is also consistent with the geographical origins of the *P. cerasifera* populations (Figure SX3). Jost's *D* estimates among the genetic clusters obtained with STRUCTURE after filtering out admixed individuals revealed highly significant genetic differentiation among the four *P. cerasifera* clusters (*P*-values < 0.001), with higher genetic distances among geographically more distant clusters (Jost's *D* ranging from 0.196 to 0.356, *P*-value < 0.005, Table SX3). The highest degree of differentiation was observed between cherry plums from Central Asia and Europe (Jost's *D*=0.361, *P*-value=0.001).

**Figure SX1. Delta *K* plotted against *K* values for the *Prunus cerasifera* STRUCTURE analysis**

The delta K ($\Delta K$) was estimated by Structure Harvester for the full *P. cerasifera* dataset of 66 individuals (STRUCTURE results based on 23 microsatellite markers corresponding to the barplots in Figure SX2).

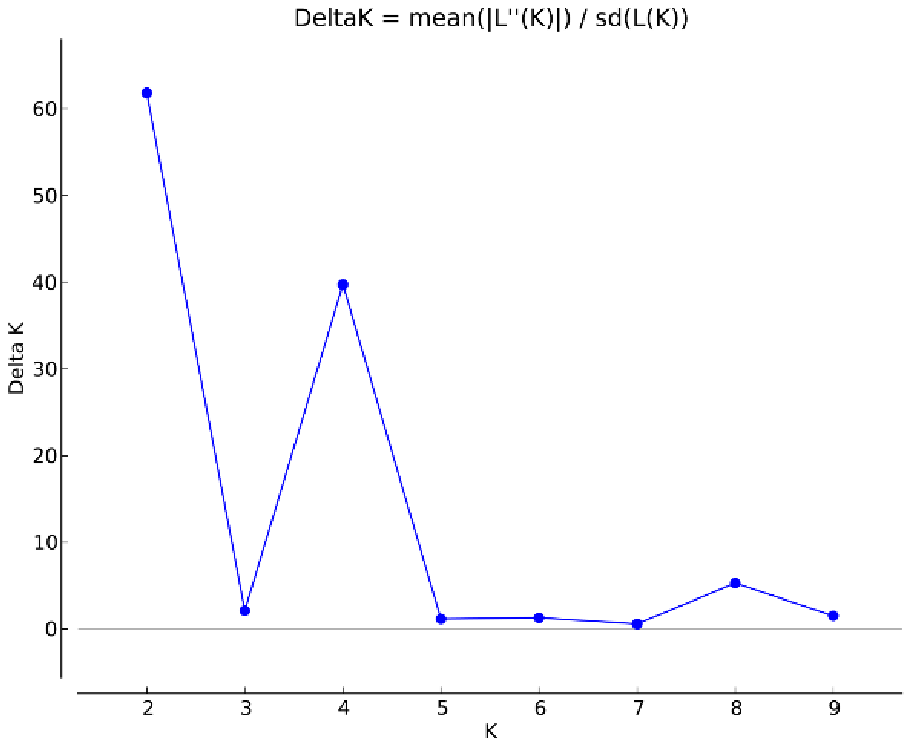

**Figure SX2. Bayesian clustering inferred with STRUCTURE for *K*=2 to *K*= 10 for the 66 *Prunus cerasifera* analyzed in this study with 23 microsatellite markers.**

The 66 *P. cerasifera* accessions include samples from Central Asia (more precisely from Kazakhstan and Kyrgyzstan, *N*=24), Caucasia (*N*=28) and Europe (*N*=14). Each individual is represented by a vertical bar, partitioned into *K* segments representing the proportions of ancestry of its genome in *K* clusters. At the bottom are represented the four main *P. cerasifera* genetic clusters at *K*=4.

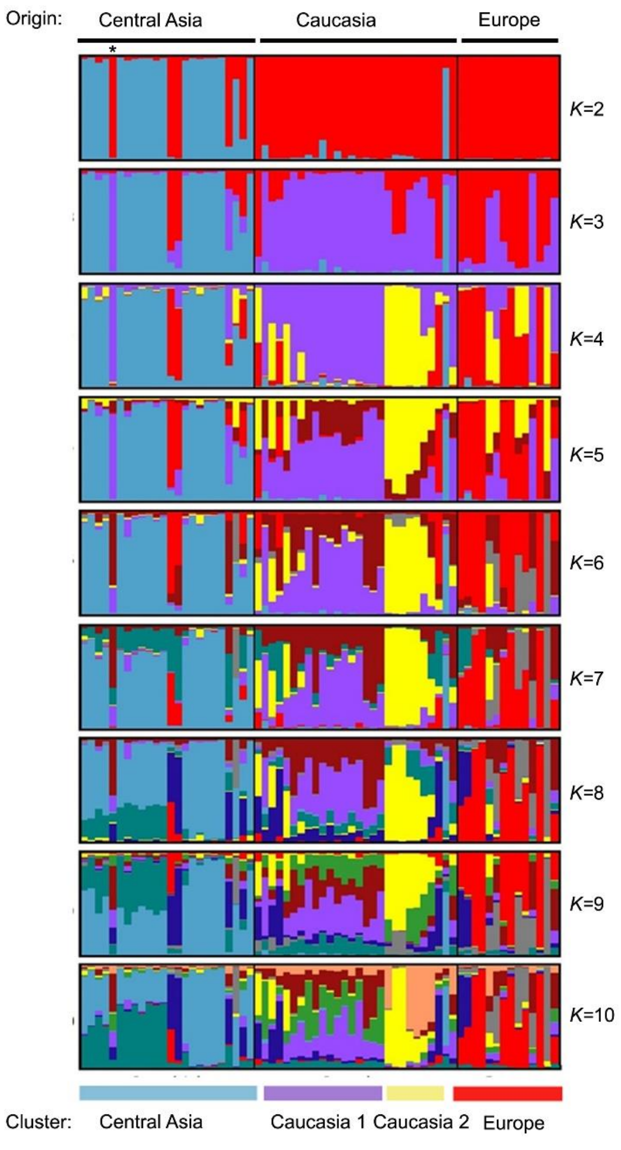

**Figure SX3. Genetic clustering and spatial distribution of *Prunus cerasifera* over Eurasia.**

Genetic structure inferred with STRUCTURE at *K*=4 (Figure SX2), with four well-delimited *P. cerasifera* clusters: European, Caucasian 1 and 2 and Central Asian clusters. The 66 *P. cerasifera* accessions correspond to accessions from Central Asia (*N*=24), Azerbaijan (*N*=24), Russia and Armenia (*N*=4) and Europe (*N*=14). The partial enlarged views represented on the map correspond to wild *P. cerasifera* samples from Caucasia (down), in particular Azerbaijan and Armenia, and from Central Asia (top). Pie chart colors correspond to the colors of Bayesian clustering assignment: red for the European cluster, purple for Caucasian 1 and yellow for Caucasian 2 and blue for the Central Asian cluster.

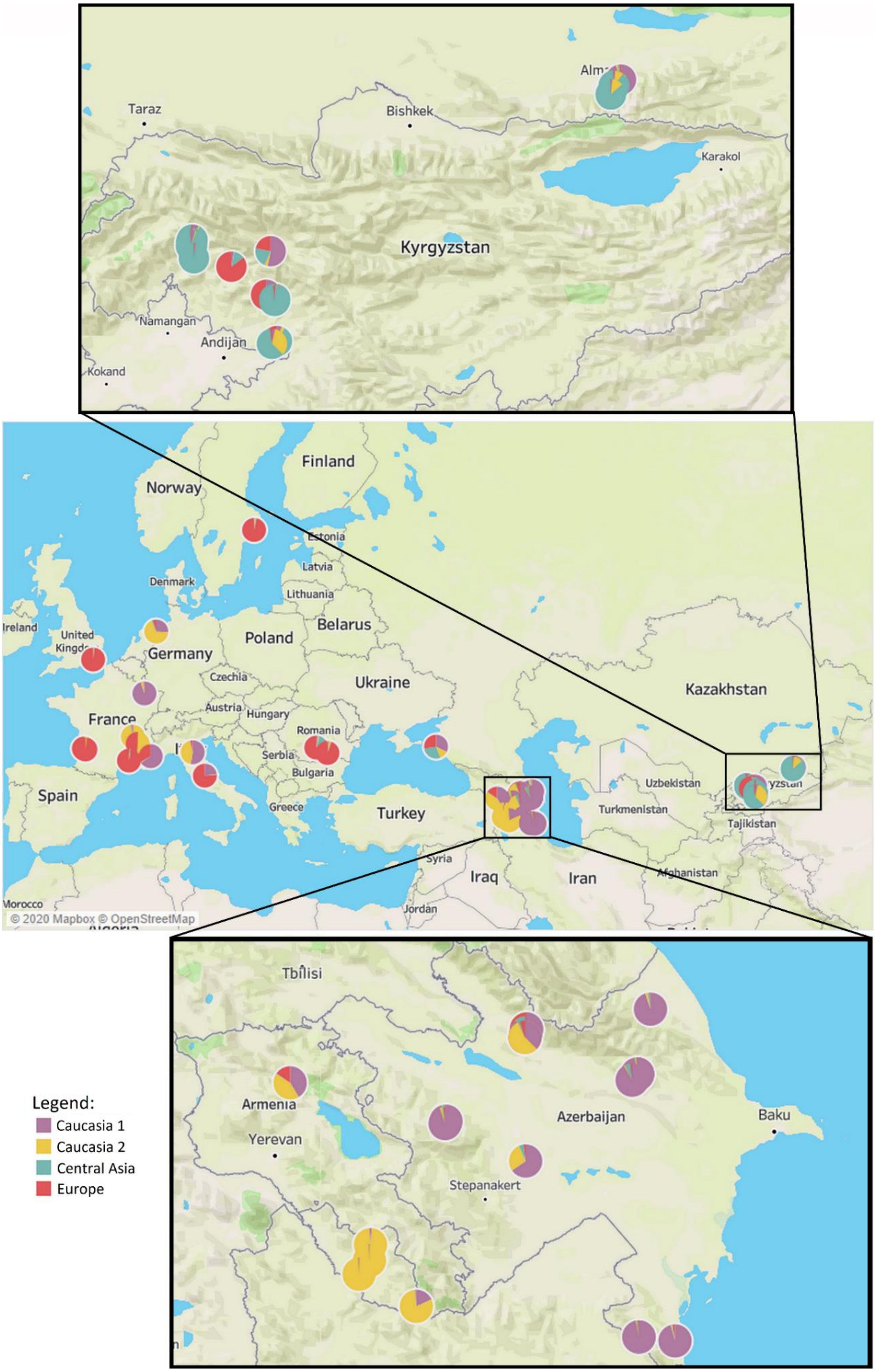
Taraz
Bishkek
Karakol
Kyrgyzstan
Namangan
Andijan
Kokand
Finland
Norway
Estonia
Latvia
Denmark
Lithuania
Ireland
United
Belarus
Poland
Germany
Czechia
Ukraine
Kazakhstan
Austria
Hungary
France
Romania
Serbia
Bulgaria
Spain
Greece
Turkey
Uzbekistan
Turkmenistan
Tajikistan
Syria
Iraq
Iran
Afghanistan
Morocco
Jordan
© 2020 Mapbox © OpenStreetMap
Tbilisi
Armenia
Yerevan
Azerbaijan
Baku
Stepanakert
Legend:
Caucasia 1
Caucasia 2
Central Asia
Europe

**Figure SX4: Neighbour-joining (NJ) tree based on Nei's standard genetic distance with sample size correction ($D_{ST}$) among genetic groups of *Prunus cerasifera* (*N*=39) and the outgroup *P. mume* (*N*=9).**

The present tree was built with PopTree2 and 30,000 bootstraps. The four *P. cerasifera* clusters encompassed accessions with a membership coefficient of 90% in the Bayesian structure analyses.

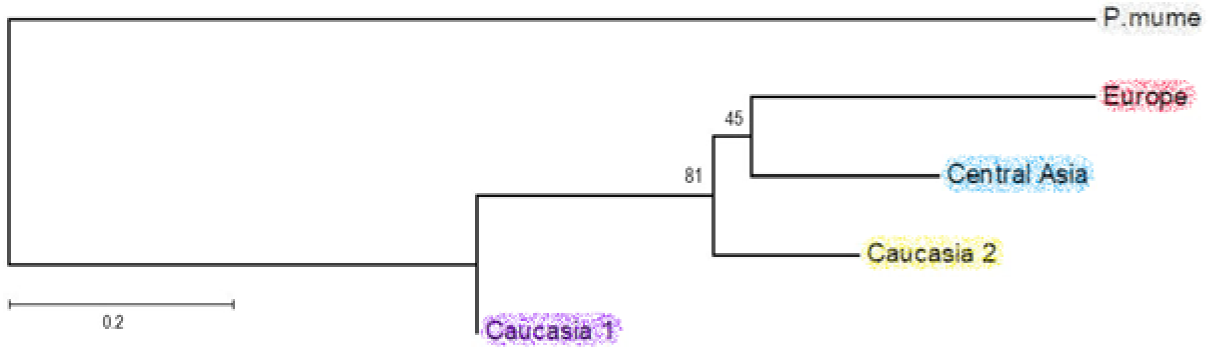

**Table SX1: List of the 23 microsatellite markers used in *Prunus cerasifera* population analyses and their characteristics: marker name, number of alleles (*Na*) and number of effective alleles (*Ne*)**

| Locus | % missing | *Na* | *Ne* |
|---|---|---|---|
| AMPA100 | 0 | 15 | 6.075 |
| AMPA101 | 2 | 24 | 11.347 |
| AMPA103 | 0 | 34 | 16.301 |
| aprigms18 | 16 | 16 | 3.58 |
| BPPCT004 | 0 | 20 | 4.059 |
| BPPCT025 | 0 | 28 | 6.269 |
| BPPCT040 | 0 | 16 | 10.472 |
| CPPCT030 | 1 | 17 | 3.474 |
| CPPCT006 | 0 | 22 | 12.569 |
| CPPCT033 | 0 | 18 | 9.461 |
| EPPCU0532 | 7 | 15 | 5.990 |
| G22 SSR | 0 | 4 | 1.42 |
| SSRLg1_11m52a | 1 | 28 | 11.327 |
| N86B11 SSR3 | 0 | 20 | 6.842 |
| pchgms03 | 0 | 24 | 7.066 |
| PGS-1.21 | 1 | 28 | 12.916 |
| SSR04 AG51 | 6 | 21 | 8.686 |
| SSR5piso4E | 0 | 8 | 2.44 |
| SSR5piso4Ga | 2 | 11 | 3.671 |
| UDA-021 | 0 | 17 | 3.426 |
| UDAp-414 | 0 | 17 | 8.576 |
| UDAp-480 | 4 | 27 | 10.599 |
| UDP98-409 | 2 | 26 | 8.474 |
| **Mean** | | **19.826** | **7.611** |

**Table SX2: Genetic variation within each of the four *Prunus cerasifera* clusters inferred with STRUCTURE at *K*=4 based on 23 microsatellite markers.**

Only individuals assigned to a genetic cluster with a membership proportion greater than or equal to 90% were retained, *i.e.* 39 accessions in total. The number of alleles ($N_A$), the observed heterozygosity ($H_O$), the expected heterozygosity ($H_E$), and the fixation index ($F_{ST}$) were calculated using GenAlEx v6.503, an add-in of Excel. The allelic richness ($A_r$) and the private allelic richness ($A_p$) were calculated using ADZE v1.0. Standard deviations are given in brackets.

| Genetic clusters of *P.cerasifera* | $N$ | $N_A$ | $A_r$ | $A_p$ | $H_O$ | $H_E$ | $F_{ST}$ |
|---|---|---|---|---|---|---|---|
| Central Asia | 15 | 5.956 (0.447) | 2.997 (0.154) | 1.081 (0.130) | 0.530 (0.060) | 0.661 (0.043) | 0.207 (0.069) |
| Caucasia 1 | 13 | 9.348 (0.673) | 3.575 (0.160) | 1.570 (0.158) | 0.655 (0.049) | 0.759 (0.033) | 0.127 (0.055) |
| Caucasia 2 | 6 | 3.522 (0.287) | 2.601 (0.157) | 0.723 (0.139) | 0.488 (0.055) | 0.553 (0.043) | 0.096 (0.071) |
| Europe | 5 | 3.652 (0.318) | 2.775 (0.178) | 1.077 (0.165) | 0.565 (0.065) | 0.578 (0.043) | 0.069 (0.082) |
| Mean | 9.75 | 5.620 (0.334) | 2.987 | 1.113 | 0.560 (0.029) | 0.638 (0.022) | 0.125 (0.035) |

**Table SX3: Pairwise population matrix of Jost's estimate of differentiation (Jost's *D*) among the four *Prunus cerasifera* clusters** computed with GenAlEx v6.503.

Only the individuals assigned to a genetic cluster with a membership proportion greater than or equal to 90%, were retained. Jost's *D* values are displayed below the diagonal and the *P*-values above the diagonal. All pairwise Jost's *D* values were significant ($P<0.05$, Number of permutations = 999).

| Jost's D | Central Asia | Caucasia 1 | Caucasia 2 | Europe |
|---|---|---|---|---|
| Central Asia | 0 | 0.001 | 0.001 | 0.001 |
| Caucasia 1 | 0.196 | 0 | 0.001 | 0.001 |
| Caucasia 2 | 0.263 | 0.226 | 0 | 0.003 |
| Europe | 0.361 | 0.330 | 0.356 | 0 |
