## Supplementary figures for "Genetic diversity and population structure analyses in the Alpine plum (*Prunus brigantina* Vill.) confirm its affiliation to the Armeniaca section"

**Figure S1. DeltaK plot as a function of K for the *Prunus brigantina* (A) and Prunophora (B) dataset.**

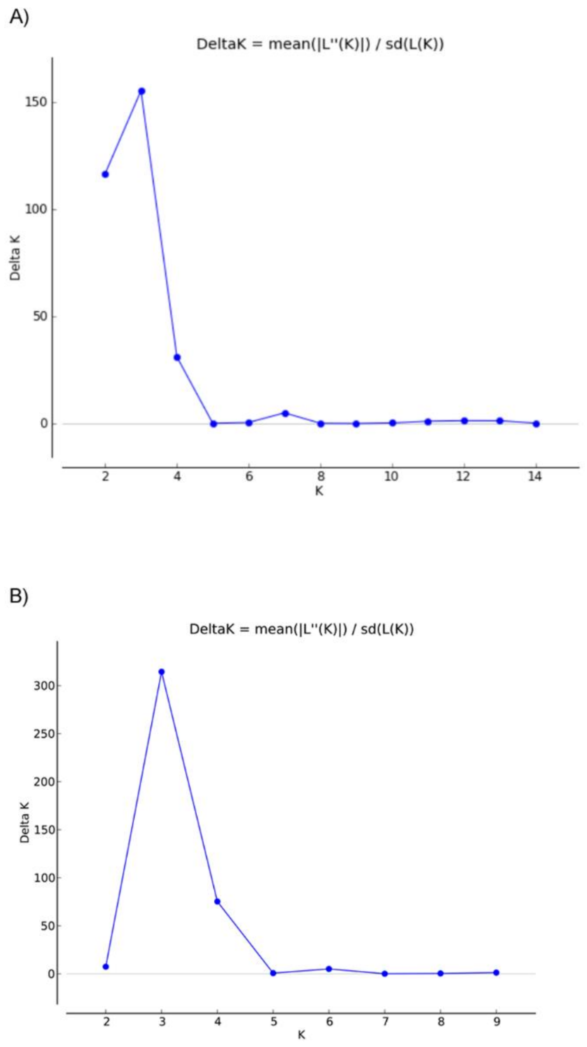

**Figure S2. Bayesian clustering on *Prunus brigantina* samples in the French Alps.**

*Prunus brigantina* dataset included 71 individuals sampled from the French Alps and two samples from the French GRC repository. Each individual is represented by a vertical bar, partitioned into *K* segments representing the inferred proportions of ancestry of its genome.

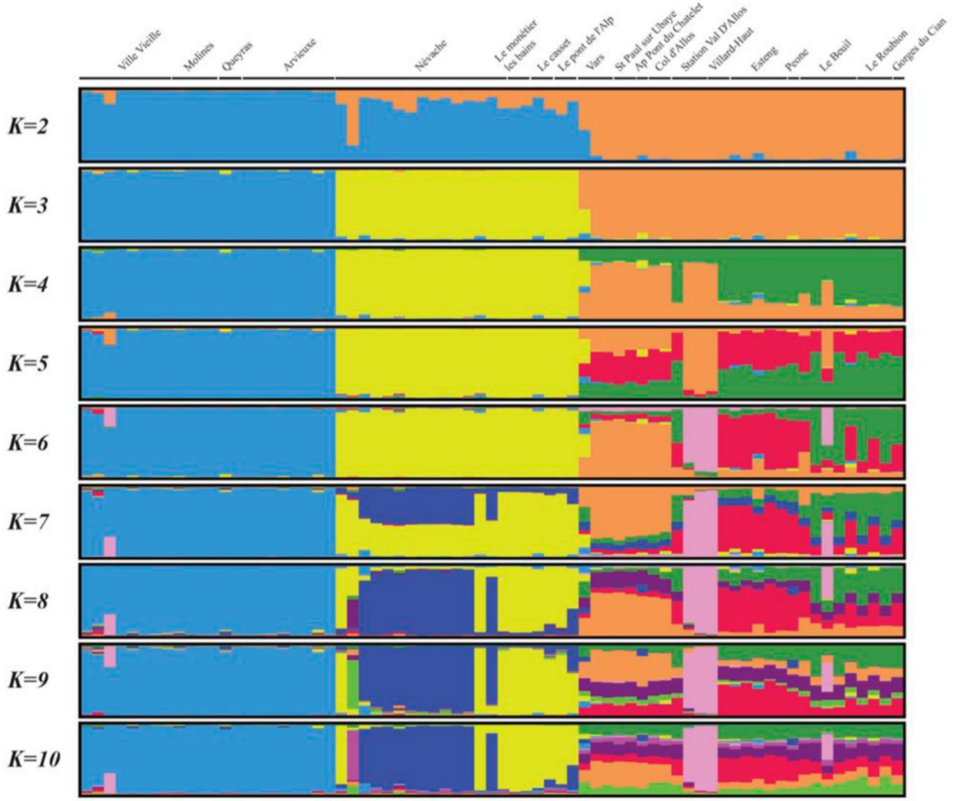

**Figure S3. Isolation by distance (IBD) test in *Prunus brigantina*.**

a. Distribution of correlation values between genetic and geographic distances under the assumption of lack of isolation by distance, drawn from permutations; the observed value of the correlation between the distance matrices, represented by the black diamond, falls within the expected distribution which indicates the lack of isolation by distance pattern.

b. Pairwise Edwards' distances plotted against Euclidean geographic distances, with local density of points plotted using a two-dimensional kernel density estimate, displayed in colour from white to red. The solid line represents the fitted linear regression between Edwards' genetic and Euclidean geographic distances.

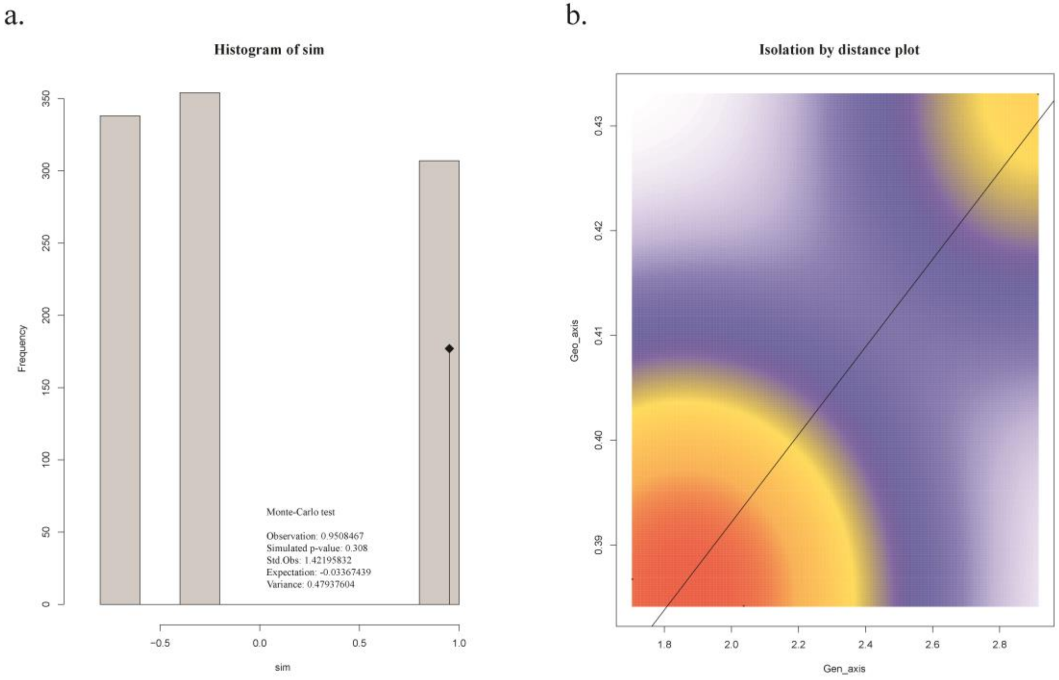

**Figure S4. Bayesian analysis on Armeniaca and wild *Prunus brigantina* accessions.**

Genetic subdivision among Armeniaca species, *P. brigantina* included, was inferred with STRUCTURE with 24 microsatellite markers. The 648 samples belong to the six Armeniaca species as follows: *P. brigantina* (*N*=73), *P. armeniaca* (European and Chinese cultivated *N*=270 and wild, *N*=204), *P. sibirica* (*N*=84), *P. mume* (*N*=9), *P. mandshurica* (*N*=8). Each individual is represented by a vertical bar, partitioned into *K* segments representing the inferred proportions of ancestry of its genome. Species and origin of the accessions are indicated on the top of the figure.

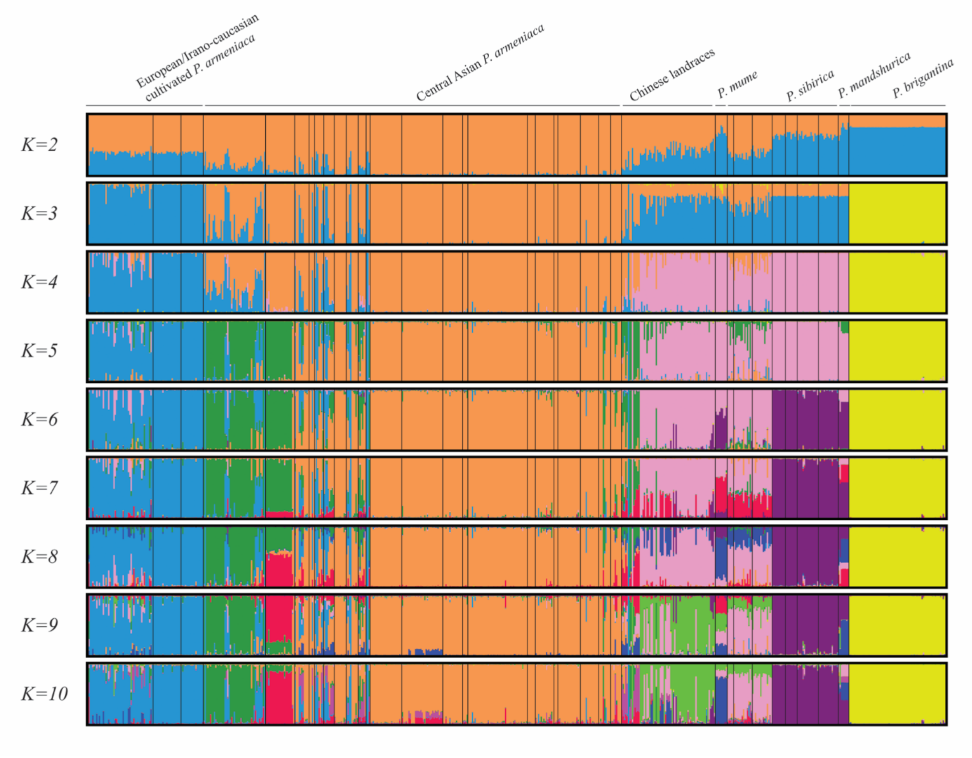

**Figure S5. Bayesian analysis on the *Prunus brigantina* dataset together with an extended Prunophora dataset.**

Genetic subdivision among Armeniaca, Prunus and Prunocerasus species was inferred with STRUCTURE with 23 microsatellite markers (supplemental information for the list of markers). The 226 samples belong to three different Prunophora species including *P. brigantina* (*N*=73), *P. cerasifera* (*N*=66), *P. armeniaca* (*N*=87), *P. salicina* (*N*=10), *P. mume* (*N*=9), *P. mexicana* (*N*=1), *P. munsoniana* (*N*=1), *P. maritima* (*N*=1), *P. americana* (*N*=1) and *P. subcordata* (*N*=1). The blue stars (*), at the bottom of the bar plots, correspond to Japanese plums (*P. salicina*) admixed with *P. cerasifera*.

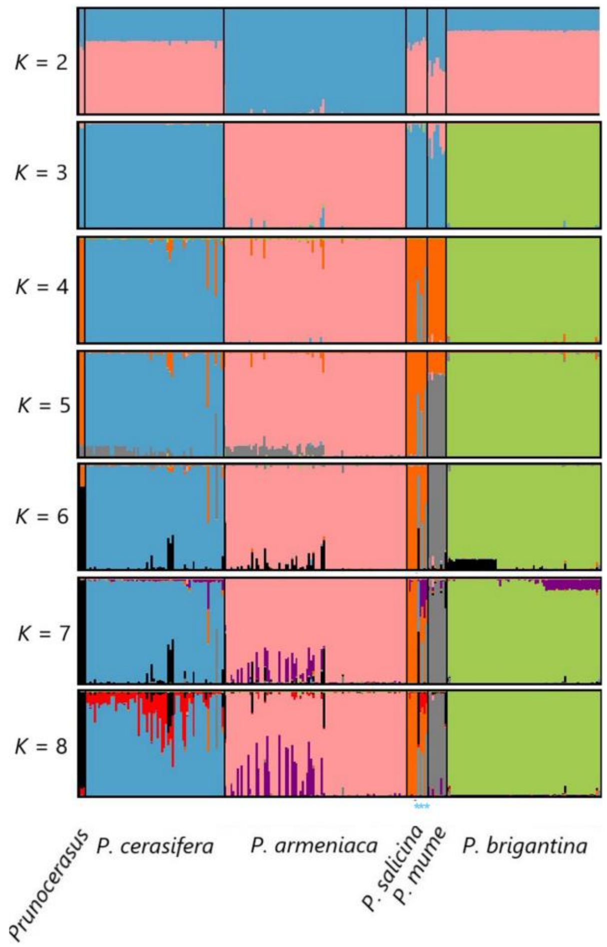
