## Supplementary Tables for "Genetic diversity and population structure analyses in the Alpine plum (*Prunus brigantina* Vill.) confirm its affiliation to the Armeniaca section"

**Table S1a. Sampling locations, geographic regions and assigned genetic cluster of *Prunus brigantina* samples in the French Alps.**

FR for an origin from the French Alps. Sampling site is indicated in GPS coordinates, N for North, E for East

| Sample code | Sampling locality | Genetic cluster | Sampling location in GPS coordinates |
|---|---|---|---|
| FR-001 | Ville Vieille | Queyras | 44°44'29N - 6°45'47E |
| FR-002 | Ville Vieille | Queyras | 44°44'27N - 6°45'43E |
| FR-003 | Ville Vieille | Queyras | 44°44'27N - 6°45'43E |
| FR-004 | Ville Vieille | Queyras | 44°44'27N - 6°45'43E |
| FR-005 | Ville Vieille | Queyras | 44°46'8N - 6°49'56E |
| FR-006 | Ville Vieille | Queyras | 44°46'8N - 6°49'56E |
| FR-007 | Ville Vieille | Queyras | 44°46'8N - 6°49'56E |
| FR-008 | Ville Vieille | Queyras | 44°46'8N - 6°49'56E |
| FR-009 | Molines | Queyras | 44°42'21N - 6°54'00E |
| FR-010 | Molines | Queyras | 44°42'21N - 6°54'04E |
| FR-011 | Molines | Queyras | 44°42'21N - 6°54'04E |
| FR-012 | Molines | Queyras | 44°42'21N - 6°54'047E |
| FR-013 | Arvieux | Queyras | 44°44'38N - 6°46'11E |
| FR-014 | Arvieux | Queyras | 44°44'27N - 6°45'43E |
| FR-015 | Arvieux | Queyras | 44°47'54N - 6°44'28E |
| FR-016 | Arvieux | Queyras | 44°47'54N - 6°44'28E |
| FR-017 | Arvieux | Queyras | 44°44'29N - 6°45'42E |
| FR-018 | Arvieux | Queyras | 44°44'27N - 6°45'43E |
| FR-019 | Arvieux | Queyras | 44°47'54N - 6°44'28E |
| FR-020 | Arvieux | Queyras | 44°47'49N - 6°44'29E |
| FR-021-A | Arvieux | Queyras | 44°44'27N - 6°45'43E |
| FR-021-B | Arvieux | Queyras | 44°44'27N - 6°45'43E |
| FR-023 | Névache | Ecrins | 44°54'3N - 6°39'4E |
| FR-024 | Névache | Ecrins | 44°54'3N - 6°39'4E |
| FR-025 | Névache | Ecrins | 44°57'34N - 6°40'28E |
| FR-026 | Névache | Ecrins | 44°57'36N - 6°40'53E |
| FR-027 | Névache | Ecrins | 44°57'37N - 6°40'30E |
| FR-028 | Névache | Ecrins | 44°57'37N - 6°40'30E |
| FR-029 | Névache | Ecrins | 44°58'30N - 6°40'32E |
| FR-030-1 | Névache | Ecrins | 45°01'48N - 6°34'46E |
| FR-030-2 | Névache | Ecrins | 44°59'11N - 6°38'48E |
| FR-031 | Névache | Ecrins | 45°01'48N - 6°34'46E |
| FR-032 | Névache | Ecrins | 45°01'48N - 6°34'46E |
| FR-033 | Névache | Ecrins | 44°57'40N - 6°40'26E |
| FR-034-1 | Névache | Ecrins | 44°56'16N - 6°34'44E |
| FR-034-2 | Névache | Ecrins | 44°01'15N - 6°35'17E |
| FR-035 | Névache | Ecrins | 44°58'5N - 6°31'37 |
| FR-036 | Le monétier les bains | Ecrins | 44°58'34N - 6°30'51E |
| FR-037 | Le monétier les bains | Ecrins | 44°58'52N - 6°29'58E |
| FR-038 | Le casset | Ecrins | 44°59'17N - 6°29'26E |
| FR-039-A | Le casset | Ecrins | 45°0'39N - 6°28'24E |
| FR-039-B | Le pont de l'Alp | Ecrins | 45°1'7N - 6°27'54E |
| FR-040-A | Le pont de l'Alp | Ecrins | 45°1'24N - 6°27'21E |
| FR-041 | Vars | Mercantour | 44°26'22N - 6°41'19E |
| FR-042 | Vars | Mercantour | 44°35'57N - 6°41'23E |
| FR-043 | Vars | Mercantour | 44°31'45N - 6°43'2E |

| | | | |
|---|---|---|---|
| FR-044 | St Paul sur Ubaye | Mercantour | 44°31'2N - 6°45'25E |
| FR-045 | St Paul sur Ubaye | Mercantour | 44°31'28N - 6°47'47E |
| FR-046 | Ap Pont du Chatelet | Mercantour | 44°31'45N - 6°47'18E |
| FR-047 | Col d'Allos | Mercantour | 44°19'18N - 6°36'6E |
| FR-048 | Col d'Allos | Mercantour | 44°19'12N - 6°35'57E |
| FR-049 | Station Val D'Allos | Mercantour | 44°16'54N - 6°34'23E |
| FR-050 | Station Val D'Allos | Mercantour | 44°16'3N - 6°34'52E |
| FR-051 | Station Val D'Allos | Mercantour | 44°14'36N - 6°38'46E |
| FR-052 | Villard-Haut | Mercantour | 44°14'43N - 6°38'58E |
| FR-053 | Villard-Haut | Mercantour | 44°9'56N - 6°42'40E |
| FR-054 | Esteng | Mercantour | 44°9'13N - 6°43'53E |
| FR-055 | Esteng | Mercantour | 44°14'5N - 6°45'8E |
| FR-056 | Esteng | Mercantour | 44°14'39N - 6-45'18E |
| FR-057 | Esteng | Mercantour | 44°14'28N - 6°45'9E |
| FR-058 | Esteng | Mercantour | 44°13'10N - 6°44'45E |
| FR-059 | Peone | Mercantour | 44°7'38N - 6°55'2E |
| FR-060 | Le Beuil | Mercantour | 44°6'7N - 6°59'11E |
| FR-061 | Le Beuil | Mercantour | 44°6'4N - 6°59'36E |
| FR-062 | Le Beuil | Mercantour | 44°6'4N - 6°59'36E |
| FR-063 | Le Beuil | Mercantour | 44°6'7N - 6°59'36E |
| FR-064 | Le Beuil | Mercantour | 44°6'9N - 6°59'45E |
| FR-065 | Le Roubion | Mercantour | 44°5'15N - 7°2'18E |
| FR-066 | Le Roubion | Mercantour | 44°5'13N - 7°2'18E |
| FR-067 | Le Roubion | Mercantour | 44°5'24N - 7°2'21E |
| FR-068 | Gorges du Cian | Mercantour | 44°1'50N - 6°58'37E |

### Table S1b. Sampling locations or germplasm repository of *Prunus cerasifera* samples and other Prunus and Prunocerasus sp

[1] as indicated by the curator of the germplasm collection where the sample is maintained or as identified *in situ*. [2] In decimal degrees. [3] Origin as indicated in the database of the germplasm repository. n/a, not applicable because admixed and thus not used in the correlation tests

| Sample code | Specie[1] | Sampling locality (country) | Sampling location in GPS coordinates[2] | | Provider (Germplasm repository) | Voucher information[3] | Coordinates used in the correlation tests[2] | |
|---|---|---|---|---|---|---|---|---|
| | | | Latitude | Longitude | | | Latitude | Longitude |
| FR-070 | *P. cerasifera* | Entraunes (Alpes-Maritimes, France) | 44.1891667 | 6.75083 | | | 44.1891667 | 6.75083 |
| X29 | *P. cerasifera* | Le Tourne (Gironde, France) | 44.710585 | -0.4045023 | | | 44.710585 | -0.4045023 |
| az117 | *P. cerasifera* | LaHic (Azerbaijan) | 40.83912167 | 48.37526167 | | | 40.83912167 | 48.37526167 |
| az118 | *P. cerasifera* | LaHic (Azerbaijan) | 40.839415 | 48.37533667 | | | 40.839415 | 48.37533667 |
| az119 | *P. cerasifera* | Lənkəran (Azerbaijan) | 38.653935 | 48.77528833 | | | 38.653935 | 48.77528833 |
| az120 | *P. cerasifera* | Lənkəran (Azerbaijan) | 38.653935 | 48.77528833 | | | 38.653935 | 48.77528833 |
| az121 | *P. cerasifera* | Lərik - Zuvand (Talysh region, Azerbaijan) | 38.68598333 | 48.38991333 | | | 38.68598333 | 48.38991333 |
| az122 | *P. cerasifera* | Mountain sheki (Sheki region, Azerbaijan) | 41.1777778 | 47.19011667 | | | n/a | n/a |
| az123 | *P. cerasifera* | Mountain sheki (Sheki region, Azerbaijan) | 41.1359361 | 47.18225833 | | | n/a | n/a |
| az124 | *P. cerasifera* | Sheki (Azerbaijan) | 41.2032861 | 47.19708611 | | | n/a | n/a |
| az125 | *P. cerasifera* | Gəcən (Azerbaijan) | 40.45901667 | 46.33495833 | | | 40.45901667 | 46.33495833 |
| az126 | *P. cerasifera* | Gəcən (Azerbaijan) | 40.43975333 | 46.32485167 | | | 40.43975333 | 46.32485167 |
| az127 | *P. cerasifera* | Sheki (Azerbaijan) | 41.2049111 | 47.19834722 | | | n/a | n/a |
| az128 | *P. cerasifera* | Genofond (Sheki region, Azerbaijan) | 41.1366611 | 47.18333889 | | | n/a | n/a |
| az129 | *P. cerasifera* | Genofond (Sheki region, Azerbaijan) | 41.1365667 | 47.18285556 | | | n/a | n/a |
| az130 | *P. cerasifera* | LaHic (Azerbaijan) | 40.8452194 | 48.38313056 | | | n/a | n/a |
| az131 | *P. cerasifera* | Lərik - Zuvand (Talysh region, Azerbaijan) | 38.686835 | 48.392005 | | | 38.686835 | 48.392005 |
| az192 | *P. cerasifera* | Lahic (Azerbaijan) | 40.83949333 | 48.37518833 | | | 40.83949333 | 48.37518833 |
| az193 | *P. cerasifera* | Lahic (Azerbaijan) | 40.79094167 | 48.31801167 | | | 40.79094167 | 48.31801167 |
| az197 | *P. cerasifera* | Badamli village (Naxçivan, Azerbaijan) | 39.4504141 | 45.530454 | | | 39.4504141 | 45.530454 |
| az239 | *P. cerasifera* | Naxçivan, Botanic Garden (Azerbaijan) | 39.2067236 | 45.408394 | | | 39.2067236 | 45.408394 |
| az24 | *P. cerasifera* | Naxçivan (Azerbaijan) | 39.3194099 | 45.5190282 | | | 39.3194099 | 45.5190282 |
| az25 | *P. cerasifera* | Naxçivan (Azerbaijan) | 39.3194099 | 45.5190282 | | | n/a | n/a |
| az48 | *P. cerasifera* | orduddad mountains, Naxçivan (Azerbaijan) | 38.940745 | 46.019205 | | | 38.940745 | 46.019205 |
| az49 | *P. cerasifera* | orduddad mountains, Naxçivan (Azerbaijan) | 38.940745 | 46.019205 | | | 38.940745 | 46.019205 |
| az51 | *P. cerasifera* | orduddad mountains, Naxçivan (Azerbaijan) | 38.9456861 | 46.02089722 | | | n/a | n/a |
| az59 | *P. cerasifera* | Guba (Azerbaijan) | 41.3614349 | 48.5137557 | | | 41.3614349 | 48.5137557 |
| KR208 | *P. cerasifera* | Arslanbob (Kyrgyzstan) | 41.30525 | 72.97438333 | | | 41.30525 | 72.97438333 |

| | | | | | | | | |
|---|---|---|---|---|---|---|---|---|
| KR209 | *P. cerasifera* | Arslanbob (Kyrgyzstan) | 41.317495 | 72.9674 | | | 41.317495 | 72.9674 |
| KR210 | *P. cerasifera* | Arslanbob (Kyrgyzstan) | 41.31742167 | 72.97399167 | | | 41.31742167 | 72.97399167 |
| KR211 | *P. cerasifera* | Arslanbob (Kyrgyzstan) | 41.31679 | 72.97526 | | | 41.31679 | 72.97526 |
| KR215 | *P. cerasifera* | Arslanbob (Kyrgyzstan) | 41.30111333 | 72.96915 | | | 41.30111333 | 72.96915 |
| KR216 | *P. cerasifera* | Arslanbob (Kyrgyzstan) | 41.30201 | 72.96727 | | | 41.30201 | 72.96727 |
| KR219 | *P. cerasifera* | Sary-Chelek (Kyrgyzstan) | 41.80082 | 71.96757167 | | | 41.80082 | 71.96757167 |
| KR225 | *P. cerasifera* | Sary-Chelek (Kyrgyzstan) | 41.85466667 | 71.968755 | | | 41.85466667 | 71.968755 |
| KR226 | *P. cerasifera* | Sary-Chelek (Kyrgyzstan) | 41.85529167 | 71.97241167 | | | 41.85529167 | 71.97241167 |
| KR227 | *P. cerasifera* | Sary-Chelek (Kyrgyzstan) | 41.85459 | 71.96683833 | | | 41.85459 | 71.96683833 |
| KR228 | *P. cerasifera* | Sary-Chelek (Kyrgyzstan) | 41.82641667 | 71.95463833 | | | 41.82641667 | 71.95463833 |
| KR230 | *P. cerasifera* | Lake Toktogul (Kyrgyzstan) | 41.75388889 | 72.92527778 | | | n/a | n/a |
| KR231 | *P. cerasifera* | Jalal-Abad (Kyrgyzstan) | 40.92445 | 72.9412 | | | n/a | n/a |
| KR232 | *P. cerasifera* | Jalal-Abad (Kyrgyzstan) | 40.9326849 | 72.9981116 | | | 40.9326849 | 72.9981116 |
| KR233 | *P. salicina* | Jalal-Abad (Kyrgyzstan) | 40.91245 | 72.951123 | | | n/a | n/a |
| KR234 | *P. cerasifera* | Jalal-Abad (Kyrgyzstan) | 40.9119444 | 72.94055556 | | | n/a | n/a |
| KR244 | *P. cerasifera* | between Arslanbob and Sary-Chelek (Kyrgyz | 41.3602778 | 72.86680556 | | | n/a | n/a |
| KR245 | *P. cerasifera* | between Arslanbob and Sary-Chelek (Kyrgyz | 41.361 | 72.866905 | | | n/a | n/a |
| KR255 | *P. cerasifera* | between Arslanbob and Sary-Chelek (Kyrgyz | 41.68809167 | 71.99074 | | | 41.68809167 | 71.99074 |
| KR256 | *P. cerasifera* | between Arslanbob and Sary-Chelek (Kyrgyz | 41.800815 | 71.95337833 | | | 41.800815 | 71.95337833 |
| kz281 | *P. cerasifera* | Medeu (Kazakhstan) | 43.1614722 | 77.05327778 | | | n/a | n/a |
| kz282 | *P. cerasifera* | Medeu (Kazakhstan) | 43.1575 | 77.0506 | | | 43.1575 | 77.0506 |
| kz284 | *P. cerasifera* | Kunterbulak (Kazakhstan) | 43.23625 | 77.09980556 | | | n/a | n/a |
| kz298 | *P. cerasifera* | Kunterbulak (Kazakhstan) | 43.2345556 | 77.08783333 | | | n/a | n/a |
| kz303 | *P. cerasifera* | Berezki garden (Kazakhstan) | 43.2884 | 77.17421667 | | | 43.2884 | 77.17421667 |
| P0016-5 | *P. cerasifera* | | | | CRB INRAE -Bourran- | St-Marcellin, Guillot nurserie | n/a | n/a |
| P0018 | *P. cerasifera* | | | | CRB INRAE -Bourran- | East Malling Research Statio | 51.28786 | 0.438343 |
| P0489 | *P. salicina* | | | | CRB INRAE -Bourran- | Louisiana (United States) / U | n/a | n/a |
| P1079 | *P. cerasifera* | | | | CRB INRAE -Bourran- | Bordeaux, INRAe Grande Fe | n/a | n/a |
| P2126 | *P. cerasifera* | | | | CRB INRAE -Bourran- | Bologna (Italy) / Italie - Pep | n/a | n/a |
| P2175 | *P. cerasifera* | | | | CRB INRAE -Bourran- | Bucharest, Horticultural and | 44.4361414 | 26.1027202 |
| P2196 | *P. cerasifera* | | | | CRB INRAE -Bourran- | Botanical Garden of Metz, tr | 48.90786 | 6.0589 |
| P2214 | *P. cerasifera* | | | | CRB INRAE -Bourran- | Nîmes (Department of Gard, | 43.8374249 | 4.3600687 |
| P2415 | *P. cerasifera* | | | | CRB INRAE -Bourran- | Saint-Genis-Laval (France) | 45.6950021 | 4.7928827 |
| P2550 | *P. cerasifera* | | | | CRB INRAE -Bourran- | Weener (Germany) / Allemag | n/a | n/a |

| | | | | | |
|---|---|---|---|---|---|
| P2646 | *P. cerasifera* | CRB INRAE -Bourran- | Fruit Tree Research Station I | 59.33258 | 18.0649 |
| P3103 | *P. cerasifera* | CRB INRAE -Bourran- | Pitesti (Romania) | n/a | n/a |
| P3188 | *P. cerasifera* | CRB INRAE -Bourran- | Krimsk (Krasnodar Krai, Rus | n/a | n/a |
| P3195 | *P. cerasifera* | CRB INRAE -Bourran- | Krimsk (Krasnodar Krai, Rus | n/a | n/a |
| P3196 | *P. cerasifera* | CRB INRAE -Bourran- | Armenia | n/a | n/a |
| P3209 | *P. cerasifera* | CRB INRAE -Bourran- | Krimsk (Krasnodar Krai, Rus | n/a | n/a |
| P3289 | *P. cerasifera* | CRB INRAE -Bourran- | Italy | n/a | n/a |
| P3676 | *P. subcordata* | CRB INRAE -Bourran- | Oregon (United States) | n/a | n/a |
| US13 | *P. salicina* | ARS-USDA repository, UC | South Africa | n/a | n/a |
| US14 | *P. mexicana* | ARS-USDA repository, UC | United States | n/a | n/a |
| US3 | *P. musoniana* | ARS-USDA repository, UC | United States | n/a | n/a |
| US63 | *P. salicina* | ARS-USDA repository, UC | Japan | n/a | n/a |
| US64 | *P. maritima* | ARS-USDA repository, UC | United States | n/a | n/a |
| US88 | *P. americana* | ARS-USDA repository, UC | Europe (Latvia) | n/a | n/a |
| US91 | *P. salicina* | ARS-USDA repository, UC | India | n/a | n/a |
| CH333 | *P. salicina* | National Germplasm Reposi | China (Changbai mountains) | n/a | n/a |
| CH334 | *P. salicina* | National Germplasm Reposi | China (Changbai mountains) | n/a | n/a |
| CH335 | *P. salicina* | National Germplasm Reposi | China (Changbai mountains) | n/a | n/a |
| CH336 | *P. salicina* | National Germplasm Reposi | China (Changbai mountains) | n/a | n/a |
| CH337 | *P. salicina* | National Germplasm Reposi | China (Changbai mountains) | n/a | n/a |

### Table S1c: List of individuals included in the different datasets of Table 1 together with the *P. brigantina* samples

[1] FR refers to France, AZ to Azerbaijan, CH to China, KR to Kyrgyzstan, KZ to Kazakhstan, OUZ to Uzbekistan, TCH to Czech republic (Lednice repository), TURC to Turkey (Malatya repository), US to USA (ARS-USDA Prunus germplasm repository). For more details, see Liu et al (2019). Accession numbers starting with A indicate apricot cultivars and with P, plum cultivars, as indicated in the French GRC database. - : maintained in germplasm repository, not collected in situ. The cross in the last four columns (dataset 1 to 4) means that this sample was used in the corresponding dataset.

| Sample [1] | Species | Type | Origin | Dataset 1 | Dataset 2 | Dataset 3 | Dataset 4 |
|---|---|---|---|---|---|---|---|
| FR-001 | *P. brigantina* | Wild | Alps, France | x | x | x | |
| FR-002 | *P. brigantina* | Wild | Alps, France | x | x | x | |
| FR-003 | *P. brigantina* | Wild | Alps, France | x | x | x | |
| FR-004 | *P. brigantina* | Wild | Alps, France | x | x | x | |
| FR-005 | *P. brigantina* | Wild | Alps, France | x | x | x | |
| FR-006 | *P. brigantina* | Wild | Alps, France | x | x | x | |
| FR-007 | *P. brigantina* | Wild | Alps, France | x | x | x | |
| FR-008 | *P. brigantina* | Wild | Alps, France | x | x | x | |
| FR-009 | *P. brigantina* | Wild | Alps, France | x | x | x | |
| FR-010 | *P. brigantina* | Wild | Alps, France | x | x | x | |
| FR-011 | *P. brigantina* | Wild | Alps, France | x | x | x | |
| FR-012 | *P. brigantina* | Wild | Alps, France | x | x | x | |
| FR-013 | *P. brigantina* | Wild | Alps, France | x | x | x | |
| FR-014 | *P. brigantina* | Wild | Alps, France | x | x | x | |
| FR-015 | *P. brigantina* | Wild | Alps, France | x | x | x | |
| FR-016 | *P. brigantina* | Wild | Alps, France | x | x | x | |
| FR-017 | *P. brigantina* | Wild | Alps, France | x | x | x | |
| FR-018 | *P. brigantina* | Wild | Alps, France | x | x | x | |
| FR-019 | *P. brigantina* | Wild | Alps, France | x | x | x | |
| FR-020 | *P. brigantina* | Wild | Alps, France | x | x | x | |
| FR-021-A | *P. brigantina* | Wild | Alps, France | x | x | x | |
| FR-021-B | *P. brigantina* | Wild | Alps, France | x | x | x | |
| FR-023 | *P. brigantina* | Wild | Alps, France | x | x | x | |
| FR-024 | *P. brigantina* | Wild | Alps, France | x | x | x | |
| FR-025 | *P. brigantina* | Wild | Alps, France | x | x | x | |
| FR-026 | *P. brigantina* | Wild | Alps, France | x | x | x | |
| FR-027 | *P. brigantina* | Wild | Alps, France | x | x | x | |
| FR-028 | *P. brigantina* | Wild | Alps, France | x | x | x | |
| FR-029 | *P. brigantina* | Wild | Alps, France | x | x | x | |
| FR-030-1 | *P. brigantina* | Wild | Alps, France | x | x | x | |
| FR-030-2 | *P. brigantina* | Wild | Alps, France | x | x | x | |
| FR-031 | *P. brigantina* | Wild | Alps, France | x | x | x | |
| FR-032 | *P. brigantina* | Wild | Alps, France | x | x | x | |
| FR-033 | *P. brigantina* | Wild | Alps, France | x | x | x | |
| FR-034-1 | *P. brigantina* | Wild | Alps, France | x | x | x | |
| FR-034-2 | *P. brigantina* | Wild | Alps, France | x | x | x | |
| FR-035 | *P. brigantina* | Wild | Alps, France | x | x | x | |
| FR-036 | *P. brigantina* | Wild | Alps, France | x | x | x | |
| FR-037 | *P. brigantina* | Wild | Alps, France | x | x | x | |
| FR-038 | *P. brigantina* | Wild | Alps, France | x | x | x | |
| FR-039-A | *P. brigantina* | Wild | Alps, France | x | x | x | |
| FR-039-B | *P. brigantina* | Wild | Alps, France | x | x | x | |
| FR-040-A | *P. brigantina* | Wild | Alps, France | x | x | x | |
| FR-041 | *P. brigantina* | Wild | Alps, France | x | x | x | |
| FR-042 | *P. brigantina* | Wild | Alps, France | x | x | x | |
| FR-043 | *P. brigantina* | Wild | Alps, France | x | x | x | |
| FR-044 | *P. brigantina* | Wild | Alps, France | x | x | x | |
| FR-045 | *P. brigantina* | Wild | Alps, France | x | x | x | |
| FR-046 | *P. brigantina* | Wild | Alps, France | x | x | x | |
| FR-047 | *P. brigantina* | Wild | Alps, France | x | x | x | |
| FR-048 | *P. brigantina* | Wild | Alps, France | x | x | x | |
| FR-049 | *P. brigantina* | Wild | Alps, France | x | x | x | |
| FR-050 | *P. brigantina* | Wild | Alps, France | x | x | x | |
| FR-051 | *P. brigantina* | Wild | Alps, France | x | x | x | |
| FR-052 | *P. brigantina* | Wild | Alps, France | x | x | x | |
| FR-053 | *P. brigantina* | Wild | Alps, France | x | x | x | |
| FR-054 | *P. brigantina* | Wild | Alps, France | x | x | x | |
| FR-055 | *P. brigantina* | Wild | Alps, France | x | x | x | |
| FR-056 | *P. brigantina* | Wild | Alps, France | x | x | x | |
| FR-057 | *P. brigantina* | Wild | Alps, France | x | x | x | |

| | | | | | | |
|---|---|---|---|---|---|---|
| FR-058 | *P. brigantina* | Wild | Alps, France | x | x | x |
| FR-059 | *P. brigantina* | Wild | Alps, France | x | x | x |
| FR-060 | *P. brigantina* | Wild | Alps, France | x | x | x |
| FR-061 | *P. brigantina* | Wild | Alps, France | x | x | x |
| FR-062 | *P. brigantina* | Wild | Alps, France | x | x | x |
| FR-063 | *P. brigantina* | Wild | Alps, France | x | x | x |
| FR-064 | *P. brigantina* | Wild | Alps, France | x | x | x |
| FR-065 | *P. brigantina* | Wild | Alps, France | x | x | x |
| FR-066 | *P. brigantina* | Wild | Alps, France | x | x | x |
| FR-067 | *P. brigantina* | Wild | Alps, France | x | x | x |
| FR-068 | *P. brigantina* | Wild | Alps, France | x | x | x |
| B5Rg1Lg7 | *P. brigantina* | - | France (French repository) | x | x | x |
| B6Rg1Lg8 | *P. brigantina* | - | France (French repository) | x | x | x |
| 3024 | *P. armeniaca* | Cultivated | Uzbekistan (province of Toshkent Shahri, Tashkent) | | x | x |
| A0804 | *P. armeniaca* | Cultivated | France | | x | x |
| A1236 | *P. armeniaca* | Cultivated | France | | x | x |
| A1314 | *P. armeniaca* | Cultivated | Spain | | x | x |
| A1693 | *P. armeniaca* | Cultivated | Ukrainia (Krimea province) | | x | x |
| A1731 | *P. armeniaca* | Cultivated | France | | x | x |
| A1809 | *P. armeniaca* | Cultivated | Greece | | x | x |
| A1939 | *P. armeniaca* | Cultivated | France | | x | x |
| A1956 | *P. armeniaca* | Cultivated | South Africa | | x | x |
| A2065 | *P. armeniaca* | Cultivated | Morocco | | x | x |
| A2069 | *P. armeniaca* | Cultivated | Morocco | | x | x |
| A2129 | *P. armeniaca* | Cultivated | France | | x | x |
| A2204 | *P. armeniaca* | Cultivated | Greece | | x | x |
| A2205 | *P. armeniaca* | Cultivated | Iran | | x | x |
| A2265 | *P. armeniaca* | Cultivated | France | | x | x |
| A2311 | *P. armeniaca* | Cultivated | France | | x | x |
| A2335 | *P. armeniaca* | Cultivated | Romania | | x | x |
| A2360 | *P. armeniaca* | Cultivated | Tunisia | | x | x |
| A2490 | *P. armeniaca* | Cultivated | France | | x | x |
| A3698VS | *P. armeniaca* | Cultivated | Slovakia | | x | x |
| A3862 | *P. armeniaca* | Cultivated | USA | | x | x |
| A4374 | *P. armeniaca* | Cultivated | Spain | | x | x |
| Bakour | *P. armeniaca* | Cultivated | Tunisia | | x | x |
| Bergeron | *P. armeniaca* | Cultivated | France | | x | x |
| Canino | *P. armeniaca* | Cultivated | Spain | | x | x |
| Currot | *P. armeniaca* | Cultivated | Spain | | x | x |
| Goldrich | *P. armeniaca* | Cultivated | USA | | x | x |
| Luizet | *P. armeniaca* | Cultivated | France | | x | x |
| Mandorla | *P. armeniaca* | Cultivated | Italy | | x | x |
| Moniqui | *P. armeniaca* | Cultivated | Spain | | x | x |
| Precoce_d_in | *P. armeniaca* | Cultivated | Italy | | x | x |
| San Castrese | *P. armeniaca* | Cultivated | Italy | | x | x |
| Stark Early O | *P. armeniaca* | Cultivated | USA | | x | x |
| Stella | *P. armeniaca* | Cultivated | USA | | x | x |
| TCH_1 | *P. armeniaca* | Cultivated | Ukrainia (Krimea province) | | x | x |
| TCH_100 | *P. armeniaca* | Cultivated | France | | x | x |
| TCH_12 | *P. armeniaca* | Cultivated | Ukrainia (Krimea province) | | x | x |
| TCH_13 | *P. armeniaca* | Cultivated | Ukrainia (Krimea province) | | x | x |
| TCH_20 | *P. armeniaca* | Cultivated | Armenia | | x | x |
| TCH_25 | *P. armeniaca* | Cultivated | Romania | | x | x |
| TCH_27 | *P. armeniaca* | Cultivated | Italy | | x | x |
| TCH_47 | *P. armeniaca* | Cultivated | Romania | | x | x |
| TCH_5 | *P. armeniaca* | Cultivated | Ukrainia | | x | x |
| TCH_67 | *P. armeniaca* | Cultivated | Hungaria | | x | x |
| TCH_84 | *P. armeniaca* | Cultivated | France | | x | x |
| TCH_97 | *P. armeniaca* | Cultivated | Bulgaria | | x | x |
| US066 | *P. armeniaca* | Cultivated | Uzbekistan | | x | x |
| US067 | *P. armeniaca* | Cultivated | Former Russian federation | | x | x |
| Velázquez | *P. armeniaca* | Cultivated | Spain | | x | x |
| A2348 | *P. armeniaca* | Cultivated | Armenia | | x | x |
| turc_04 | *P. armeniaca* | Cultivated | Malatya germplasm repository (Malatya province, Turkey) | | x | x |
| turc_27 | *P. armeniaca* | Cultivated | Adilcevaz (Bitlis province, Turkey) | | x | x |
| turc_28 | *P. armeniaca* | Cultivated | Adilcevaz (Bitlis province, Turkey) | | x | x |
| turc_29 | *P. armeniaca* | Cultivated | Adilcevaz (Bitlis province, Turkey) | | x | x |
| turc_33 | *P. armeniaca* | Cultivated | Ahlat (Bitlis province, Turkey) | | x | x |
| turc_35 | *P. armeniaca* | Cultivated | Ahlat (Bitlis province, Turkey) | | x | x |
| turc_36 | *P. armeniaca* | Cultivated | Ahlat (Bitlis province, Turkey) | | x | x |

| | | | | |
|---|---|---|---|---|
| turc_41 | *P. armeniaca* | Cultivated Arapgir (Malatya province, Turkey) | x | x |
| turc_42 | *P. armeniaca* | Cultivated Arapgir (Malatya province, Turkey) | x | x |
| turc_43 | *P. armeniaca* | Cultivated Arapgir (Malatya province, Turkey) | x | x |
| turc_44 | *P. armeniaca* | Cultivated Edremit (Van province, Turkey) | x | x |
| turc_45 | *P. armeniaca* | Cultivated Edremit (Van province, Turkey) | x | x |
| turc_46 | *P. armeniaca* | Cultivated Edremit (Van province, Turkey) | x | x |
| turc_52 | *P. armeniaca* | Cultivated Zerdali (Adana province, Turkey) | x | x |
| turc_53 | *P. armeniaca* | Cultivated Zerdali (Adana province, Turkey) | x | x |
| turc_55 | *P. armeniaca* | Cultivated Zerdali (Adana province, Turkey) | x | x |
| turc_60 | *P. armeniaca* | Cultivated Malatya germplasm repository (Malatya province, Turkey) | x | x |
| turc_68 | *P. armeniaca* | Cultivated Iskenderun surroundings (Hatay province, Turkey) | x | x |
| turc_69 | *P. armeniaca* | Cultivated Iskenderun surroundings (Hatay province, Turkey) | x | x |
| turc_70 | *P. armeniaca* | Cultivated Iskenderun surroundings (Hatay province, Turkey) | x | x |
| az_100 | *P. armeniaca* | Cultivated Ganja city (Ganja region, Azerbaijan) | x | x |
| az_111 | *P. armeniaca* | Cultivated Qobustan (Baku region, Azerbaijan) | x | x |
| az_172 | *P. armeniaca* | Cultivated Baku city (Baku region, Azerbaijan) | x | x |
| az_201_1 | *P. armeniaca* | Cultivated Badamli village (Nakhchivan, Azerbaijan) | x | x |
| az_223_5_1 | *P. armeniaca* | Cultivated Ayrinc mountains (Nakhchivan, Azerbaijan) | x | x |
| az_255_1 | *P. armeniaca* | Cultivated Ordubad (Nakhchivan, Azerbaijan) | x | x |
| az_69 | *P. armeniaca* | Cultivated Lerik (Talysh region, Azerbaijan) | x | x |
| az_71 | *P. armeniaca* | Cultivated Lerik (Talysh region, Azerbaijan) | x | x |
| az_75 | *P. armeniaca* | Cultivated Lerik (Talysh region, Azerbaijan) | x | x |
| az_76 | *P. armeniaca* | Cultivated Lerik (Talysh region, Azerbaijan) | x | x |
| az_77 | *P. armeniaca* | Cultivated Lerik (Talysh region, Azerbaijan) | x | x |
| az_80 | *P. armeniaca* | Cultivated Abseran peninsula (Baku region, Azerbaijan) | x | x |
| az_84 | *P. armeniaca* | Cultivated Sheki Genofond (Sheki region, Azerbaijan) | x | x |
| az_86 | *P. armeniaca* | Cultivated Sheki mountains (Sheki region, Azerbaijan) | x | x |
| az_89 | *P. armeniaca* | Cultivated Lahic village (Sheki region, Azerbaijan) | x | x |
| az_90 | *P. armeniaca* | Cultivated Seki mountains (Sheki, Azerbaijan) | x | x |
| az_91 | *P. armeniaca* | Cultivated Seki mountains (Sheki, Azerbaijan) | x | x |
| kz_1 | *P. armeniaca* | Cultivated Almaty Pomological garden (Almaty region, Kazakhstan) | x | |
| kz_10 | *P. armeniaca* | Cultivated Almaty Pomological garden (Almaty region, Kazakhstan) | x | |
| kz_11 | *P. armeniaca* | Cultivated Almaty Pomological garden (Almaty region, Kazakhstan) | x | |
| kz_2 | *P. armeniaca* | Cultivated Almaty Pomological garden (Almaty region, Kazakhstan) | x | |
| kz_3 | *P. armeniaca* | Cultivated Almaty Pomological garden (Almaty region, Kazakhstan) | x | |
| kz_4 | *P. armeniaca* | Cultivated Almaty Pomological garden (Almaty region, Kazakhstan) | x | |
| kz_5 | *P. armeniaca* | Cultivated Almaty Pomological garden (Almaty region, Kazakhstan) | x | |
| kz_6 | *P. armeniaca* | Cultivated Almaty Pomological garden (Almaty region, Kazakhstan) | x | |
| kz_7 | *P. armeniaca* | Cultivated Almaty Pomological garden (Almaty region, Kazakhstan) | x | |
| kz_8 | *P. armeniaca* | Cultivated Almaty Pomological garden (Almaty region, Kazakhstan) | x | |
| kz_9 | *P. armeniaca* | Cultivated Almaty Pomological garden (Almaty region, Kazakhstan) | x | |
| kz_53 | *P. armeniaca* | Cultivated Ak-kain village (Almaty region, Kazakhstan) | x | |
| kz_54 | *P. armeniaca* | Cultivated Ak-kain village (Almaty region, Kazakhstan) | x | |
| kz_55 | *P. armeniaca* | Cultivated Ak-kain village (Almaty region, Kazakhstan) | x | |
| kz_56 | *P. armeniaca* | Cultivated Ak-kain village (Almaty region, Kazakhstan) | x | |
| kz_132_1 | *P. armeniaca* | Cultivated Shymkent Dendro Park (South Kazakhstan, Kazakhstan) sampl | x | |
| kz_133_1 | *P. armeniaca* | Cultivated Shymkent Dendro Park (South Kazakhstan, Kazakhstan) sampl | x | |
| kz_134_1 | *P. armeniaca* | Cultivated Shymkent Dendro Park (South Kazakhstan, Kazakhstan) sampl | x | |
| kz_135_1 | *P. armeniaca* | Cultivated Shymkent Dendro Park (South Kazakhstan, Kazakhstan) sampl | x | |
| kz_136_1 | *P. armeniaca* | Cultivated Shymkent Dendro Park (South Kazakhstan, Kazakhstan) sampl | x | |
| kz_137_1 | *P. armeniaca* | Cultivated Shymkent Dendro Park (South Kazakhstan, Kazakhstan) sampl | x | |
| kz_138_C_1 | *P. armeniaca* | Cultivated Shymkent Dendro Park (South Kazakhstan, Kazakhstan) sampl | x | |
| kz_139_1 | *P. armeniaca* | Cultivated Sayram city surroundings (South Kazakhstan, Kazakhstan) | x | |
| kz_140_3 | *P. armeniaca* | Cultivated Sayram city surroundings (South Kazakhstan, Kazakhstan) | x | |
| kz_141_3 | *P. armeniaca* | Cultivated Sayram city surroundings (South Kazakhstan, Kazakhstan) | x | |
| kz_142_2 | *P. armeniaca* | Cultivated Sayram city surroundings (South Kazakhstan, Kazakhstan) | x | |
| kz_143_B_1 | *P. armeniaca* | Cultivated Sayram city surroundings (South Kazakhstan, Kazakhstan) | x | |
| kz_144_1 | *P. armeniaca* | Cultivated Shymkent (South Kazakhstan, Kazakhstan) | x | |
| kz_145_1 | *P. armeniaca* | Cultivated Sayram city surroundings (South Kazakhstan, Kazakhstan) | x | |
| kz_146_1 | *P. armeniaca* | Cultivated Sayram city surroundings (South Kazakhstan, Kazakhstan) | x | |
| kz_73 | *P. armeniaca* | Cultivated Turgen surroundings (Almaty province, Kazakhstan) | x | |
| kz_73A_2 | *P. armeniaca* | Cultivated Turgen surroundings (Almaty region, Kazakhstan) | x | |
| kz_73A_3 | *P. armeniaca* | Cultivated Turgen surroundings (Almaty region, Kazakhstan) | x | |
| kz_73B_10 | *P. armeniaca* | Cultivated Turgen surroundings (Almaty region, Kazakhstan) | x | |
| kz_73B_3 | *P. armeniaca* | Cultivated Turgen surroundings (Almaty region, Kazakhstan) | x | |
| kz_73B_4 | *P. armeniaca* | Cultivated Turgen surroundings (Almaty region, Kazakhstan) | x | |
| kz_73B_5 | *P. armeniaca* | Cultivated Turgen surroundings (Almaty region, Kazakhstan) | x | |
| kz_73B_8 | *P. armeniaca* | Cultivated Turgen surroundings (Almaty region, Kazakhstan) | x | |
| kz_73B_9 | *P. armeniaca* | Cultivated Turgen surroundings (Almaty region, Kazakhstan) | x | |
| kz_ma_1 | *P. armeniaca* | Cultivated Almaty green market (Almaty region, Kazakhstan) | x | |

| | | | | |
|---|---|---|---|---|
| kz_ma_10 | *P. armeniaca* | Cultivated | Almaty green market (Almaty region, Kazakhstan) | x |
| kz_ma_3 | *P. armeniaca* | Cultivated | Almaty green market (Almaty region, Kazakhstan) | x |
| kz_ma_4 | *P. armeniaca* | Cultivated | Almaty green market (Almaty region, Kazakhstan) | x |
| kz_ma_5 | *P. armeniaca* | Cultivated | Almaty green market (Almaty region, Kazakhstan) | x |
| kz_ma_6 | *P. armeniaca* | Cultivated | Almaty green market (Almaty region, Kazakhstan) | x |
| kz_ma_7 | *P. armeniaca* | Cultivated | Almaty green market (Almaty region, Kazakhstan) | x |
| kz_ma_8 | *P. armeniaca* | Cultivated | Almaty green market (Almaty region, Kazakhstan) | x |
| kz_ma_9 | *P. armeniaca* | Cultivated | Almaty green market (Almaty region, Kazakhstan) | x |
| ouz_10_1 | *P. armeniaca* | Cultivated | Bukhara old city (Bukhara region, Uzbekistan) | x |
| ouz_11_2 | *P. armeniaca* | Cultivated | Bukhara old city (Bukhara region, Uzbekistan) | x |
| ouz_15_1 | *P. armeniaca* | Cultivated | Samarkand old city (Samarkand region, Uzbekistan) | x |
| ouz_3_2 | *P. armeniaca* | Cultivated | Chimgan Beldersay (Uzbekistan) | x |
| ouz_4_3 | *P. armeniaca* | Cultivated | Chimgan Beldersay (Uzbekistan) | x |
| ouz_5_2 | *P. armeniaca* | Cultivated | Chimgan Beldersay (Uzbekistan) | x |
| US015 | *P. armeniaca* | Cultivated | Pakistan, About 12 km south of Gilgit | x |
| US016 | *P. armeniaca* | Cultivated | Pakistan, province of Azad Kashmir | x |
| US018 | *P. armeniaca* | Cultivated | Pakistan, Murtazabad (Hunza) in the Gilgit district | x |
| US019 | *P. armeniaca* | Cultivated | Pakistan, Murtazabad (Hunza) in the Gilgit district | x |
| US021 | *P. armeniaca* | Cultivated | Pakistan, Murtazabad (Hunza) in the Gilgit district | x |
| US022 | *P. armeniaca* | Cultivated | Pakistan, Murtazabad (Hunza) in the Gilgit district | x |
| US023 | *P. armeniaca* | Cultivated | Pakistan, Murtazabad (Hunza) in the Gilgit district | x |
| US024 | *P. armeniaca* | Cultivated | Pakistan, Murtazabad (Hunza) in the Gilgit district | x |
| US025 | *P. armeniaca* | Cultivated | Pakistan, Murtazabad (Hunza) in the Gilgit district | x |
| US026 | *P. armeniaca* | Cultivated | Pakistan, Kachura, Baltistan district | x |
| US027 | *P. armeniaca* | Cultivated | Pakistan, Barchu (Keris Area), Baltistan district | x |
| US028 | *P. armeniaca* | Cultivated | Pakistan, Barchu (Keris Area), Baltistan district | x |
| US030 | *P. armeniaca* | Cultivated | Pakistan, Near Rahimabad, north of Gilgit | x |
| US031 | *P. armeniaca* | Cultivated | Pakistan, Shiskat (Hunza), Gilgit district | x |
| US033 | *P. armeniaca* | Cultivated | Pakistan | x |
| US034 | *P. armeniaca* | Cultivated | Pakistan, Gulmit (Hunza), Gilgit district | x |
| US097 | *P. armeniaca* | Cultivated | India | x |
| US101 | *P. armeniaca* | Cultivated | Turkmenistan | x |
| US196 | *P. armeniaca* | Cultivated | Uzbekistan, Turkistan | x |
| US197 | *P. armeniaca* | Cultivated | Afghanistan, Province of Parwān | x |
| US199 | *P. armeniaca* | Cultivated | Pakistan, Near Hanuchal | x |
| US201 | *P. armeniaca* | Cultivated | Pakistan, Just west of Nasirabad (Hunza), Gilgit district | x |
| US204 | *P. armeniaca* | Cultivated | Pakistan, Near Gulmit (Nagar) | x |
| US206 | *P. armeniaca* | Cultivated | Pakistan, Near Gulmit (Nagar) | x |
| US207 | *P. armeniaca* | Cultivated | Pakistan, Murtazabad (Hunza) in the Gilgit district | x |
| US208 | *P. armeniaca* | Cultivated | Pakistan, Murtazabad (Hunza) in the Gilgit district | x |
| US210 | *P. armeniaca* | Cultivated | Pakistan, Murtazabad (Hunza) in the Gilgit district | x |
| US211 | *P. armeniaca* | Cultivated | Pakistan, Murtazabad (Hunza) in the Gilgit district | x |
| US212 | *P. armeniaca* | Cultivated | Pakistan, Murtazabad (Hunza) in the Gilgit district | x |
| US213 | *P. armeniaca* | Cultivated | Pakistan, Murtazabad (Hunza) in the Gilgit district | x |
| US214 | *P. armeniaca* | Cultivated | Pakistan, Garbidas, in the Baltistan district | x |
| US215 | *P. armeniaca* | Cultivated | Pakistan, Sordas, Baltistan district | x |
| US216 | *P. armeniaca* | Cultivated | Pakistan, Kothung in the Baltistan district | x |
| US217 | *P. armeniaca* | Cultivated | Pakistan, Kothung, in the Baltistan district | x |
| US218 | *P. armeniaca* | Cultivated | Pakistan, Chappi, Baltistan district | x |
| US219 | *P. armeniaca* | Cultivated | Pakistan, Kinapa, Baltistan district | x |
| US220 | *P. armeniaca* | Cultivated | Pakistan, Guram Chasma, Chitral district | x |
| US221 | *P. armeniaca* | Cultivated | Pakistan, Gulmit (Hunza), Gilgit district | x |
| US222 | *P. armeniaca* | Cultivated | Pakistan, Kagan, Hazara district, NWFD province | x |
| US229 | *P. armeniaca* | Cultivated | Afghanistan (No passport information given, Donated to NCGF | x |
| US230 | *P. armeniaca* | Cultivated | Turkmenistan | x |
| US231 | *P. armeniaca* | Cultivated | Turkmenistan (Donated to NCGR, Davis | x |
| US235 | *P. armeniaca* | Cultivated | Turkmenistan (Donated to NCGR, Davis) | x |
| US236 | *P. armeniaca* | Cultivated | Turkmenistan (Donated to NCGR, Davis) | x |
| US237 | *P. armeniaca* | Cultivated | Turkmenistan (Donated to NCGR, Davis) | x |
| US239 | *P. armeniaca* | Cultivated | Turkmenistan (Donated to NCGR, Davis) | x |
| US241 | *P. armeniaca* | Cultivated | Turkmenistan | x |
| KR28 | *P. armeniaca* | Cultivated | Arslanbob village (Jalal-Abad Province, Kyrgyzstan) | x |
| KR29 | *P. armeniaca* | Cultivated | Arslanbob village (Jalal-Abad Province, Kyrgyzstan) | x |
| KR37 | *P. armeniaca* | Cultivated | Arslanbob village (Jalal-Abad Province, Kyrgyzstan) | x |
| CH_100 | *P. armeniaca* | Cultivated | China, Hubei province | x |
| CH_113 | *P. armeniaca* | Cultivated | China, Turpan, Xinjiang province | x |
| CH_114 | *P. armeniaca* | Cultivated | China, Turpan, Xinjiang province | x |
| CH_117 | *P. armeniaca* | Cultivated | China, Luntai, Xinjiang province | x |
| CH_119 | *P. armeniaca* | Cultivated | China, Luntai, Xinjiang province | x |
| CH_120 | *P. armeniaca* | Cultivated | China, Luntai, Xinjiang province | x |

| | | | | |
|---|---|---|---|---|
| CH_121 | *P. armeniaca* | Cultivated | China, Luntai, Xinjiang province | x |
| CH_136 | *P. armeniaca* | Cultivated | China, Ili valley, Xinjiang province | x |
| CH_137 | *P. armeniaca* | Cultivated | China, Ili valley, Xinjiang province | x |
| CH_38 | *P. armeniaca* | Cultivated | Liaoning province, China | x |
| CH_44 | *P. armeniaca* | Cultivated | Heilongjiang province, China | x |
| CH_45 | *P. armeniaca* | Cultivated | Inner Mongolia province, China | x |
| CH_46 | *P. armeniaca* | Cultivated | Hebei province, China | x |
| CH_80 | *P. armeniaca* | Cultivated | Beijing province, China | x |
| CH_155 | *P. armeniaca* | Cultivated | Beijing province, China | x |
| CH_156 | *P. armeniaca* | Cultivated | Beijing province, China | x |
| CH_157 | *P. armeniaca* | Cultivated | Beijing province, China | x |
| CH_158 | *P. armeniaca* | Cultivated | Beijing province, China | x |
| CH_159 | *P. armeniaca* | Cultivated | Beijing province, China | x |
| CH_161 | *P. armeniaca* | Cultivated | Gansu province, China | x |
| CH_162 | *P. armeniaca* | Cultivated | Gansu province, China | x |
| CH_163 | *P. armeniaca* | Cultivated | Gansu province, China | x |
| CH_164 | *P. armeniaca* | Cultivated | Guizhou province, China | x |
| CH_165 | *P. armeniaca* | Cultivated | Guizhou province, China | x |
| CH_166 | *P. armeniaca* | Cultivated | Guizhou province, China | x |
| CH_167 | *P. armeniaca* | Cultivated | Guizhou province, China | x |
| CH_168 | *P. armeniaca* | Cultivated | Guizhou province, China | x |
| CH_169 | *P. armeniaca* | Cultivated | Hebei province, China | x |
| CH_170 | *P. armeniaca* | Cultivated | Hebei province, China | x |
| CH_171 | *P. armeniaca* | Cultivated | Hebei province, China | x |
| CH_172 | *P. armeniaca* | Cultivated | Hebei province, China | x |
| CH_173 | *P. armeniaca* | Cultivated | Hebei province, China | x |
| CH_174 | *P. armeniaca* | Cultivated | Hebei province, China | x |
| CH_176 | *P. armeniaca* | Cultivated | Hebei province, China | x |
| CH_177 | *P. armeniaca* | Cultivated | He'nan province, China | x |
| CH_178 | *P. armeniaca* | Cultivated | He'nan province, China | x |
| CH_179 | *P. armeniaca* | Cultivated | Heilongjiang province, China | x |
| CH_181 | *P. armeniaca* | Cultivated | Heilongjiang province, China | x |
| CH_184 | *P. armeniaca* | Cultivated | Liaoning province, China | x |
| CH_185 | *P. armeniaca* | Cultivated | Liaoning province, China | x |
| CH_186 | *P. armeniaca* | Cultivated | Liaoning province, China | x |
| CH_187 | *P. armeniaca* | Cultivated | Liaoning province, China | x |
| CH_188 | *P. armeniaca* | Cultivated | Liaoning province, China | x |
| CH_190 | *P. armeniaca* | Cultivated | Inner Mongolia province, China | x |
| CH_191 | *P. armeniaca* | Cultivated | Inner Mongolia province, China | x |
| CH_192 | *P. armeniaca* | Cultivated | Inner Mongolia province, China | x |
| CH_193 | *P. armeniaca* | Cultivated | Inner Ningxia province, China | x |
| CH_195 | *P. armeniaca* | Cultivated | Inner Ningxia province, China | x |
| CH_196 | *P. armeniaca* | Cultivated | Shandong province, China | x |
| CH_197 | *P. armeniaca* | Cultivated | Shandong province, China | x |
| CH_198 | *P. armeniaca* | Cultivated | Shandong province, China | x |
| CH_199 | *P. armeniaca* | Cultivated | Shandong province, China | x |
| CH_200 | *P. armeniaca* | Cultivated | Shandong province, China | x |
| CH_201 | *P. armeniaca* | Cultivated | Shaanxi province, China | x |
| CH_202 | *P. armeniaca* | Cultivated | Shaanxi province, China | x |
| CH_203 | *P. armeniaca* | Cultivated | Shaanxi province, China | x |
| CH_206 | *P. armeniaca* | Cultivated | Sichuan province, China | x |
| CH_207 | *P. armeniaca* | Cultivated | Sichuan province, China | x |
| CH_208 | *P. armeniaca* | Cultivated | Sichuan province, China | x |
| CH_209 | *P. armeniaca* | Cultivated | Sichuan province, China | x |
| CH_210 | *P. armeniaca* | Cultivated | Sichuan province, China | x |
| CH_211 | *P. armeniaca* | Cultivated | Xinjiang province, China | x |
| CH_213 | *P. armeniaca* | Cultivated | Xinjiang province, China | x |
| CH_214 | *P. armeniaca* | Cultivated | Xinjiang province, China | x |
| CH_215 | *P. armeniaca* | Cultivated | Xinjiang province, China | x |
| CH_216 | *P. armeniaca* | Cultivated | Xinjiang province, China | x |
| CH_217 | *P. armeniaca* | Cultivated | Xinjiang province, China | x |
| CH_219 | *P. armeniaca* | Cultivated | Yunnan province, China | x |
| CH_222 | *P. armeniaca* | Cultivated | Zhejiang province, China | x |
| CH_223 | *P. armeniaca* | Cultivated | Zhejiang province, China | x |
| CH_224 | *P. armeniaca* | Cultivated | Zhejiang province, China | x |
| CH_122 | *P. armeniaca* | Wild | Mohu'er Arele (Xinjiang province, China) | x |
| CH_123 | *P. armeniaca* | Wild | Mohu'er Arele (Xinjiang province, China) | x |
| CH_124 | *P. armeniaca* | Wild | Mohu'er Arele (Xinjiang province, China) | x |
| CH_127 | *P. armeniaca* | Wild | Luntai germplasm repository (Xinjiang province, China) | x |
| CH_128 | *P. armeniaca* | Wild | Xinjuan mountains (Xinjiang province, China) | x |

| | | | | |
|---|---|---|---|---|
| CH_129 | *P. armeniaca* | Wild | Xinjuan mountains (Xinjiang province, China) | x |
| CH_130 | *P. armeniaca* | Wild | Xinjuan mountains (Xinjiang province, China) | x |
| CH_131 | *P. armeniaca* | Wild | Xinjuan mountains (Xinjiang province, China) | x |
| kz_12 | *P. armeniaca* | Wild | Baybulak (Almaty region, Kazakhstan) | x |
| kz_14 | *P. armeniaca* | Wild | Baybulak (Almaty region, Kazakhstan) | x |
| kz_16 | *P. armeniaca* | Wild | Baybulak (Almaty region, Kazakhstan) | x |
| kz_18 | *P. armeniaca* | Wild | Baybulak (Almaty region, Kazakhstan) | x |
| kz_20 | *P. armeniaca* | Wild | Baybulak (Almaty region, Kazakhstan) | x |
| kz_22 | *P. armeniaca* | Wild | Baybulak (Almaty region, Kazakhstan) | x |
| kz_24 | *P. armeniaca* | Wild | Baybulak (Almaty region, Kazakhstan) | x |
| kz254 | *P. armeniaca* | Wild | Baybulak (Almaty region, Kazakhstan) | x |
| kz255 | *P. armeniaca* | Wild | Baybulak (Almaty region, Kazakhstan) | x |
| kz257 | *P. armeniaca* | Wild | Baybulak (Almaty region, Kazakhstan) | x |
| kz258 | *P. armeniaca* | Wild | Baybulak (Almaty region, Kazakhstan) | x |
| kz260 | *P. armeniaca* | Wild | Baybulak (Almaty region, Kazakhstan) | x |
| kz261 | *P. armeniaca* | Wild | Baybulak (Almaty region, Kazakhstan) | x |
| kz264 | *P. armeniaca* | Wild | Baybulak (Almaty region, Kazakhstan) | x |
| kz266 | *P. armeniaca* | Wild | Baybulak (Almaty region, Kazakhstan) | x |
| kz268 | *P. armeniaca* | Wild | Baybulak (Almaty region, Kazakhstan) | x |
| kz269 | *P. armeniaca* | Wild | Baybulak (Almaty region, Kazakhstan) | x |
| kz270 | *P. armeniaca* | Wild | Baybulak (Almaty region, Kazakhstan) | x |
| kz272 | *P. armeniaca* | Wild | Baybulak (Almaty region, Kazakhstan) | x |
| kz274 | *P. armeniaca* | Wild | Baybulak (Almaty region, Kazakhstan) | x |
| kz275 | *P. armeniaca* | Wild | Baybulak (Almaty region, Kazakhstan) | x |
| kz276 | *P. armeniaca* | Wild | Baybulak (Almaty region, Kazakhstan) | x |
| kz277 | *P. armeniaca* | Wild | Baybulak (Almaty region, Kazakhstan) | x |
| kz278 | *P. armeniaca* | Wild | Baybulak (Almaty region, Kazakhstan) | x |
| kz_27 | *P. armeniaca* | Wild | Esik Lake (Almaty region, Kazakhstan) | x |
| kz_29 | *P. armeniaca* | Wild | Esik Lake (Almaty region, Kazakhstan) | x |
| kz_31 | *P. armeniaca* | Wild | Esik Lake (Almaty region, Kazakhstan) | x |
| kz_33 | *P. armeniaca* | Wild | Esik Lake (Almaty region, Kazakhstan) | x |
| kz_35 | *P. armeniaca* | Wild | Esik Lake (Almaty region, Kazakhstan) | x |
| kz_37 | *P. armeniaca* | Wild | Esik Lake (Almaty region, Kazakhstan) | x |
| kz_39 | *P. armeniaca* | Wild | Esik Lake (Almaty region, Kazakhstan) | x |
| kz_40 | *P. armeniaca* | Wild | Esik Lake (Almaty region, Kazakhstan) | x |
| kz218 | *P. armeniaca* | Wild | Esik Lake (Almaty region, Kazakhstan) | x |
| kz219 | *P. armeniaca* | Wild | Esik Lake (Almaty region, Kazakhstan) | x |
| kz221 | *P. armeniaca* | Wild | Esik Lake (Almaty region, Kazakhstan) | x |
| kz222 | *P. armeniaca* | Wild | Esik Lake (Almaty region, Kazakhstan) | x |
| kz223 | *P. armeniaca* | Wild | Esik Lake (Almaty region, Kazakhstan) | x |
| kz224 | *P. armeniaca* | Wild | Esik Lake (Almaty region, Kazakhstan) | x |
| kz226 | *P. armeniaca* | Wild | Esik Lake (Almaty region, Kazakhstan) | x |
| kz230 | *P. armeniaca* | Wild | Esik Lake (Almaty region, Kazakhstan) | x |
| kz231 | *P. armeniaca* | Wild | Esik Lake (Almaty region, Kazakhstan) | x |
| kz232 | *P. armeniaca* | Wild | Esik Lake (Almaty region, Kazakhstan) | x |
| kz233 | *P. armeniaca* | Wild | Esik Lake (Almaty region, Kazakhstan) | x |
| kz235 | *P. armeniaca* | Wild | Esik Lake (Almaty region, Kazakhstan) | x |
| kz236 | *P. armeniaca* | Wild | Esik Lake (Almaty region, Kazakhstan) | x |
| kz237 | *P. armeniaca* | Wild | Esik Lake (Almaty region, Kazakhstan) | x |
| kz238 | *P. armeniaca* | Wild | Esik Lake (Almaty region, Kazakhstan) | x |
| kz240 | *P. armeniaca* | Wild | Esik Lake (Almaty region, Kazakhstan) | x |
| kz241 | *P. armeniaca* | Wild | Esik Lake (Almaty region, Kazakhstan) | x |
| kz242 | *P. armeniaca* | Wild | Esik Lake (Almaty region, Kazakhstan) | x |
| kz243 | *P. armeniaca* | Wild | Esik Lake (Almaty region, Kazakhstan) | x |
| kz244 | *P. armeniaca* | Wild | Esik Lake (Almaty region, Kazakhstan) | x |
| kz245 | *P. armeniaca* | Wild | Esik Lake (Almaty region, Kazakhstan) | x |
| kz246 | *P. armeniaca* | Wild | Esik Lake (Almaty region, Kazakhstan) | x |
| kz247 | *P. armeniaca* | Wild | Esik Lake (Almaty region, Kazakhstan) | x |
| kz_66 | *P. armeniaca* | Wild | Medeu valley (Almaty region, Kazakhstan) | x |
| kz_71 | *P. armeniaca* | Wild | Medeu valley (Almaty region, Kazakhstan) | x |
| kz156 | *P. armeniaca* | Wild | Medeu valley (Almaty region, Kazakhstan) | x |
| kz157 | *P. armeniaca* | Wild | Medeu valley (Almaty region, Kazakhstan) | x |
| kz158 | *P. armeniaca* | Wild | Medeu valley (Almaty region, Kazakhstan) | x |
| kz160 | *P. armeniaca* | Wild | Medeu valley (Almaty region, Kazakhstan) | x |
| kz161 | *P. armeniaca* | Wild | Medeu valley (Almaty region, Kazakhstan) | x |
| kz162 | *P. armeniaca* | Wild | Medeu valley (Almaty region, Kazakhstan) | x |
| kz163 | *P. armeniaca* | Wild | Medeu valley (Almaty region, Kazakhstan) | x |
| kz164 | *P. armeniaca* | Wild | Medeu valley (Almaty region, Kazakhstan) | x |
| kz165 | *P. armeniaca* | Wild | Medeu valley (Almaty region, Kazakhstan) | x |
| kz166 | *P. armeniaca* | Wild | Medeu valley (Almaty region, Kazakhstan) | x |

| | | | | |
|---|---|---|---|---|
| kz167 | *P. armeniaca* | Wild | Medeu valley (Almaty region, Kazakhstan) | x |
| kz168 | *P. armeniaca* | Wild | Medeu valley (Almaty region, Kazakhstan) | x |
| kz169 | *P. armeniaca* | Wild | Medeu valley (Almaty region, Kazakhstan) | x |
| kz_74 | *P. armeniaca* | Wild | Turgen valley (Almaty region, Kazakhstan) | x |
| kz_77 | *P. armeniaca* | Wild | Turgen valley (Almaty region, Kazakhstan) | x |
| kz_79 | *P. armeniaca* | Wild | Turgen valley (Almaty region, Kazakhstan) | x |
| kz_82 | *P. armeniaca* | Wild | Turgen valley (Almaty region, Kazakhstan) | x |
| kz_100 | *P. armeniaca* | Wild | Big Almaty lake (Almaty region, Kazakhstan) | x |
| kz_101 | *P. armeniaca* | Wild | Big Almaty lake (Almaty region, Kazakhstan) | x |
| kz_104 | *P. armeniaca* | Wild | Big Almaty lake (Almaty region, Kazakhstan) | x |
| kz_122_1 | *P. armeniaca* | Wild | Big Almaty lake (Almaty region, Kazakhstan) | x |
| kz_46 | *P. armeniaca* | Wild | Big Almaty lake (Almaty region, Kazakhstan) | x |
| kz_51 | *P. armeniaca* | Wild | Big Almaty lake (Almaty region, Kazakhstan) | x |
| kz_98 | *P. armeniaca* | Wild | Big Almaty lake (Almaty region, Kazakhstan) | x |
| kz175 | *P. armeniaca* | Wild | Big Almaty lake (Almaty region, Kazakhstan) | x |
| kz176 | *P. armeniaca* | Wild | Big Almaty lake (Almaty region, Kazakhstan) | x |
| kz177 | *P. armeniaca* | Wild | Big Almaty lake (Almaty region, Kazakhstan) | x |
| kz178 | *P. armeniaca* | Wild | Big Almaty lake (Almaty region, Kazakhstan) | x |
| kz179 | *P. armeniaca* | Wild | Big Almaty lake (Almaty region, Kazakhstan) | x |
| kz180 | *P. armeniaca* | Wild | Big Almaty lake (Almaty region, Kazakhstan) | x |
| kz181 | *P. armeniaca* | Wild | Big Almaty lake (Almaty region, Kazakhstan) | x |
| kz184 | *P. armeniaca* | Wild | Big Almaty lake (Almaty region, Kazakhstan) | x |
| kz185 | *P. armeniaca* | Wild | Big Almaty lake (Almaty region, Kazakhstan) | x |
| kz186 | *P. armeniaca* | Wild | Big Almaty lake (Almaty region, Kazakhstan) | x |
| kz187 | *P. armeniaca* | Wild | Big Almaty lake (Almaty region, Kazakhstan) | x |
| kz188 | *P. armeniaca* | Wild | Big Almaty lake (Almaty region, Kazakhstan) | x |
| kz189 | *P. armeniaca* | Wild | Big Almaty lake (Almaty region, Kazakhstan) | x |
| kz190 | *P. armeniaca* | Wild | Big Almaty lake (Almaty region, Kazakhstan) | x |
| kz192 | *P. armeniaca* | Wild | Big Almaty lake (Almaty region, Kazakhstan) | x |
| kz193 | *P. armeniaca* | Wild | Big Almaty lake (Almaty region, Kazakhstan) | x |
| kz194 | *P. armeniaca* | Wild | Big Almaty lake (Almaty region, Kazakhstan) | x |
| kz195 | *P. armeniaca* | Wild | Big Almaty lake (Almaty region, Kazakhstan) | x |
| kz196 | *P. armeniaca* | Wild | Big Almaty lake (Almaty region, Kazakhstan) | x |
| kz197 | *P. armeniaca* | Wild | Big Almaty lake (Almaty region, Kazakhstan) | x |
| kz198 | *P. armeniaca* | Wild | Big Almaty lake (Almaty region, Kazakhstan) | x |
| kz199 | *P. armeniaca* | Wild | Big Almaty lake (Almaty region, Kazakhstan) | x |
| kz200 | *P. armeniaca* | Wild | Big Almaty lake (Almaty region, Kazakhstan) | x |
| kz201 | *P. armeniaca* | Wild | Big Almaty lake (Almaty region, Kazakhstan) | x |
| kz202 | *P. armeniaca* | Wild | Big Almaty lake (Almaty region, Kazakhstan) | x |
| kz203 | *P. armeniaca* | Wild | Big Almaty lake (Almaty region, Kazakhstan) | x |
| kz204 | *P. armeniaca* | Wild | Big Almaty lake (Almaty region, Kazakhstan) | x |
| kz205 | *P. armeniaca* | Wild | Big Almaty lake (Almaty region, Kazakhstan) | x |
| kz206 | *P. armeniaca* | Wild | Big Almaty lake (Almaty region, Kazakhstan) | x |
| kz207 | *P. armeniaca* | Wild | Big Almaty lake (Almaty region, Kazakhstan) | x |
| kz208 | *P. armeniaca* | Wild | Big Almaty lake (Almaty region, Kazakhstan) | x |
| kz209 | *P. armeniaca* | Wild | Big Almaty lake (Almaty region, Kazakhstan) | x |
| kz210 | *P. armeniaca* | Wild | Big Almaty lake (Almaty region, Kazakhstan) | x |
| kz211 | *P. armeniaca* | Wild | Big Almaty lake (Almaty region, Kazakhstan) | x |
| kz212 | *P. armeniaca* | Wild | Big Almaty lake (Almaty region, Kazakhstan) | x |
| kz213 | *P. armeniaca* | Wild | Big Almaty lake (Almaty region, Kazakhstan) | x |
| kz215 | *P. armeniaca* | Wild | Big Almaty lake (Almaty region, Kazakhstan) | x |
| kz216 | *P. armeniaca* | Wild | Big Almaty lake (Almaty region, Kazakhstan) | x |
| kz_147_1 | *P. armeniaca* | Wild | Aksu-Zhabagly (South Kazakhstan, Kazakhstan) | x |
| kz_148_1 | *P. armeniaca* | Wild | Aksu-Zhabagly (South Kazakhstan, Kazakhstan) | x |
| kz_149_1 | *P. armeniaca* | Wild | Aksu-Zhabagly (South Kazakhstan, Kazakhstan) | x |
| kz_150_1 | *P. armeniaca* | Wild | Aksu-Zhabagly (South Kazakhstan, Kazakhstan) | x |
| kz_152_1 | *P. armeniaca* | Wild | Aksu-Zhabagly (South Kazakhstan, Kazakhstan) | x |
| kz_154_1 | *P. armeniaca* | Wild | Aksu-Zhabagly (South Kazakhstan, Kazakhstan) | x |
| KR143 | *P. armeniaca* | Wild | Issyk Kul lake surroundings - Urukty river valley (Issyk Kul reg | x |
| KR144 | *P. armeniaca* | Wild | Issyk Kul lake surroundings - Urukty river valley (Issyk Kul reg | x |
| KR146 | *P. armeniaca* | Wild | Issyk Kul lake surroundings - Urukty river valley (Issyk Kul reg | x |
| KR147 | *P. armeniaca* | Wild | Issyk Kul lake surroundings - Urukty river valley (Issyk Kul reg | x |
| KR148 | *P. armeniaca* | Wild | Issyk Kul lake surroundings - Urukty river valley (Issyk Kul reg | x |
| KR149 | *P. armeniaca* | Wild | Issyk Kul lake surroundings - Urukty river valley (Issyk Kul reg | x |
| KR150 | *P. armeniaca* | Wild | Issyk Kul lake surroundings - Urukty river valley (Issyk Kul reg | x |
| KR151 | *P. armeniaca* | Wild | Issyk Kul lake surroundings - Urukty river valley (Issyk Kul reg | x |
| KR152 | *P. armeniaca* | Wild | Issyk Kul lake surroundings - Urukty river valley (Issyk Kul reg | x |
| KR153 | *P. armeniaca* | Wild | Issyk Kul lake surroundings - Urukty river valley (Issyk Kul reg | x |
| KR154 | *P. armeniaca* | Wild | Issyk Kul lake surroundings - Urukty river valley (Issyk Kul reg | x |
| KR155 | *P. armeniaca* | Wild | Issyk Kul lake surroundings - Urukty river valley (Issyk Kul reg | x |

| | | | | |
|---|---|---|---|---|
| KR156 | *P. armeniaca* | Wild | Issyk Kul lake surroundings - Urukty river valley (Issyk Kul reg | x |
| KR158 | *P. armeniaca* | Wild | Issyk Kul lake surroundings - Urukty river valley (Issyk Kul reg | x |
| KR159 | *P. armeniaca* | Wild | Ananyevo village (Issyk Kul region, Kirgizstan) | x |
| KR160 | *P. armeniaca* | Wild | Ananyevo village (Issyk Kul region, Kirgizstan) | x |
| KR161 | *P. armeniaca* | Wild | Ananyevo village (Issyk Kul region, Kirgizstan) | x |
| KR162 | *P. armeniaca* | Wild | Issyk Kul lake surroundings - Orto Byrosun river valley (Issyk I | x |
| KR163 | *P. armeniaca* | Wild | Issyk Kul lake surroundings - Orto Byrosun river valley (Issyk I | x |
| KR165 | *P. armeniaca* | Wild | Issyk Kul lake surroundings - Orto Byrosun river valley (Issyk I | x |
| KR166 | *P. armeniaca* | Wild | Issyk Kul lake surroundings - Orto Byrosun river valley (Issyk I | x |
| KR167 | *P. armeniaca* | Wild | Issyk Kul lake surroundings - Orto Byrosun river valley (Issyk I | x |
| KR168 | *P. armeniaca* | Wild | Issyk Kul lake surroundings - Orto Byrosun river valley (Issyk I | x |
| KR169 | *P. armeniaca* | Wild | Issyk Kul lake surroundings - Orto Byrosun river valley (Issyk I | x |
| KR170 | *P. armeniaca* | Wild | Issyk Kul lake surroundings - Orto Byrosun river valley (Issyk I | x |
| KR171 | *P. armeniaca* | Wild | Issyk Kul lake surroundings - Orto Byrosun river valley (Issyk I | x |
| KR172 | *P. armeniaca* | Wild | Issyk Kul lake surroundings - Orto Byrosun river valley (Issyk I | x |
| KR173 | *P. armeniaca* | Wild | Issyk Kul lake surroundings - Orto Byrosun river valley (Issyk I | x |
| KR174 | *P. armeniaca* | Wild | Issyk Kul lake surroundings - Orto Byrosun river valley (Issyk I | x |
| KR175 | *P. armeniaca* | Wild | Issyk Kul lake surroundings - Orto Byrosun river valley (Issyk I | x |
| KR176 | *P. armeniaca* | Wild | Issyk Kul lake surroundings - Orto Byrosun river valley (Issyk I | x |
| KR177 | *P. armeniaca* | Wild | Issyk Kul lake surroundings - Orto Byrosun river valley (Issyk I | x |
| KR178 | *P. armeniaca* | Wild | Issyk Kul lake surroundings - Orto Byrosun river valley (Issyk I | x |
| KR179 | *P. armeniaca* | Wild | Issyk Kul lake surroundings - Orto Byrosun river valley (Issyk I | x |
| KR180 | *P. armeniaca* | Wild | Chuy river - Boom canyon (Chuy region, Kirgizstan) | x |
| KR181_1 | *P. armeniaca* | Wild | Chuy river - Boom canyon (Chuy region, Kirgizstan) | x |
| KR182 | *P. armeniaca* | Wild | Chuy river - Boom canyon (Chuy region, Kirgizstan) | x |
| KR183 | *P. armeniaca* | Wild | Chuy river - Boom canyon (Chuy region, Kirgizstan) | x |
| KR184 | *P. armeniaca* | Wild | Chuy river - Boom canyon (Chuy region, Kirgizstan) | x |
| KR185 | *P. armeniaca* | Wild | Chuy river - Boom canyon (Chuy region, Kirgizstan) | x |
| KR186 | *P. armeniaca* | Wild | Chuy river - Boom canyon (Chuy region, Kirgizstan) | x |
| KR187_2 | *P. armeniaca* | Wild | Chuy river - Boom canyon (Chuy region, Kirgizstan) | x |
| KR188 | *P. armeniaca* | Wild | Chuy river - Boom canyon (Chuy region, Kirgizstan) | x |
| KR189 | *P. armeniaca* | Wild | Chuy river - Boom canyon (Chuy region, Kirgizstan) | x |
| KR190 | *P. armeniaca* | Wild | Chuy river - Boom canyon (Chuy region, Kirgizstan) | x |
| KR191 | *P. armeniaca* | Wild | Chuy river - Boom canyon (Chuy region, Kirgizstan) | x |
| KR192 | *P. armeniaca* | Wild | Chuy river - Boom canyon (Chuy region, Kirgizstan) | x |
| KR193 | *P. armeniaca* | Wild | Chuy river - Boom canyon (Chuy region, Kirgizstan) | x |
| KR195 | *P. armeniaca* | Wild | Kyrgyz National Park Ala-Archa (Chuy region, Kirgizstan) | x |
| KR196 | *P. armeniaca* | Wild | Kyrgyz National Park Ala-Archa (Chuy region, Kirgizstan) | x |
| KR199 | *P. armeniaca* | Wild | Kyrgyz National Park Ala-Archa (Chuy region, Kirgizstan) | x |
| KR200 | *P. armeniaca* | Wild | Kyrgyz National Park Ala-Archa (Chuy region, Kirgizstan) | x |
| KR202 | *P. armeniaca* | Wild | Kyrgyz National Park Ala-Archa (Chuy region, Kirgizstan) | x |
| KR203 | *P. armeniaca* | Wild | Kyrgyz National Park Ala-Archa (Chuy region, Kirgizstan) | x |
| KR204 | *P. armeniaca* | Wild | Kyrgyz National Park Ala-Archa (Chuy region, Kirgizstan) | x |
| KR205 | *P. armeniaca* | Wild | Kyrgyz National Park Ala-Archa (Chuy region, Kirgizstan) | x |
| KR206 | *P. armeniaca* | Wild | Kyrgyz National Park Ala-Archa (Chuy region, Kirgizstan) | x |
| KR72_1 | *P. armeniaca* | Wild | Sary-Chelek Biosphere reserve (Jalal-Abad region, Kyrgyzstan) | x |
| KR73_1 | *P. armeniaca* | Wild | Sary-Chelek Biosphere reserve (Jalal-Abad region, Kyrgyzstan) | x |
| KR74 | *P. armeniaca* | Wild | Sary-Chelek Biosphere reserve (Jalal-Abad region, Kyrgyzstan) | x |
| KR75_1 | *P. armeniaca* | Wild | Sary-Chelek Biosphere reserve (Jalal-Abad region, Kyrgyzstan) | x |
| KR76_1 | *P. armeniaca* | Wild | Sary-Chelek Biosphere reserve (Jalal-Abad region, Kyrgyzstan) | x |
| KR77_1 | *P. armeniaca* | Wild | Sary-Chelek Biosphere reserve (Jalal-Abad region, Kyrgyzstan) | x |
| KR78_1 | *P. armeniaca* | Wild | Sary-Chelek Biosphere reserve (Jalal-Abad region, Kyrgyzstan) | x |
| KR79_1 | *P. armeniaca* | Wild | Sary-Chelek Biosphere reserve (Jalal-Abad region, Kyrgyzstan) | x |
| KR80_2 | *P. armeniaca* | Wild | Sary-Chelek Biosphere reserve (Jalal-Abad region, Kyrgyzstan) | x |
| KR81_1 | *P. armeniaca* | Wild | Sary-Chelek Biosphere reserve (Jalal-Abad region, Kyrgyzstan) | x |
| KR82_1 | *P. armeniaca* | Wild | Sary-Chelek Biosphere reserve (Jalal-Abad region, Kyrgyzstan) | x |
| KR83_1 | *P. armeniaca* | Wild | Sary-Chelek Biosphere reserve (Jalal-Abad region, Kyrgyzstan) | x |
| KR84_1 | *P. armeniaca* | Wild | Sary-Chelek Biosphere reserve (Jalal-Abad region, Kyrgyzstan) | x |
| KR85_2 | *P. armeniaca* | Wild | Sary-Chelek Biosphere reserve (Jalal-Abad region, Kyrgyzstan) | x |
| KR87_1 | *P. armeniaca* | Wild | Sary-Chelek Biosphere reserve (Jalal-Abad region, Kyrgyzstan) | x |
| KR88_1 | *P. armeniaca* | Wild | Sary-Chelek Biosphere reserve (Jalal-Abad region, Kyrgyzstan) | x |
| KR89_1 | *P. armeniaca* | Wild | Sary-Chelek Biosphere reserve (Jalal-Abad region, Kyrgyzstan) | x |
| KR90_1 | *P. armeniaca* | Wild | Sary-Chelek Biosphere reserve (Jalal-Abad region, Kyrgyzstan) | x |
| KR91_1 | *P. armeniaca* | Wild | Sary-Chelek Biosphere reserve (Jalal-Abad region, Kyrgyzstan) | x |
| KR92_1 | *P. armeniaca* | Wild | Sary-Chelek Biosphere reserve (Jalal-Abad region, Kyrgyzstan) | x |
| KR94 | *P. armeniaca* | Wild | Sary-Chelek Biosphere reserve (Jalal-Abad region, Kyrgyzstan) | x |
| KR95 | *P. armeniaca* | Wild | Sary-Chelek Biosphere reserve (Jalal-Abad region, Kyrgyzstan) | x |
| B15Rg1Lg16 | *P.mandshurica* | Wild | | x |
| CH_260 | *P.mandshurica* | Wild | Khanka Lake (Heilongjiang province, China) | x |
| CH_261 | *P.mandshurica* | Wild | Khanka Lake (Heilongjiang province, China) | x |

| | | | | |
|---|---|---|---|---|
| CH_263 | *P.mandshurica* | Wild | Khanka Lake (Heilongjiang province, China) | x |
| CH_264 | *P.mandshurica* | Wild | Khanka Lake (Heilongjiang province, China) | x |
| CH_265 | *P.mandshurica* | Wild | Khanka Lake (Heilongjiang province, China) | x |
| CH_270 | *P.mandshurica* | Wild | Khanka Lake (Heilongjiang province, China) | x |
| CH_273 | *P.mandshurica* | Wild | Khanka Lake (Heilongjiang province, China) | x |
| CH_304 | *P. sibirica* | Wild | Pingliang (Gansu province, China) | x |
| CH_305 | *P. sibirica* | Wild | Pingliang (Gansu province, China) | x |
| CH_307 | *P. sibirica* | Wild | Pingliang (Gansu province, China) | x |
| CH_308 | *P. sibirica* | Wild | Pingliang (Gansu province, China) | x |
| CH_309 | *P. sibirica* | Wild | Pingliang (Gansu province, China) | x |
| CH_311 | *P. sibirica* | Wild | Pingliang (Gansu province, China) | x |
| CH_312 | *P. sibirica* | Wild | Pingliang (Gansu province, China) | x |
| CH_313 | *P. sibirica* | Wild | Pingliang (Gansu province, China) | x |
| CH_314 | *P. sibirica* | Wild | Pingliang (Gansu province, China) | x |
| CH_316 | *P. sibirica* | Wild | Pingliang (Gansu province, China) | x |
| CH_317 | *P. sibirica* | Wild | Pingliang (Gansu province, China) | x |
| CH_318 | *P. sibirica* | Wild | Pingliang (Gansu province, China) | x |
| CH_319 | *P. sibirica* | Wild | Pingliang (Gansu province, China) | x |
| CH_320 | *P. sibirica* | Wild | Pingliang (Gansu province, China) | x |
| CH_289 | *P. sibirica* | Wild | Jingbian (Shaanxi province, China) | x |
| CH_290 | *P. sibirica* | Wild | Jingbian (Shaanxi province, China) | x |
| CH_291 | *P. sibirica* | Wild | Jingbian (Shaanxi province, China) | x |
| CH_292 | *P. sibirica* | Wild | Jingbian (Shaanxi province, China) | x |
| CH_293 | *P. sibirica* | Wild | Jingbian (Shaanxi province, China) | x |
| CH_274 | *P. sibirica* | Wild | Ansai (Shaanxi province, China) | x |
| CH_275 | *P. sibirica* | Wild | Ansai (Shaanxi province, China) | x |
| CH_276 | *P. sibirica* | Wild | Ansai (Shaanxi province, China) | x |
| CH_277 | *P. sibirica* | Wild | Ansai (Shaanxi province, China) | x |
| CH_278 | *P. sibirica* | Wild | Ansai (Shaanxi province, China) | x |
| CH_279 | *P. sibirica* | Wild | Ansai (Shaanxi province, China) | x |
| CH_280 | *P. sibirica* | Wild | Ansai (Shaanxi province, China) | x |
| CH_281 | *P. sibirica* | Wild | Ansai (Shaanxi province, China) | x |
| CH_282 | *P. sibirica* | Wild | Ansai (Shaanxi province, China) | x |
| CH_283 | *P. sibirica* | Wild | Ansai (Shaanxi province, China) | x |
| CH_284 | *P. sibirica* | Wild | Ansai (Shaanxi province, China) | x |
| CH_285 | *P. sibirica* | Wild | Ansai (Shaanxi province, China) | x |
| CH_286 | *P. sibirica* | Wild | Ansai (Shaanxi province, China) | x |
| CH_287 | *P. sibirica* | Wild | Ansai (Shaanxi province, China) | x |
| CH_288 | *P. sibirica* | Wild | Ansai (Shaanxi province, China) | x |
| CH_294 | *P. sibirica* | Wild | Yanggao (Shanxi province, China) | x |
| CH_295 | *P. sibirica* | Wild | Yanggao (Shanxi province, China) | x |
| CH_296 | *P. sibirica* | Wild | Yanggao (Shanxi province, China) | x |
| CH_297 | *P. sibirica* | Wild | Yanggao (Shanxi province, China) | x |
| CH_298 | *P. sibirica* | Wild | Yanggao (Shanxi province, China) | x |
| CH_299 | *P. sibirica* | Wild | Yanggao (Shanxi province, China) | x |
| CH_300 | *P. sibirica* | Wild | Yanggao (Shanxi province, China) | x |
| CH_301 | *P. sibirica* | Wild | Yanggao (Shanxi province, China) | x |
| CH_302 | *P. sibirica* | Wild | Yanggao (Shanxi province, China) | x |
| CH_303 | *P. sibirica* | Wild | Yanggao (Shanxi province, China) | x |
| CH_321 | *P. sibirica* | Wild | Weichang (Liaoning province, China) | x |
| CH_322 | *P. sibirica* | Wild | Weichang (Liaoning province, China) | x |
| CH_323 | *P. sibirica* | Wild | Weichang (Liaoning province, China) | x |
| CH_325 | *P. sibirica* | Wild | Weichang (Liaoning province, China) | x |
| CH_326 | *P. sibirica* | Wild | Weichang (Liaoning province, China) | x |
| CH_327 | *P. sibirica* | Wild | Weichang (Liaoning province, China) | x |
| CH_328 | *P. sibirica* | Wild | Weichang (Liaoning province, China) | x |
| CH_329 | *P. sibirica* | Wild | Weichang (Liaoning province, China) | x |
| CH_330 | *P. sibirica* | Wild | Weichang (Liaoning province, China) | x |
| CH_244 | *P. sibirica* | Wild | Chaoyang (Liaoning province, China) | x |
| CH_245 | *P. sibirica* | Wild | Chaoyang (Liaoning province, China) | x |
| CH_246 | *P. sibirica* | Wild | Chaoyang (Liaoning province, China) | x |
| CH_247 | *P. sibirica* | Wild | Chaoyang (Liaoning province, China) | x |
| CH_248 | *P. sibirica* | Wild | Chaoyang (Liaoning province, China) | x |
| CH_249 | *P. sibirica* | Wild | Chaoyang (Liaoning province, China) | x |
| CH_250 | *P. sibirica* | Wild | Chaoyang (Liaoning province, China) | x |
| CH_251 | *P. sibirica* | Wild | Chaoyang (Liaoning province, China) | x |
| CH_252 | *P. sibirica* | Wild | Chaoyang (Liaoning province, China) | x |
| CH_253 | *P. sibirica* | Wild | Chaoyang (Liaoning province, China) | x |
| CH_254 | *P. sibirica* | Wild | Chaoyang (Liaoning province, China) | x |
| CH_255 | *P. sibirica* | Wild | Chaoyang (Liaoning province, China) | x |

| | | | | | | |
|---|---|---|---|---|---|---|
| CH_256 | *P. sibirica* | Wild | Chaoyang (Liaoning province, China) | x | | |
| CH_257 | *P. sibirica* | Wild | Chaoyang (Liaoning province, China) | x | | |
| CH_258 | *P. sibirica* | Wild | Chaoyang (Liaoning province, China) | x | | |
| CH_259 | *P. sibirica* | Wild | Chaoyang (Liaoning province, China) | x | | |
| CH_229 | *P. sibirica* | Wild | Fuxin (Liaoning province, China) | x | | |
| CH_230 | *P. sibirica* | Wild | Fuxin (Liaoning province, China) | x | | |
| CH_231 | *P. sibirica* | Wild | Fuxin (Liaoning province, China) | x | | |
| CH_232 | *P. sibirica* | Wild | Fuxin (Liaoning province, China) | x | | |
| CH_233 | *P. sibirica* | Wild | Fuxin (Liaoning province, China) | x | | |
| CH_234 | *P. sibirica* | Wild | Fuxin (Liaoning province, China) | x | | |
| CH_235 | *P. sibirica* | Wild | Fuxin (Liaoning province, China) | x | | |
| CH_236 | *P. sibirica* | Wild | Fuxin (Liaoning province, China) | x | | |
| CH_237 | *P. sibirica* | Wild | Fuxin (Liaoning province, China) | x | | |
| CH_238 | *P. sibirica* | Wild | Fuxin (Liaoning province, China) | x | | |
| CH_239 | *P. sibirica* | Wild | Fuxin (Liaoning province, China) | x | | |
| CH_240 | *P. sibirica* | Wild | Fuxin (Liaoning province, China) | x | | |
| CH_241 | *P. sibirica* | Wild | Fuxin (Liaoning province, China) | x | | |
| CH_242 | *P. sibirica* | Wild | Fuxin (Liaoning province, China) | x | | |
| CH_243 | *P. sibirica* | Wild | Fuxin (Liaoning province, China) | x | | |
| A2779 | *P.mume* | Cultivated | | x | x | |
| CH_225 | *P.mume* | Cultivated | Jiangsu province, China | x | x | |
| CH_226 | *P.mume* | Cultivated | Jiangsu province, China | x | x | |
| CH_227 | *P.mume* | Cultivated | Jiangsu province, China | x | x | |
| CH_228 | *P.mume* | Cultivated | Jiangsu province, China | x | x | |
| US242 | *P.mume* | Cultivated | Japan | x | x | |
| US243 | *P.mume* | Cultivated | Japan | x | x | |
| US244 | *P.mume* | Cultivated | Japan | x | x | |
| US245 | *P.mume* | Cultivated | Taiwan | x | x | |
| ch333 | *P. salicina* | Wild | China | | x | |
| ch334 | *P. salicina* | Wild | China | | x | |
| ch335 | *P. salicina* | Wild | China | | x | |
| ch336 | *P. salicina* | Wild | China | | x | |
| ch337 | *P. salicina* | Wild | China | | x | |
| US91 | *P. salicina* | Cultivated | | | x | |
| US13 | *P. salicina* | Cultivated | | | x | |
| US63 | *P. salicina* | Cultivated | | | x | |
| KR233 | *P. salicina* | Cultivated | | | x | x |
| P0489 | *P. salicina* | Cultivated | | | x | x |
| FR-070 | *P. cerasifera* | Wild | Alps, France | | x | x |
| X29 | *P. cerasifera* | Wild | | | x | x |
| az117 | *P. cerasifera* | Wild | | | x | x |
| az118 | *P. cerasifera* | Wild | | | x | x |
| az119 | *P. cerasifera* | Wild | | | x | x |
| az120 | *P. cerasifera* | Wild | | | x | x |
| az121 | *P. cerasifera* | Wild | | | x | x |
| az122 | *P. cerasifera* | Wild | | | x | x |
| az123 | *P. cerasifera* | Wild | | | x | x |
| az124 | *P. cerasifera* | Wild | | | x | x |
| az125 | *P. cerasifera* | Wild | | | x | x |
| az126 | *P. cerasifera* | Wild | | | x | x |
| az127 | *P. cerasifera* | Wild | | | x | x |
| az128 | *P. cerasifera* | Wild | | | x | x |
| az129 | *P. cerasifera* | Wild | | | x | x |
| az130 | *P. cerasifera* | Wild | | | x | x |
| az131 | *P. cerasifera* | Wild | | | x | x |
| az192 | *P. cerasifera* | Wild | | | x | x |
| az193 | *P. cerasifera* | Wild | | | x | x |
| az197 | *P. cerasifera* | Wild | | | x | x |
| az239 | *P. cerasifera* | Wild | | | x | x |
| az24 | *P. cerasifera* | Wild | | | x | x |
| az25 | *P. cerasifera* | Wild | | | x | x |
| az48 | *P. cerasifera* | Wild | | | x | x |
| az49 | *P. cerasifera* | Wild | | | x | x |
| az51 | *P. cerasifera* | Wild | | | x | x |
| az59 | *P. cerasifera* | Wild | | | x | x |
| KR208 | *P. cerasifera* | Wild | | | x | x |
| KR209 | *P. cerasifera* | Wild | | | x | x |
| KR210 | *P. cerasifera* | Wild | | | x | x |
| KR211 | *P. cerasifera* | Wild | | | x | x |
| KR215 | *P. cerasifera* | Wild | | | x | x |

| | | | | |
|---|---|---|---|---|
| KR216 | *P. cerasifera* | Wild | x | x |
| KR219 | *P. cerasifera* | Wild | x | x |
| KR225 | *P. cerasifera* | Wild | x | x |
| KR226 | *P. cerasifera* | Wild | x | x |
| KR227 | *P. cerasifera* | Wild | x | x |
| KR228 | *P. cerasifera* | Wild | x | x |
| KR230 | *P. cerasifera* | Wild | x | x |
| KR231 | *P. cerasifera* | Wild | x | x |
| KR232 | *P. cerasifera* | Wild | x | x |
| KR234 | *P. cerasifera* | Wild | x | x |
| KR244 | *P. cerasifera* | Wild | x | x |
| KR245 | *P. cerasifera* | Wild | x | x |
| KR255 | *P. cerasifera* | Wild | x | x |
| KR256 | *P. cerasifera* | Wild | x | x |
| kz281 | *P. cerasifera* | Wild | x | x |
| kz282 | *P. cerasifera* | Wild | x | x |
| kz284 | *P. cerasifera* | Wild | x | x |
| kz298 | *P. cerasifera* | Wild | x | x |
| kz303 | *P. cerasifera* | Wild | x | x |
| P016-5 | *P. cerasifera* | - | x | x |
| P0018 | *P. cerasifera* | - | x | x |
| P1079 | *P. cerasifera* | - | x | x |
| P2126 | *P. cerasifera* | - | x | x |
| P2175 | *P. cerasifera* | - | x | x |
| P2196 | *P. cerasifera* | - | x | x |
| P2214 | *P. cerasifera* | - | x | x |
| P2415 | *P. cerasifera* | - | x | x |
| P2550 | *P. cerasifera* | - | x | x |
| P2646 | *P. cerasifera* | - | x | x |
| P3103 | *P. cerasifera* | - | x | x |
| P3188 | *P. cerasifera* | - | x | x |
| P3195 | *P. cerasifera* | - | x | x |
| P3196 | *P. cerasifera* | - | x | x |
| P3209 | *P. cerasifera* | - | x | x |
| P3289 | *P. cerasifera* | - | x | x |
| P3676 | *P. subcordata* | - | x | |
| US14 | *P. mexicana* | - | x | |
| US3 | *P. musoniana* | - | x | |
| US64 | *P. maritima* | - | x | |
| US88 | *P. americana* | - | x | |

**Table S2. Analysis of genetic variability from microsatellites for *P. brigantina* population.**

| Locus | % missing | *Na* | *Ne* | *I* | *Ho* | *He* |
|---|---|---|---|---|---|---|
| AMPA100 | 7.04 | 1 | 1.00 | 0.00 | 0.00 | 0.00 |
| AMPA101 | 1.41 | 5 | 2.57 | 1.09 | 0.21 | 0.61 |
| AMPA103 | 1.41 | 8 | 3.67 | 1.50 | 0.56 | 0.73 |
| aprigms18 | 4.23 | 5 | 2.34 | 1.00 | 0.49 | 0.57 |
| BPPCT004 | 4.23 | 2 | 1.14 | 0.24 | 0.10 | 0.12 |
| BPPCT025 | 2.82 | 3 | 1.52 | 0.64 | 0.25 | 0.34 |
| BPPCT040 | 5.63 | 8 | 5.25 | 1.75 | 0.60 | 0.81 |
| CPPCT030 | 1.41 | 1 | 1.00 | 0.00 | 0.00 | 0.00 |
| CPPCT006 | 0.00 | 6 | 3.40 | 1.32 | 0.49 | 0.71 |
| CPPCT033 | 1.41 | 11 | 3.94 | 1.78 | 0.69 | 0.75 |
| EPPCU0532 | 4.23 | 1 | 1.00 | 0.00 | 0.00 | 0.00 |
| G22SSR | 1.41 | 2 | 1.01 | 0.04 | 0.01 | 0.01 |
| SSRLg1_11m52a | 0.00 | 13 | 4.50 | 1.89 | 0.66 | 0.78 |
| N86B11ssr3 | 2.82 | 3 | 1.92 | 0.83 | 0.43 | 0.48 |
| pchgms03 | 2.82 | 1 | 1.00 | 0.00 | 0.00 | 0.00 |
| PGS-1.21 | 1.41 | 3 | 1.24 | 0.37 | 0.19 | 0.19 |
| SSR04AG51 | 0.00 | 9 | 4.06 | 1.72 | 0.55 | 0.75 |
| SSR5piso4E | 2.82 | 1 | 1.00 | 0.00 | 0.00 | 0.00 |
| SSR5piso4Ga | 1.41 | 3 | 2.35 | 0.94 | 0.37 | 0.57 |
| UDA-021 | 2.82 | 2 | 1.33 | 0.41 | 0.17 | 0.25 |
| UDAp-414 | 0.00 | 8 | 5.20 | 1.79 | 0.48 | 0.81 |
| UDAp-480 | 2.82 | 8 | 2.61 | 1.25 | 0.43 | 0.62 |
| UDP98-409 | 2.82 | 11 | 4.28 | 1.77 | 0.58 | 0.77 |
| UDP96-018 | 4.23 | 6 | 2.19 | 0.95 | 0.31 | 0.54 |
| Mean | **2.46** | **5.042** | **2.48** | **0.89** | **0.32** | **0.43** |

Footnote: *Na* : number of different alleles, and *Ne* : number of

### Table S3. The description of individuals retained for the core collection of *Prunus brigantina*

| **Percentage of genetic diversity of the core collection** | 100% | 90% | 80% | 70% | 60% | 50% | 40% | 30% | <= 20% |
|---|---|---|---|---|---|---|---|---|---|
| | FR_003 | FR_003 | FR_010 | FR_025 | FR_003 | FR_032 | FR_003 | FR_003 | FR_039_B |
| | FR_009 | FR_010 | FR_014 | FR_030_1 | FR_037 | FR_036 | FR_037 | FR_058 | |
| | FR_010 | FR_014 | FR_032 | FR_051 | FR_043 | FR_051 | FR_057 | | |
| | FR_012 | FR_029 | FR_037 | FR_052 | FR_052 | FR_065 | | | |
| | FR_014 | FR_032 | FR_043 | FR_057 | FR_059 | | | | |
| | FR_015 | FR_037 | FR_045 | FR_063 | FR_065 | | | | |
| | FR_019 | FR_043 | FR_051 | FR_065 | | | | | |
| | FR_023 | FR_045 | FR_052 | FR_066 | | | | | |
| | FR_025 | FR_050 | FR_057 | | | | | | |
| | FR_028 | FR_051 | FR_063 | | | | | | |
| | FR_029 | FR_052 | FR_065 | | | | | | |
| | FR_030_1 | FR_057 | FR_066 | | | | | | |
| | FR_032 | FR_059 | | | | | | | |
| | FR_036 | FR_063 | | | | | | | |
| | FR_037 | FR_065 | | | | | | | |
| | FR_039_A | FR_066 | | | | | | | |
| | FR_043 | FR_067 | | | | | | | |
| **Individuals in the core** | FR_045 | FR_068 | | | | | | | |
| **collection** | FR_046 | | | | | | | | |
| | FR_047 | | | | | | | | |
| | FR_048 | | | | | | | | |
| | FR_049 | | | | | | | | |
| | FR_050 | | | | | | | | |
| | FR_051 | | | | | | | | |
| | FR_052 | | | | | | | | |
| | FR_053 | | | | | | | | |
| | FR_057 | | | | | | | | |
| | FR_058 | | | | | | | | |
| | FR_059 | | | | | | | | |
| | FR_061 | | | | | | | | |
| | FR_063 | | | | | | | | |
| | FR_064 | | | | | | | | |
| | FR_065 | | | | | | | | |
| | FR_066 | | | | | | | | |
| | FR_067 | | | | | | | | |
| | FR_068 | | | | | | | | |
| **Core collection size** | 36 | 18 | 12 | 8 | 6 | 4 | 3 | 2 | 1 |

**Table S4. Genetic variability from microsatellites for *P. brigantina* core collection.**

| Locus | % miss | *Na* | *Ne* | *I* | *Ho* | *He* |
|---|---|---|---|---|---|---|
| AMPA100 | 3.1 | 1 | 1.00 | 0.00 | 0.00 | 0.00 |
| AMPA101 | 0.0 | 5 | 2.83 | 1.21 | 0.31 | 0.65 |
| ampa103 | 0.0 | 8 | 3.39 | 1.51 | 0.47 | 0.71 |
| aprigms18 | 3.1 | 5 | 2.45 | 1.07 | 0.55 | 0.59 |
| BPPCT004 | 6.3 | 2 | 1.14 | 0.25 | 0.07 | 0.12 |
| BPPCT025 | 0.0 | 3 | 1.47 | 0.60 | 0.13 | 0.32 |
| BPPCT040 | 6.3 | 8 | 5.83 | 1.84 | 0.67 | 0.83 |
| CCPCT030 | 0.0 | 1 | 1.00 | 0.00 | 0.00 | 0.00 |
| CPPCT006 | 0.0 | 6 | 3.75 | 1.43 | 0.53 | 0.73 |
| CPPCT033 | 0.0 | 11 | 5.69 | 2.03 | 0.75 | 0.82 |
| eppco0532 | 0.0 | 1 | 1.00 | 0.00 | 0.00 | 0.00 |
| g22ssr | 3.1 | 2 | 1.03 | 0.08 | 0.03 | 0.03 |
| L11m52a | 0.0 | 13 | 5.36 | 2.10 | 0.69 | 0.81 |
| N86B11SSR3 | 0.0 | 3 | 1.96 | 0.84 | 0.41 | 0.49 |
| pchms03 | 0.0 | 1 | 1.00 | 0.00 | 0.00 | 0.00 |
| PGS-21a | 3.1 | 3 | 1.34 | 0.47 | 0.23 | 0.25 |
| ssr04ag51 | 0.0 | 9 | 4.46 | 1.78 | 0.56 | 0.78 |
| ssr5piso4e | 3.1 | 1 | 1.00 | 0.00 | 0.00 | 0.00 |
| ssr5piso4ga | 0.0 | 3 | 2.00 | 0.80 | 0.22 | 0.50 |
| uda021 | 0.0 | 2 | 1.44 | 0.48 | 0.25 | 0.31 |
| udap-414 | 0.0 | 8 | 4.84 | 1.75 | 0.41 | 0.79 |
| udap-480 | 0.0 | 8 | 3.14 | 1.46 | 0.38 | 0.68 |
| UDP409 | 6.3 | 11 | 6.48 | 2.06 | 0.70 | 0.85 |
| udp96-018 | 3.1 | 6 | 2.30 | 1.07 | 0.39 | 0.57 |
| Mean | **1.6** | **5.042** | **2.75** | **0.95** | **0.32** | **0.45** |

Footnote: *Na* : number of different alleles, and *Ne* : number

**Table S5. Mann-Whitney U Test (two-tailed) between the whole dataset *P. brigantina* population and its core collection.**

| Mann-Whitney U Test (two-tailed) | *I* | *Ho* | *He* | µ*He* |
|---|---|---|---|---|
| *Z-value* | 0.330 | -0.052 | -0.340 | -0.412 |
| *p-value* | 0.741 | 0.960 | 0.728 | 0.682 |

Footnotes: significant at 0.05 level
